# Structure and dynamics of a muti-domain nitric oxide synthase regulated by a C2 domain

**DOI:** 10.1101/2025.08.15.670619

**Authors:** Dhruva Nair, Brian R. Crane

## Abstract

Nitric oxide synthase (NOS) is a widely studied multidomain redox enzyme that produces the key signaling molecule and cytotoxic agent nitric oxide (NO) for functions that range from mammalian vasodilation to prokaryotic antibiotic resistance. NOS enzymes from metazoans and cyanobacteria rely on dynamic associations of their oxygenase and coupled di-flavin reductase domains that have largely evaded detailed structural characterization. CryoEM studies of a representative dimeric six-domain Synechococcus NOS reveal the architecture of the full-length enzyme, which contains an unusual regulatory C2 domain, and additional nitric oxide deoxygenase (NOD) and pseudo-globin modules. Five distinct structural states depict how pterin binding couples to tight and loose oxygenase conformations and how the Ca^2+^-sensitive C2 domain moves over 85 Å to alternatively regulate either the NOS or NOD heme center. The extended C-terminal tail and its dynamic interactions highlight an added layer of regulation required by multidomain NOSs compared to other di-flavin reductases.

**Teaser:** syNOS is a highly dynamic multi-domain oxidoreductase that harnesses a Ca^2+^-sensitive C2 domain to modulate activity.

## Introduction

Nitric oxide (NO) is a potent free-radical signal and cytotoxic agent found throughout biology (*1–8*). Nitric Oxide Synthases (NOS) are homodimeric P450-like enzymes that produce NO through the 5-electron oxidation of L-arginine (L-arg) to L-citrulline (*9–12*). NOSs comprise a heme-containing oxygenase domain (NOS_Oxy_) that is reductively activated by either an appended reductase domain (NOS_Red_, characteristic of metazoans) or an auxiliary reductase partner (characteristic of prokaryotes) (Fig 1A) (*10–12*).

**Figure 1.**
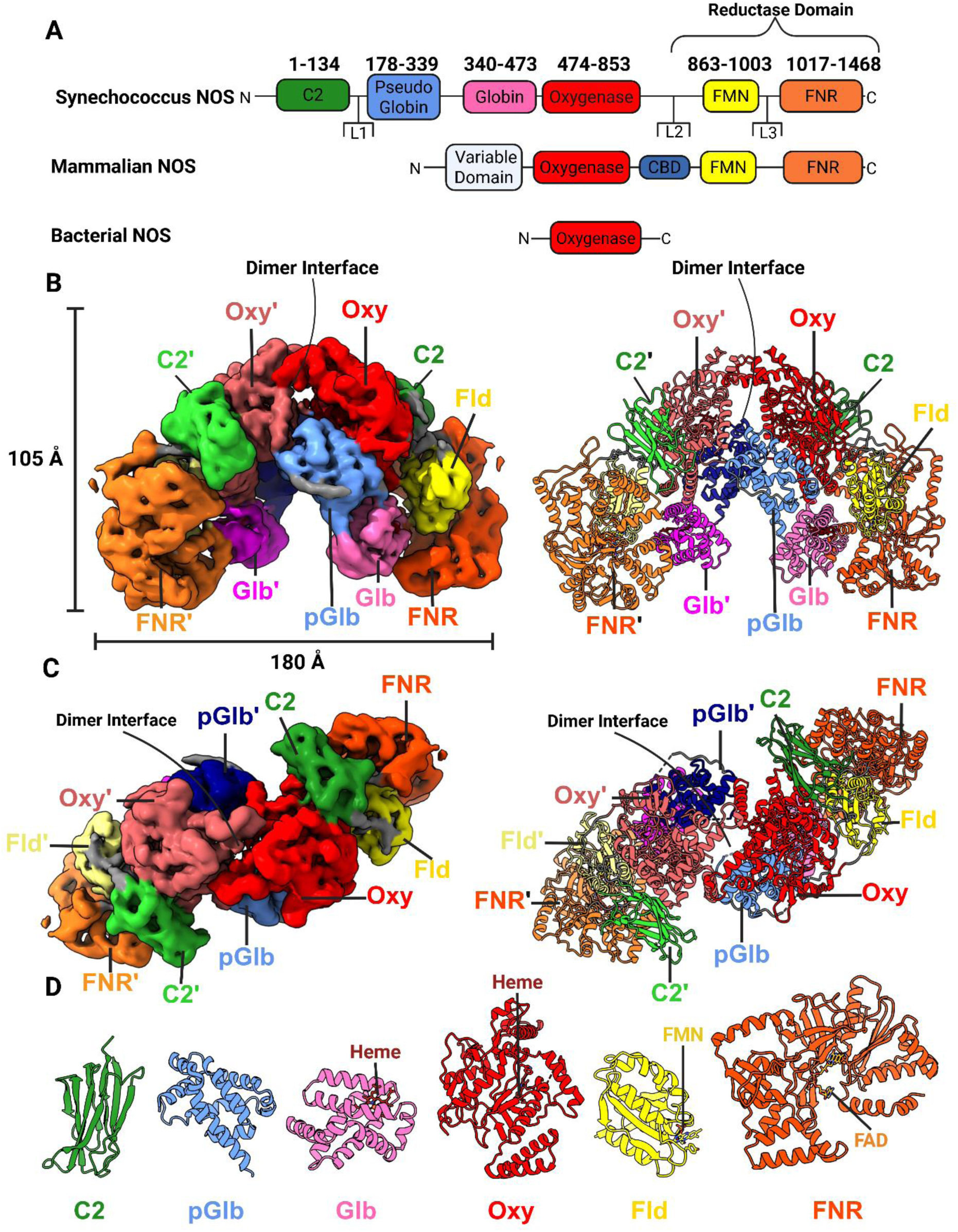
Architecture of Ca^2+^-free locked syNOS. (A) Gene architecture of Synechococcus NOS, mammalian NOS and a typical bacterial NOS. Note that some non-cyanobacterial prokaryotic NOSs have auxiliary domains. (B) Sideview of electron density (left, threshold = 0.0298, C2 symmetry applied) and corresponding ribbon representation (right) for the locked syNOS homodimer (NOS-Sym). Subunit 1 (right) colored as designated by the gene architecture, Subunit 2 (left) colored as follows: C2’ – lime green, pGlb’ – navy, Glb’ – magenta, Oxy’ – indian red, Fld’ – pale yellow and FNR’ – bright orange. (C) Top view of (B). (D) Individual syNOS domains with cofactors indicated.

Mammalian NOSs (mNOSs) are multidomain enzymes composed of an N-terminal NOS_Oxy_ and a C-terminal NADPH-dependent NOS_Red_, bridged by a calmodulin (CAM)-binding domain (NOS_CBD_; Fig. 1A). NOS_Red_ belongs to the cytochrome P450 reductase (CYPOR) family and as such, is further divided into a NOS_Oxy_-proximal FMN-binding flavodoxin domain (NOS_Fld_) and an FAD-and NADPH-binding ferredoxin-NADP^+^ reductase module (NOS_FNR_) (*10–13*). NOS_Oxy_ utilizes tetrahydrobiopterin (BH_4_) or tetrahydrofolate for rapid heme reduction during oxygen activation (*14*). To reduce heme for oxygen binding, electrons flow from NADPH –> FAD –> FMN(H)• –> Heme-Fe^3+^. Ca^2+^-CAM enhances heme reduction by facilitating the traverse of NOS_Fld_ ∼70 Å from a shielded position in a conformationally locked state of NOS_Red_ to a productive binding interaction with NOS_Oxy_ (*15–18*).

Given their medicinal importance (*19*), NOSs have been extensively studied in mammals, which have three isoforms: 1) endothelial NOS (eNOS, regulation of the vasculature), 2) neuronal NOS (nNOS, regulation of neuronal function), and 3) inducible NOS (iNOS, immune responses to pathogens and cancer cells) (*11*, *20–22*). Most prokaryotic NOSs have fewer domains than their metazoan counterparts and have diverse functions that include participation in stress responses, antibiotic resistance and biosynthesis of secondary metabolites (*23–27*).

To mitigate NO cytotoxicity, prokaryotes oxidize NO to NO_3_^-^ with flavohemoprotein (Hmp), which combines a heme-reducing flavin reductase with an oxygen-activating globin domain. In contrast, mammals oxidize NO with hemoglobin, cytoglobin and neuroglobin, which obtain reduced equivalents from various environmental factors (*28–30*).

Cyanobacterial NOSs represent a bridge between metazoan and prokaryotic NOSs (*26*, *31–34*). They comprise an mNOS architecture fused to an N-terminal globin domain, as exemplified by the 1468-residue NOS from *Synechococcus sp*. PCC 7335 (i.e. syNOS, Fig. 1A). Like mNOS, Ca^2+^ activates syNOS, despite syNOS lacking a CBD and Synechococcus lacking an obvious CAM homolog (*34*). The syNOS globin domain (NOS_Glb_) oxidizes NO to NO_3_ ^-^ as a nitric oxide dioxygenase (NOD) and, when considered with syNOS_FNR_, resembles Hmp (*34*).

Ca^2+^-CAM regulates mNOS activity by controlling the motions and associations of NOS_Oxy_ and NOS_Red_ that allow electron transfer (ET) between them. Owing to the dynamic nature of turnover and inactivated states of full-length NOSs, they have evaded characterization by direct structural methods. Single particle electron microscopy (EM) in conjunction with chemical crosslinking and hydrogen-deuterium exchange mass spectrometry (HDX-MS) have yielded informative, if incomplete, structural information (*35–39*). These studies, along with biochemical and spectroscopic characterization (*12*, *40–44*) have motivated a model of mNOS conformational dynamics that involves three states: i) input, wherein NOS_Fld_ closely associates with NOS_FNR_ so that the NOS_Red_ FAD can be reduced by NADPH; ii) intermediate, wherein NOS_Fld_ transitions between its closed position next to NOS_FNR_ and an interaction with NOS_Oxy_; and iii) output, wherein Ca^2+^-CAM promotes docking of NOS_Fld_ onto the NOS_Oxy_ of the adjacent subunit (*44–47*). Although cryoEM has provided residue-level information on NOS_Oxy_, NOS_Red_ positioning has been challenging to discern owing to its conformational heterogeneity (*35*, *46*).

Herein, we resolve domain configurations of the multi-domain syNOS by cryoEM in conjunction with further characterizing ET properties associated with its NOS and NOD activities. We capture resting and dynamic states of the enzyme and identify an atypical C2-like domain that acts as a reversible lynchpin to mediate Ca^2+^ sensitivity. An elaborated C-terminal tail (CTT) (*22*) of NOS_Red_, key for regulation of mNOS, maintains a conformationally locked dimer and represents an expansion of the regulatory mechanisms seen within the di-flavin reductase family.

## Results and Discussion

### SyNOS expression, purification and crosslinking mass spectrometry

Full-length *Synechococcus PCC 7335* NOS (syNOS) was fused to an N-terminal-twin strep affinity tag and co-expressed along with *E. coli* HSP90 and GAPDH, in BL21(DE3) and purified through a series of affinity and size exclusion columns (SEC) (see Methods, Fig. S1) (*46*, *48*, *49*). As found previously (*34*), syNOS purified as a mixture of monomers and dimers. SEC fractions were tested for NO production with L-arg and NADPH and active fractions were applied to glow-discharged Cu-Quantifoil grids, vitrified and imaged on either a Talos Arctica or Titan Krios. Prior to freezing, samples were incubated with or without Ca^2+^/L-arg/NADPH and excess BH_4_ (Table S1).

To complement cryoEM and gain further insight into domain dynamics we subjected purified syNOS dimer fractions to disuccinimidyl dibutyric urea (DSBU) crosslinking mass spectrometry (XL-MS), under inactive (Ca^2+^-depleted) and turnover conditions (see methods for assay conditions). Peptide coverage was relatively high for both (inactive state, 91%; and turnover state, 93%) but only a limited number of intra/intersubunit Lys-to-Lys crosslinks were identified (Table S2).

### Structure determination of inactivated full-length syNOS

SyNOS data sets and the states within are summarized in Table S1 and described below. Two datasets of syNOS samples lacking Ca^2+^, L-arg and NADPH were collected on a Talos Arctica equipped with a K3 detector. Dataset (DS) 1 (1215 micrographs) and DS2 (4055 micrographs) were initially preprocessed using CryoSPARC (Fig. S2 and S3), then subjected to particle picking using CRYOLO and finally refined in CryoSPARC (*50*, *51*). During 2D classification multiple classes emerged resembling a symmetric full-length dimer (syNOS-Sym). DS1 produced a syNOS-Sym overall resolution of 3.8 Å (47 K particles), and DS2 produced a NOS-Sym map of 4.7 Å resolution (84 K particles).

In addition, an asymmetric class (syNOS-Asym) became apparent in both DS1 and DS2 that lacked density for one syNOS_Red_ and one syNOS_C2_. DS1 produced a syNOS-Asym map with the nominal resolution of 3.67 Å (67 K particles), whereas DS2 yielded a 4.00 Å resolution map (265 K particles). In syNOS-Asym, uninterpretable electron density surrounds the position of the vacated syNOS_Red_ indicative of conformational disorder for this subunit.

3D-refinement of syNOS-Sym gave clear electron density for secondary structure, cofactors and in many cases, residue side chains. Portions of each subunit were less well defined yet still allowed unambiguous placement of the constituent domains. Inspection of the electron density indicated that the homodimer interface was structurally heterogeneous. To better reconstruct the full-length dimer, we combined the syNOS-Sym particle stacks from DS1 (47 K particles) and DS2 (84 K particles) and re-refined the density alignment for 3D-variability (VA) and 3D-Flex analysis. We explored C1, C2 and C2 (relaxed) symmetries which yielded 3.99 Å, 3.95 Å and 3.95 Å nominal resolutions, respectively, with the C2(relaxed) refinement chosen for further flexibility analysis (Fig. S4, discussed below). The C2 symmetry-imposed electron density was sufficient for model building and refinement (Table S3).

### Structure determination of turnover syNOS with Ca^2+^ and L-arginine

DS3 (Fig. S5) (5019 micrographs) represented syNOS incubated with Ca^2+^, additional BH_4_ and L-arg and was collected on a Titan Krios equipped with a K3 detector. Micrographs were initially preprocessed using CryoSPARC (Fig. S5), then subjected to particle picking with CRYOLO and finally refined in CryoSPARC. During 2D classification, multiple classes emerged resembling NOS-Asym but also a state that lacked density for syNOS_Red_ on both subunits. The latter state resembles previously determined nNOS cryoEM structures that resolved a NOS_Oxy_ homodimer but lacked density for NOS_Red_ owing to conformational disorder (*36*). We refer to this class as the a “loose” turnover state (NOS-TO_L_), because the syNOS_Oxy_ interface is expanded relative to a typical “tight” mNOS_Oxy_ dimer. 2D classes of NOS-TO_L_ gave density plumes at the interface characteristic of poor signal alignment, indicative of a conformationally heterogeneous particle, hence a composite map from two local refinements was created(PDB:9Q15). syNOS-Asym (PDB:9Q0Y) was refined to a nominal resolution of 3.19 Å (71 K particles), whereas syNOS-TO_L_ (PDB:9Q0X) was resolved to 3.6 Å resolution (61 K particles, Table S3).

DS4 was collected from the same conditions on the 200 KV Talos Arctica equipped with a K3 detector. These grids, which had thicker cubic ice, produced an intermediate resolution turnover state with a tight syNOS_Oxy_ dimer interface (syNOS-TO_T_) that more closely resembles crystal structures of BH_4_-bound mNOS_Oxy_ (Fig. S6). SPA is possible from cubic ice conditions, which is known to favor contraction (shrinkage) of protein structure, although the mechanisms for this behavior are largely unknown (*52*).

### Structure determination of monomeric syNOS in turnover conditions

DS5 (9036 micrographs) of syNOS incubated with additional BH_4_, Ca^2+^, L-arg and NADPH was collected on a Titan Krios equipped with a Falcon 3 detector (Fig. S7). For this dataset, the homodimers appeared primarily dissociated into monomers (syNOS-Mon). Unlike syNOS-Asym, in syNOS-Mon, even syNOS_Oxy_ appeared as a monomer. The micrographs were processed through CryoSPARC with multiple rounds of 3D particle stack cleaning to remove damaged particles (Fig. S7). The syNOS-Mon (PDB:9Q05) density was refined to 3.09 Å resolution (45 K particles), with discernible sidechains and cofactors, although neither Ca^2+^, nor L-arg could be identified in the map (Table S3).

### Full-length syNOS (syNOS_FL_) molecular architecture

In the absence of Ca^2+^, the syNOS-Sym(PDB:9Q06) state of the full-length enzyme (syNOS_FL_) forms a compact C2-symmetric homodimer composed of six domains per subunit (Fig. 1B,C,D); the domain arrangements divide the protein into two redox modules: i) an Fld-FNR module with an N-terminal C2 domain (residues 1-134) bound at the Oxy-FNR interface and ii) a Glb-Oxy module, that also contains an 8-helical pseudo-globin domain that does not bind heme (syNOS_pGlb_, residues 179-339). Three extended linkers connect the most mobile domains, with L1 (44 residues) connecting syNOS_C2_ to syNOS_pGlb_, L2 (10 residues) connecting syNOS_Oxy_ to syNOS_Fld_ and L3 (13 residues) connecting syNOS_Fld_ to syNOS_FNR_ (Fig. S8).

In this symmetric “locked” state, syNOS dimerizes through interactions of syNOS_Oxy_ (residues 474-852), and syNOS_pGlb_, which bridges the gap between syNOS_Glb_ (residues 340-473) and syNOS_Oxy_ (Fig. 1B, C). SyNOS does not contain a β-hairpin region and tetrahedral zinc-binding site formed at the mNOS_Oxy_ dimer interface; instead syNOS_pGlb_ occupies this position, although an insertion between αD/ αE (residues 245-263), that would superimpose on the mNOS zinc-binding site is not well defined by the electron density (Fig. S9 A, B, C). The syNOS_Oxy_ C-terminal helical hairpin acts as an interaction hub at the center of the subunit by contacting syNOS_Fld_, syNOS_FNR_, syNOS_Glb_ and the syNOS_C2_ domain.

3D variable analysis (3DVA) and 3D Flexible refinement (3DFlex) conducted on syNOS-Sym (PDB:9Q06), revealed that syNOS subunits “breathe” between tight, ordered associations that bind BH_4_ within the helical lariats and more separated looser associations with lower density for these regions (Movie S1), not unlike the “tight” and “loose” dimer states previously observed for *B. subtilis* NOS (*53–55*). Most structures, except for NOS-TO_T_, lacked density for the helical lariats and BH_4_, which compose the center of the syNOS_Oxy_ dimer interface (Fig. S9). XL-MS indicated that syNOS in turnover conditions produced three additional interfacial crosslinks between syNOS_pGlb_ and syNOS_Oxy_ compared to the inactive protien (Fig S10 A – B). Thus, the active state favors a tighter dimer interface, consistent with BH_4_ binding. Samples collected in thicker ice favored this tight association (syNOS-TO_T_). In addition to an ordered NOS_Oxy_ interface and density for BH_4_, NOS-TO_T_ displayed new contacts between L1 and αA (syNOS_pGlb_) on one subunit and α9 (syNOS_Oxy_) on the opposing subunit. Hence, L1 may influence syNOS_pGlb_ positioning, which in turn propagates structural stability to the dimer interface (Fig. S9B).

The previously unannotated and uncharacterized N-terminal domain (residues 1-134) takes the form of a tightly packed 8-stranded β-sandwich, capped by peripheral loops at each end (Fig. 1C, D). This structure strongly resembles a C2 domain (syNOS_C2_), which is usually associated with Ca^2+^ –dependent membrane targeting in eukaryotes and typically not found in prokaryotes. (*56*, *57*) SyNOS_C2_ brackets the backside of the syNOS_Oxy_ heme pocket and syNOS_FNR_ (residues 1017-1468; Fig. 1C). In NOS-Sym, syNOS_FNR_ latches onto syNOS_Glb_ through the ordered CTT, which also binds to syNOS_Fld_ (residues 863-1003; Fig. 1B, C).

SyNOS_Fld_ and syNOS_FNR_ comprise a canonical NOS reductase unit in a closed conformation with the FMN-FAD isoalloxazine cofactors aligned in a planar end-to-end configuration (Fig. 2). To facilitate ET between subunits, mNOSs rearrange to position the NOS_Fld_ domain < 15 Å from the NOS_Oxy_ heme on the opposing subunit (*58*). Residue conservation, cross-linking and HDX-MS indicate that NOS_Fld_ interacts dynamically on the backside of the heme pocket at a conserved Trp residue (356 in human eNOS) (*36–38*, *55*). SyNOS_C2_ blocks this position and would have to move to allow productive ET from syNOS_Fld_, which also must displace from its shielded interaction with syNOS_FNR_ (*38*). Notably, syNOS lacks the mNOS CBD (∼30 residues) that upon CAM binding favors the ET competent ‘output state’ (Fig. 2A-C) (*17*, *38*, *59*, *60*). However, the syNOS interdomain linkers also likely mediate domain motions. These linkers generally have extended conformations, with the longest, L1, traversing ∼60 Å. The threading of L1 beneath L2 implies that any C2 movement will displace L2 and thereby cause syNOS_Fld_ to move “up-and-over” toward the adjacent syNOS_Oxy_ subunit (Fig. S8). In comparison to the interdomain interactions implied by the 25 Å resolution nNOS + CAM structure by negative-stain EM (*37*), syNOS_Fld_ likely undergoes a different trajectory to interact with syNOS_Oxy_ in part because syNOS_Glb_ will block the CAM-directed conformational change proposed for nNOS.

**Figure 2.**
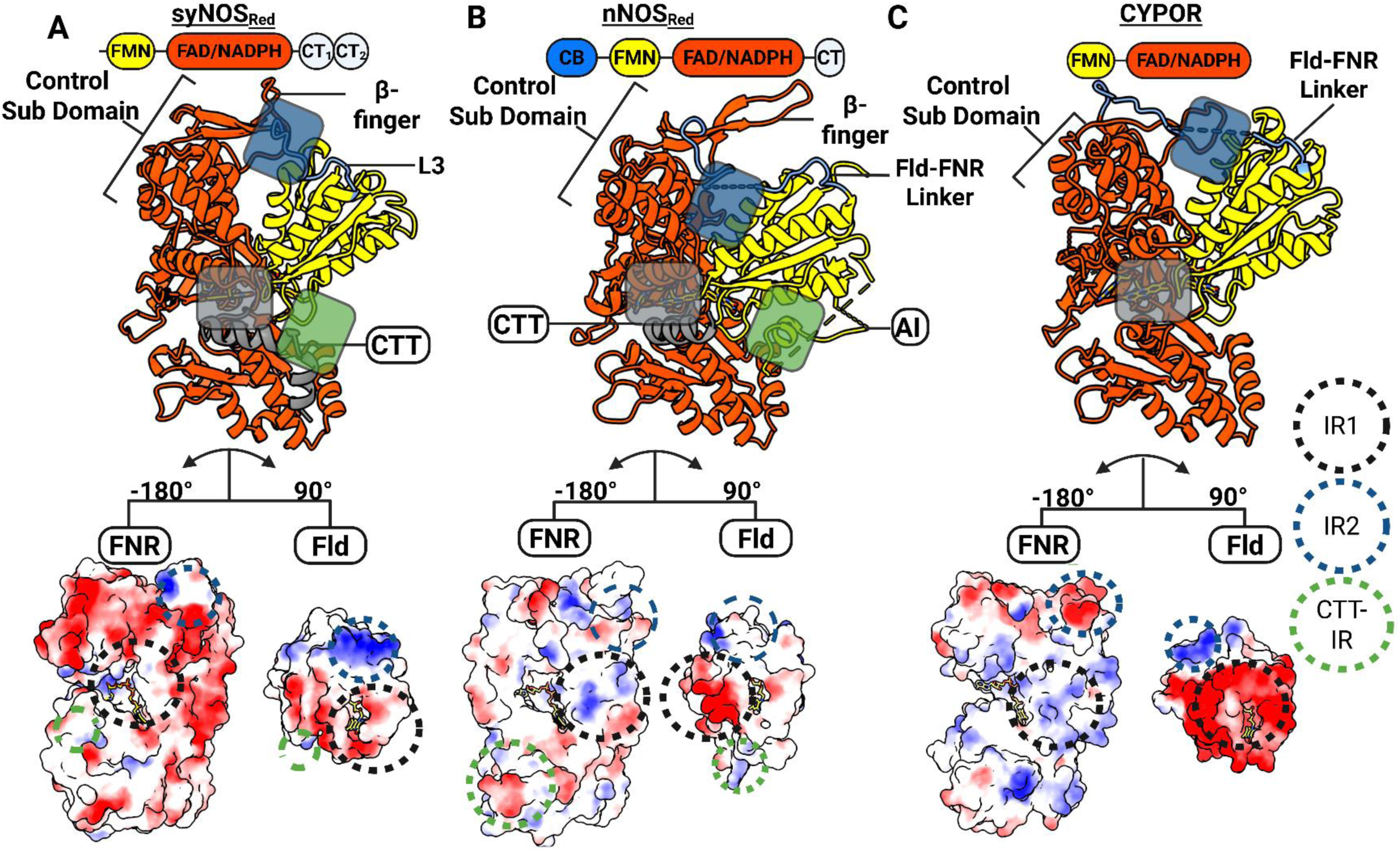
Structural comparison of syNOS_Red_, nNOS_Red_, and CYPOR. (A-C). The di-flavin closed conformation is defined by the L3 loop (situated between the Fld and the FNR domain, blue in Fig. 2A, B), the Fld domain, and what we define as Interfacing Region (IR) 1, IR2 and the CTT-IR. Domain architectures (top), the corresponding ribbon diagrams (middle), and key interfaces (lower) mediated by electrostatic interactions for syNOS_Red_ (A), nNOS_Red_ (PDB:1TLL, B) and CYPOR (PDB:1AMO, C) Surfaces are colored by coulombic electrostatic potential (blue – positive; white – neutral; red – negative)) with the Fld and FNR domains separated and rotated to view the interface. Interfacing Regions (IR) 1 (basic-to-acidic patch interactions between the FAD and FMN binding pockets), IR2 (Fld interaction-to-FNR-control region) and CTT-IR (CTT interaction with Fld) are denoted by colored circles in the electrostatic surface representation) and semi-transperant rectangles in ribbons.

### SyNOS_Red_ is distantly related, but structurally similar, to cytochrome P-450 reductase (CYPOR)

SyNOS_Red_, mNOS_Red_, sulfite reductase diflavin reductase (SIRFP), and methionine synthase reductase (MTRR) belong to the cytochrome P450 reductase (CYPOR) family of di-flavin reductases (Fig. S11) (*61–64*). Di-flavin reductases perform three-separate ET reactions i) hydride transfer from NADPH/NADH to the FNR FAD cofactor ii) sequential ET from the FNR to the Fld iii) ET from the Fld to an electron acceptor. Activities (i) and (ii) occur when the di-flavin reductase is trapped in a “closed” conformation permitting first hydride and then electron transfer along the sequence NADPH->FAD->FMN(H)• with activity (iii) requiring an “open” conformation that frees the Fld to reduce the acceptor (e.g. NOS_Oxy_ or cytochrome c).

Analysis by sequence clustering, phylogenetic estimation using maximum likelihood (PhyML) and multiple structure alignments (MSTA) indicated that syNOS_Red_ and mNOS_Red_ are outgroups relative to CYPORs (Fig. S11); yet, syNOS_Red_ is more closely related in structure to CYPOR than to nNOS_Red_ or other di-flavin reductases (Fig. 2 and S11) (*22*). Nevertheless, key differences arise in the structural and electrostatic features that stabilize the reductase closed conformation in syNOS (NOS-Sym) compared to CYPOR (Fig. 2A-C). CYPORs and mNOS_Red_ stabilize the closed conformation by way of an interfacing-region 1 (IR1) between an acidic patch surrounding the CYPOR_Fld_ FMN cofactor and a complementary basic patch surrounding the CYPOR_FNR_ FAD cofactor (Fig. 2B,C). An additional IR2 associates the Fld internal helices to the FNR control region. SyNOS_Red_ lacks electrostatic complementarity within IR1 and residue complementary within IR2. Instead, regions of the CTT (CTT-IR) and the C2 domain mediate interactions between the Fld and FNR in the syNOS_Red_ closed conformation (Fig. 3A).

**Figure 3.**
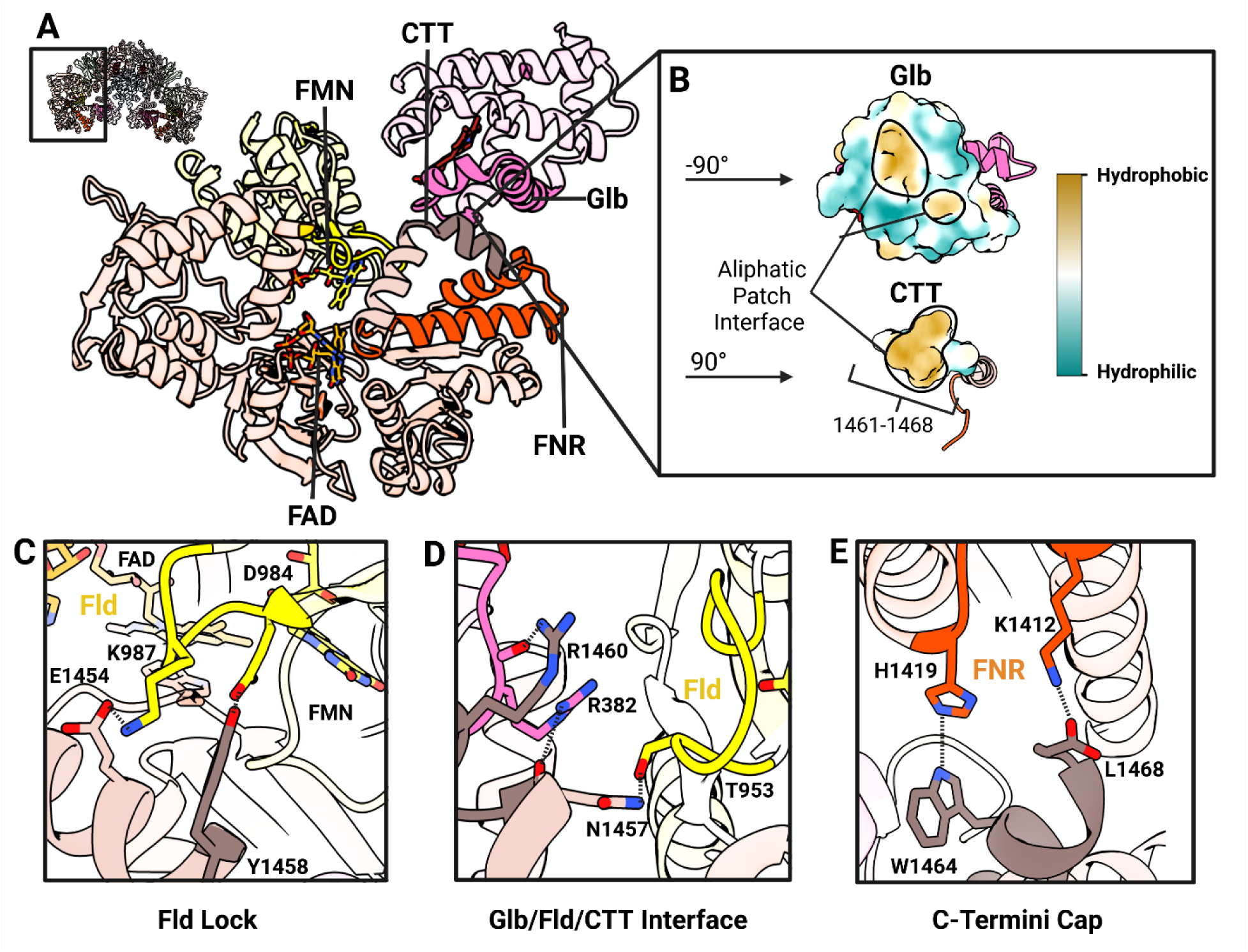
CTT interactions that stabilize the NOS_Red_-NOS_Glb_ locked state. (A) Ribbon diagram of syNOS_Red_ and syNOS_Glb_ from the boxed region of inactivated syNOS_FL_ (upper left inset) showing key stabilizing interfaces. (B) Aliphatic surface patch between syNOS_CTT_ and syNOS_Glb_ shown by hydrophobicity (yellow) after separating and rotating each domain to view the interface. (C) Fld-to-CTT lock interface mediated by Lys987(Fld)/Glu1454(CTT) salt bridge and Asp984(Fld)/Tyr1458(FNR) and Thr953/Asn1457 hydrogen bonds. (D) Fld-to-Glb interface mediated by Asn1457(CTT)/Thr953(Fld) hydrogen bond, Arg1460(CTT)/Arg382-carbonyl(Glb) hydrogen bond and Asn1457-carbonyl(CTT)/arg382(Glb) hydrogen bond. (E) C-Terminal cap interface mediated by Trp1419(CTT)/His1419(FNR) hydrogen bond and a Leu1468-Carboxyl(CTT)/Lys1412 salt bridge.

The CTT, shared by syNOS_Red_ and mNOS_Red_, are important for redox flux in NOS but are not found in canonical CYPORs (Fig. 2A-C) (*22*, *46*). These enzymes also share a β-finger (syNOS residues 1100-1132) that resides closer to the Fld domain in syNOS_Red_ (∼7.5 Å) compared to mNOS_Red_ (∼17 Å), thereby suggesting either a differing structural roles for this element or revealing a conformational state that nNOS_Red_ can also assume (Fig. 2C). syNOS_Red_ also lacks the mNOS_Red_ Fld autoinhibitory regulatory loop (AI, nNOS residues 836-849) that destabilizes CAM binding at low levels of Ca^2+^ (Fig 2B) (*61*, *65*). Although syNOS, nNOS_Red_, and CYPOR stabilize their closed conformations through different interactions, they all produce a coplanar orientation and similar separation between the FAD and FMN cofactors (∼13 Å from flavin N_5_-to-N_5_).

### NOS CTTs are enzyme-specific conformational regulators of ET

CTTs, thought to be an mNOS addition within the conserved CYPOR family (*66*), are 20-30 residue helical extensions that auto-inhibit reduction of NOS_Fld_ by NOS_FNR_ (Fig. 3). Sequence alignments show no discernable residue similarity between the syNOS CTT and CTTs of the three mNOS isoforms or that of the most similar eukaryotic NOS sequence from the diatom *T. minus* (Fig. S12). The lack of sequence similarity between syNOS_CTT_ and mNOS_CTT_, despite substantial similarity in the preceding syNOS_FNR_ domains, implies convergent evolution of these CTTs, likely in concert with the addition of syNOS_Glb_.

Unlike previous NOS_Red_ structures that capture only a portion of the CTT, the syNOS structure resolves the entire CTT and its interactions with syNOS_FLd_ and syNOS_Glb_ (Fig. 3, S13A). Compared to other NOSs, the 27-residue syNOS CTT divides into two perpendicular helices (CTT_1_ and CTT_2_; residues 1440-1468; Fig. 3A). CTT_1_ locks the position of syNOS_Fld_ in the canonical Fld-FNR closed conformation. CTT_1_ (residues 1446-1458) interacts with the FMN ligating loops of NOS_Fld_ (residues 984-989 and 951-960). In this interface, key residue contacts likely influence ET-competent conformational states of syNOS_Fld_ (Fig. 3C-D). CTT_1_ Lys987 likely modulates Fld dynamics by ordering residues that coordinate the FMN pyrophosphate groups. Indeed, the Lys987Ala variant, which disrupts the Lys987/Glu1454 salt-bridge, has a much more extended, flexible structure by small angle X-Ray scattering (SAXS) than does the WT (Fig. S14). The second CTT helix (CTT_2_, residues 1460-1468) is absent in other NOSs and CYPORs, likely because it binds to the atypical syNOS_Glb_ (Fig. 3A). Indeed, an aliphatic patch on CTT_2_ interacts with the hydrophobic crevice between residues 358-392 on syNOS_Glb_ (Fig 3A-B). Specific salt-bridges and hydrogen bonds among CTT_2_, syNOS_Glb_ and syNOS_Fld_, help to stabilize the Ca^2+^-free closed conformation of syNOS (Fig. 3D,E). In turnover conditions, XL-MS revealed a loss of crosslinks between the CTT and FNR, which underscores the role of the CTT in mediating dynamics of syNOS_Red_ (Fig S10C, Table S2).

SyNOS_Red_ has distinct reactivity toward the surrogate electron acceptor cytochrome c (Cc), compared to mNOS_Red_ and other CYPOR family members. The CTT lowers Cc reduction by NOS_Fld_ in both syNOS and nNOS, but the Ca^2+^ dependence is opposite. In the absence of Ca^2+^, removal of the CTT does not affect Cc reduction by syNOS (Fig. S15). Ca^2+^ decreases Cc reduction by almost 4-fold, but removal of the CTT reactivates Cc reduction (Fig. S15). In mNOS, Ca^2+^-CAM increases, rather than decreases Cc reduction (*67*, *68*) and CTT removal increases activity further. Furthermore, replacing the mNOS_FNR_ domain with CYPOR_FNR_, which lacks a CTT, switches the Ca^2+^-CAM dependence of mNOS such that Cc reduction now decreases ∼4-fold with Ca^2+^-CAM (*67*, *68*), much like for syNOS + Ca^2+^. These Cc reduction behaviors among NOS proteins likely reflect differences in the relative strength of interactions among NOS_Fld_, NOS_FNR_, the CTT, NOS_Oxy_ and Cc itself. Interestingly, syNOS structures in the Ca^2+^-depleted states (NOS-Sym, NOS-Asym, NOS-Mon) revealed a stable, closed Fld, whereas the Ca^2+^ turnover states (NOS-TO_1,2_) had a mobile undiscerned Fld domain, seemingly *more* conducive to Cc reduction, not less. However, in the Ca^2+^-state, the Fld domain may partition more frequently toward NOS_Oxy_, which would compete with Cc and thereby reduce Cc reduction rates. In mNOS, if the Fld-to-Oxy interaction is weaker than in syNOS. relieving a stronger CTT-to-Fld interaction with Ca^2+^-CAM may favor the Fld-to-Cc interaction.

### The CTT-stablizied closed conformation of syNOS_Red_ limits flavin reduction

SyNOS_FL_ and syNOS_Red_ rates of flavin reduction by NADPH, measured through anaerobic stopped-flow experiments, were biphasic and resembled the flavin reduction kinetics of CYPOR and nNOS_Red_ bound to Ca^2+^-CAM (Fig. S16A) (*69*). However, NADPH reduces syNOS_FL_ at a ∼74-fold slower rate than syNOS_Red_ implying that the closed conformation of syNOS_FL_ restricts hydride transfer from NADPH to FAD (Fig. S16B-D). The lower FAD reduction rate in syNOS_FL_ is surprising because syNOS_Red_ lacks an NADP(H)-ligating Arg (1400 in rat nNOS) known to inhibit electron flow in CAM-free nNOS/eNOS (*70*). In nNOS, the Arg1400Ser substitution increases flavin reduction while decreasing NO synthesis (*70*). Because syNOS contains an Asn residue at the 1400 position it would be predicted to have a faster flavin reduction rate than nNOS_Red_ (*70*). However, the syNOS_Glb_-to-CTT interaction, unique to syNOS, may especially stabilize conserved CTT residue Trp1440 (*71*), which blocks the nicotinamide group of NADPH from approaching the FAD isoalloxazine ring for hydride transfer (Fig. S16B). The CTT enforced rigidity in syNOS (also reflected in the XL-MS data, Fig. S10C) may compensate for the modified NADP^+^-binding pocket that lacks an autoinhibitory Arg1400 equivalent. Notably, compared to nNOS and CYPOR the nicotinamide moiety in syNOS-Mon adopts an atypical conformation that is intermediate to the reduction incapable NADP^+^ conformation seen in nNOS (PDB:1TLL) and the NADP^+^/FAD stacked conformation seen in the CYPOR W677X variants (PDB:1AJO, Fig. S16D).

### Ca^2+^ regulates syNOS activity through a C2-like domain

The lack of a CAM-CBD interaction in syNOS renders its Ca^2+^sensitivity enigmatic. However, syNOS_C2_ structurally resembles C2 domains that mediate membrane recruitment by binding Ca^2+^ (Fig. 4) particularly; mouse perforin C2 (Fig. 4A) (*56*, *57*, *72*).

**Figure 4.**
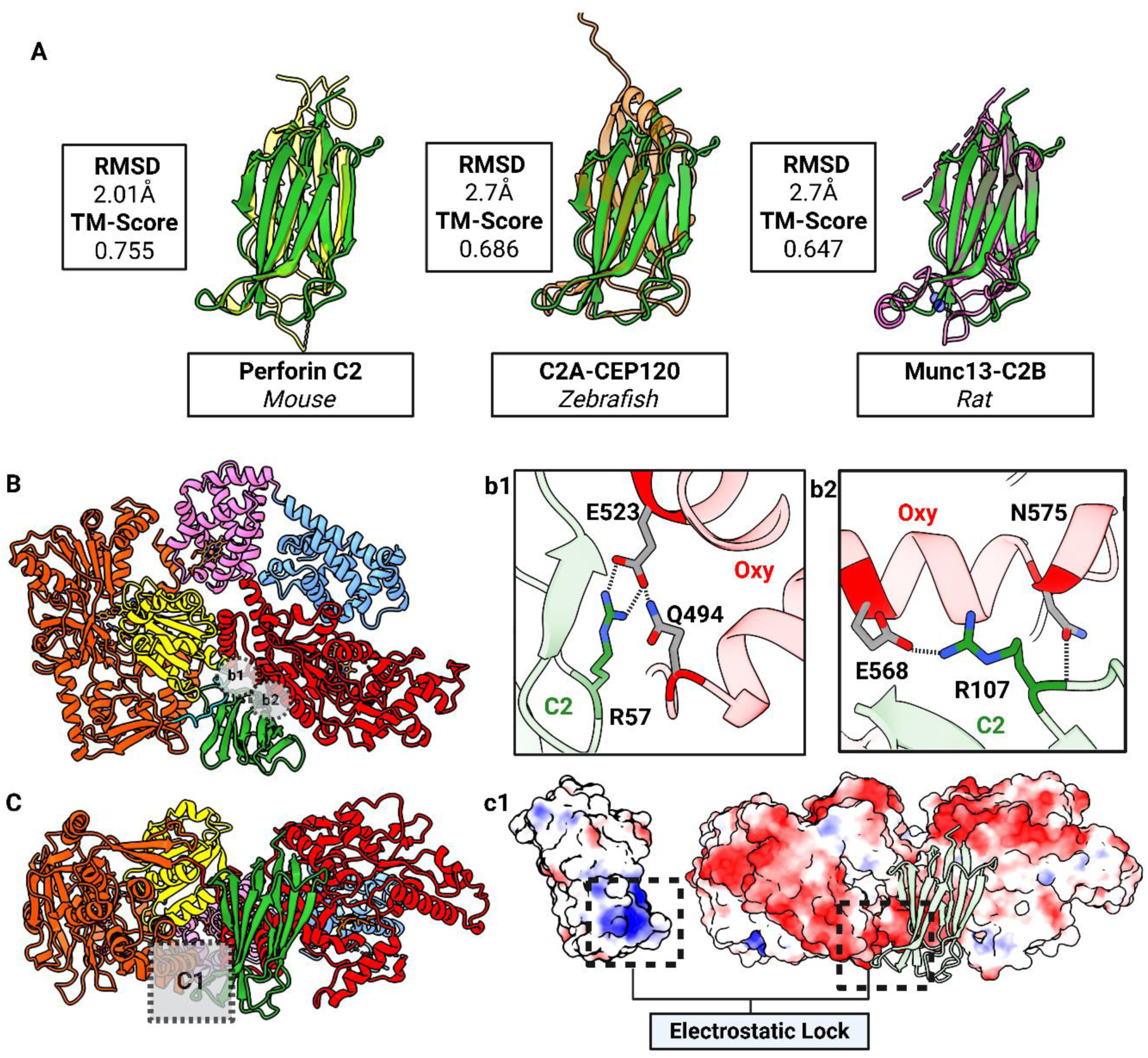
syNOS_C2_ interactions with NOS_FNR_ and NOS_Oxy_. (A) Superimposition of syNOS (C2, green) with Perforin C2 (PDB: 4Y1S, yellow), C2A-CEP120 (PDB:6EWL, orange) and Munc13-C2B (PDB:6NYT, magenta) as labeled, with root mean square deviation (RMSD) of Cα positions (RMSD), and Template Modeling (TM) scores boxed. B) Ribbon diagram of a syNOS subunit, colored by domain with key C2-Oxy interactions shown in insets b1 and b2. b1) Salt bridge between Arg57(C2)/Glu523(Oxy) with Glu523 stabilized by Gln494. b2) Salt bridge between Arg107(C2)/Glu568(Oxy) with the Arg107 amide stabilized by Asn575. (C) Ribbon diagram of a syNOS subunit showing the electrostatic lock interface between syNOS_C2_ and syNOS_Oxy_ in box inset c1. (c1) C2 rotated 90° (left) and FNR (right, orientation as in (C), colored by electrostatic surfaces (colored: blue-positive, white-intermediate and red –negative). C2 (green, ribbons) is also shown in its original position for reference.

In the closed conformation, syNOS_C2_ forms a tripartite interface with the syNOS_Oxy_ domain and the FNR control region (Fig. 4B). Two regions of salt-bridges mediate the C2-Oxy interface (Fig. 4B). In particular, the interaction between syNOS_C2_ Arg107 and NOS_Oxy_ Glu568, positions the C2 domain beside Trp715, the proposed site of mNOS_Fld_ binding onto mNOS_Oxy_ (*38*). A basic patch on the C2-domain and a complementary acidic patch on α-helix 1162-1173 of the FNR-control domain wedges the C2 between syNOS_Oxy_ and syNOS_Fld_ (Fig. 4C). Because syNOS_C2_ blocks the putative Fld-Oxy contact; displacement of syNOS_C2_ would decrease conformational restraints on syNOS_Red_ (particularly the L2 linker) and relieve steric hindrance on the backside of the heme pocket.

A variant devoid of the C2 domain (syNOS_DC2_, residues 179-1468) has no NOS activity but retains Cc-reduction activity (Fig. S14). Strikingly, syNOS_ΔC2_ has a high WT-like Cc-reduction rate that is uninhibited by Ca^2+^. In keeping with an enzyme unconstrained by inhibitory interactions. SAXS data indicates that the global flexibility of syNOS_ΔC2_ exceeds that of the WT and the more flexible K987A variant (Fig. S14). Thus, the C2 domain serves to stabilize NOS_Oxy_ and influences redox domain mobility of syNOS in response to Ca^2+^, akin to the functional role that Ca^2+^-CAM plays in the regulation of mNOSs. In other systems, Ca^2+^ binding to C2 domains neutralizes acidic regions to allow association with negatively charged phospholipid head-groups (*73*). Although, syNOS_C2_ lacks the typically conserved Asp residues that coordinate Ca^2+^, isothermal calorimetry (Fig. S17) indicated that the syNOS_C2_ binds Ca^2+^ with a 1.6 μM Kd, which compares to the Ca^2+^ affinity of PLDβ C2 domains (0.8 μM) (*74*).

Various stimuli produce surges in cyanobacterial intracellular Ca concentrations from 100-200 nm to the μM range (*75*, *76*). In vitro, syNOS activity responds to Ca^2+^ with a K_M_ of ∼ 200 μM (*34*). This larger value may reflect the conformational coupling of the C2 domain movement to Ca^2+^ binding and may be affected by the cellular context. Interestingly, some types of cyanobacteria uptake Ca^2+^ at very high concentrations to mineralize CaCO_3_ (*77*).

### Ca^2+^ causes a large movement of the C2 domain

In the presence of Ca^2+^ and L-arginine, syNOS adopts a more flexible state, wherein most of syNOS_Red_ is undiscerned by cryoEM (NOS-TO_L,T_). In this turnover state, the C2 domain has moved away from the Oxy/FNR interface and now binds at the entrance of the syNOS_Glb_ heme pocket, nearly 85 Å from its original position (Fig. 5A). The new C2 position blocks the syNOS_Glb_ heme-binding center and substantially shifts the αB-D helices and constituent residues that compose the distal heme pocket where NO is oxidized (Fig. 5B). The highly conserved C2 basic patch (Lys35, Lys37 and Arg81) that formally bound syNOS_FNR_ in the locked dimer interacts with the heme-propionic acid groups of syNOS_Glb_ and the 81-89 loop, also conserved among syNOS_C2_ domains, reaches over to block the distal heme pocket. Furthermore, the movement of syNOS_C2_ displaces the L2 linker, which swivels by ∼135° and now directs syNOS_Fld_ toward the backside of the heme pocket on the opposing subunit (Fig. 5C), consistent with the trans ET mechanism characteristic of mNOSs (*58*). As syNOS_C2_ now resides where the electron-donating FNR modules of Hmps interact with their cognate globin domains, we tested whether removal of the C2 domain (syNOS_DC2_) affected syNOS_Glb_ reduction in syNOS_FL_. Surprisingly, we observed little change in syNOS_Glb_ reduction rates in syNOS_ΔC2,_ nor did Ca^2+^ affect syNOS_Glb_ reduction rates in syNOS_FL_ (Fig. S18).

**Figure 5.**
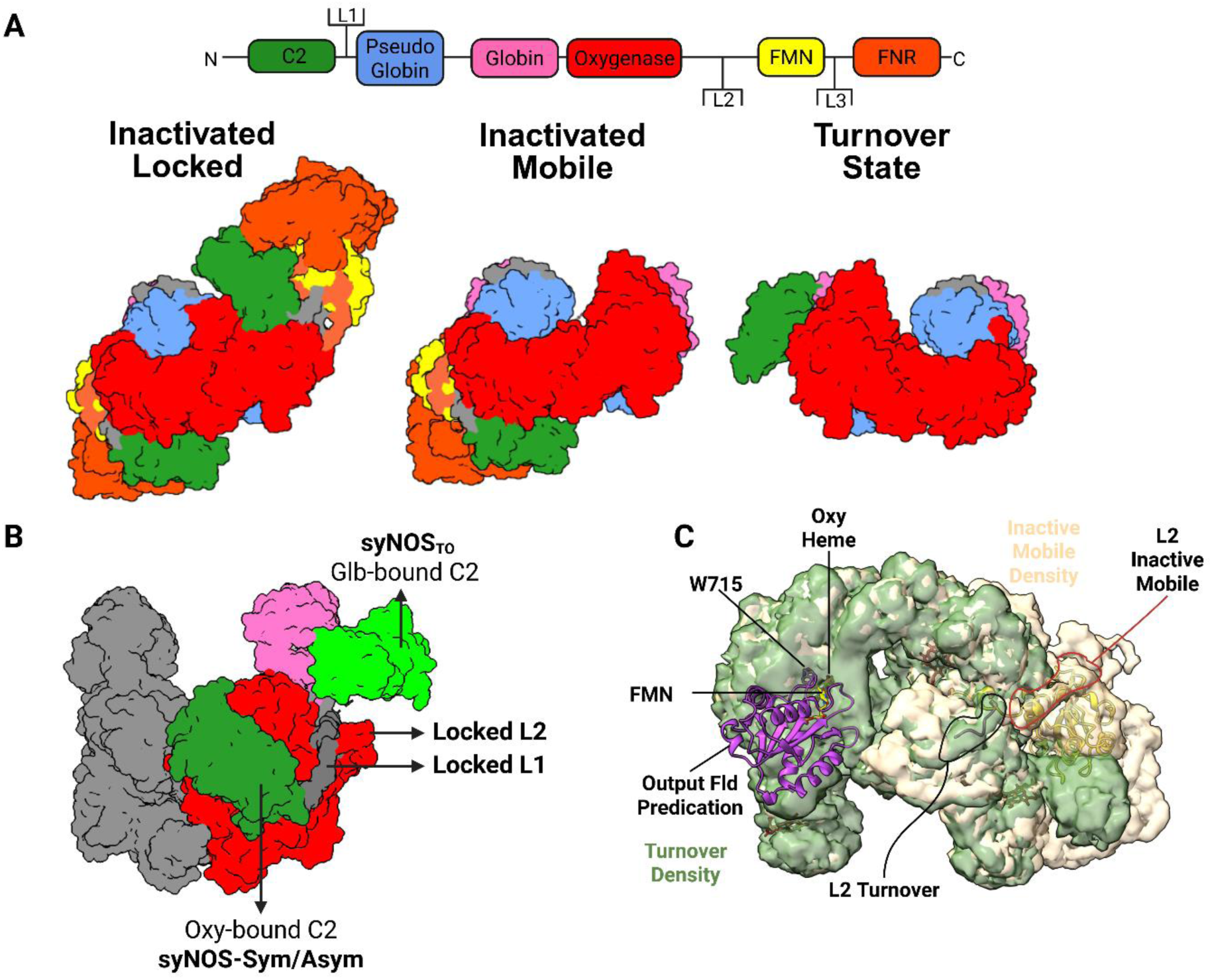
Ca^2+^-activated C2 mobility. (A) syNOS gene architecture colored by domain, with linkers labeled L1-3(above). Surface representation of three separate syNOS states with relative orientations of the C2 domain. (B) Superimposed surface representations of the turnover state (C2 – lime green) and inactivated states (C2 – forest green, Oxy – red and Glb – hot pink) reveal dramatically different C2 positions. (C) L2 in the turnover state (tan electron density) connects to the Fld (yellow ribbons) but then swivels 135° (green density) from its position in the inactivated state to direct the NOS_Fld_ to the adjacent NOS_Oxy_ subunit (purple ribbons of AlphaFold prediction).

### syNOS_Glb_ reduction requires an altered configuration of syNOS_FNR_

Flavohemoglobins reductively activate dioxygen with an FNR domain in order to oxidize NO to NO_3_^−^; however, in principle, either syNOS_Fld_ or syNOS_FNR_ could reduce syNOS_Glb_. In the locked state, both the syNOS_Fld_ FMN and the syNOS_FNR_ FAD are too far from the syNOS_Glb_ Fe^3+^ to yield reasonably fast ET rates (27 Å and 31 Å, respectively). When considering that syNOS_Glb_ has a lower potential (E^0’^ = –250 mV) than syNOS_Fld_ (E_m_ = –170 mV), but not syNOS_FNR_ (E_m_ = –290 mV), syNOS_FNR_ is likely the better electron donor. However, the relatively low expected driving force for reduction between syNOS_FNR_ and syNOS_Glb_ (Table S5) and a distance of separation of ∼31 Å, suggests that syNOS_FNR_ displaces from its position in NOS-Sym to reduce syNOS_Glb_ at the observed rate (Fig. 6; (*78*)). Indeed, the isolated syNOS_FNR_ domain reduces the syNOS_Glb_ domain with saturation behavior (K_m_ = 11.5 μM) at nearly the same rate (k_cat_ = 0.25 s^-1^) as syNOS_Glb_ is reduced in full-length Ca^2+^-free syNOS (k_obs_ = 0.22 s^-1^, Fig. S18). Furthermore, if syNOS_Fld_ in the locked confirmation can directly reduce syNOS_Glb_, we would expect the conformationally destabilized syNOS_ΔC2_ to show slower syNOS_Glb_ reduction, but this is not the case (Fig. S16). A modest decrease in syNOS NOD activity with Ca^2+^ suggests an inhibitory effect of syNOS_C2_ (Fig. S18), which would be expected from its relocation to the site where FNR domains typically bind globin domains in Hmps.

**Figure 6.**
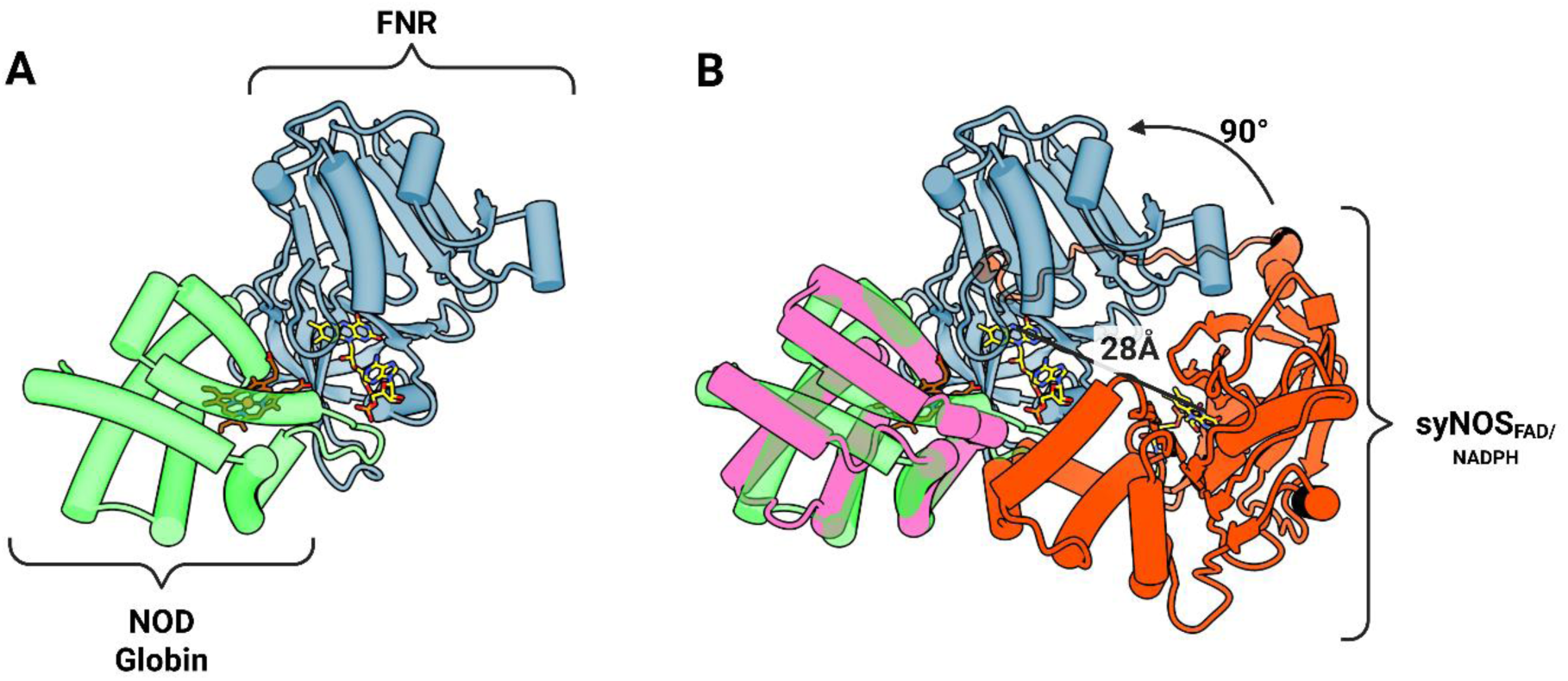
syNOS_FNR_ likely requires a large conformational change to reduce syNOS_Glb_. (A) Cartoon representation of E.*coli* Hmp (PDB: 1GVH) with the NOD domain (lime green) and FNR (metallic blue) domains orientated to show the Heme-to-FAD separation of 17.8 Å. (B) syNOS_Glb_ (purple) and associated syNOS_FNR_ domain (red, control region 1084-1224 not shown) superimposed on the Hmp Glb (green) and associated FNR (blue). SyNOS_FNR_ would need to rotate 90° and translate 28 Å to adopt an equivalent Hmp conformation.

### The asymmetric state may act as a conformational intermediate between NOS and NOD activities

The mNOS input, intermediate, and output states are not discrete, but sampled within a continuum of motion (*18*, *35*). In the output state, Fld-CAM moves 70 Å to dock onto an opposing NOS_Oxy_ domain at the backside of the heme pocket. Similarly, syNOS-Asym (PDB:9Q0Y,9Q15) represents an intermediate structural state between the locked dimer and the turnover state that is likely in dynamic equilibrium with NOS-Sym and NOS-TO. An Alphafold3 prediction of the Oxy dimer unit with one Fld domain places syNOS_Fld_ at the experimentally validated position on syNOS_Oxy_ found by HDX-MS and crosslinking studies of mNOS (*38*). Upon superimposing this prediction onto either NOS-Asym or NOS-Sym, the Fld binding surface on syNOS_Oxy_ is blocked by the syNOS_C2_ (Fig. 5C and S19). Furthermore, compared to Hmp, syNOS_FNR_ would have to rotate 90° and move 28 Å to reduce syNOS_Glb_ (Fig. 6). This configuration is likely accessible in the dynamic asymmetric subunit but would be prevented in syNOS_TO_ (PDB:9Q0X) by the rearranged syNOS_C2_. Hence, it is likely that syNOS-Asym is capable of syNOS_Glb_ reduction but not syNOS_Oxy_ reduction.

The intermediate conformation of syNOS-Asym (PDB:9Q0Y,9Q15) may be important for the switch between NOS and NOD activities and explain the inhibitory effect of Ca^2+^ on Cc-reduction. In the absence of Ca^2+^, syNOS-Sym locks down and inhibits NOS activity, but NOS-Asym allows syNOS_FNR_ to reduce syNOS_Glb_, and syNOS_Fld_ to reduce Cc. In syNOS-Asym (PDB:9Q0Y,9Q15), syNOS_Fld_ cannot reduce syNOS_Oxy_ because syNOS_C2_ blocks syNOS_Fld_ from the syNOS_Oxy_ interaction site on the opposing subunit. Ca^2+^ promotes the relocation of the C2 domain from syNOS_Oxy_ to syNOS_Glb_; however, some C2 displacement from NOS_Oxy_ may occur in the absence of Ca^2+^ because the C2 domain is not well defined in the more mobile subunit of syNOS-Asym. In addition to preventing syNOS_FNR_ from reducing the syNOS_Glb_, C2 movement disrupts L2, destabilizing the interaction of syNOS_Fld_ with syNOS_FNR_ and favors partitioning of syNOS_Fld_ toward the unblocked syNOS_Oxy_ and away from Cc (Fig. 7).

**Figure 7.**
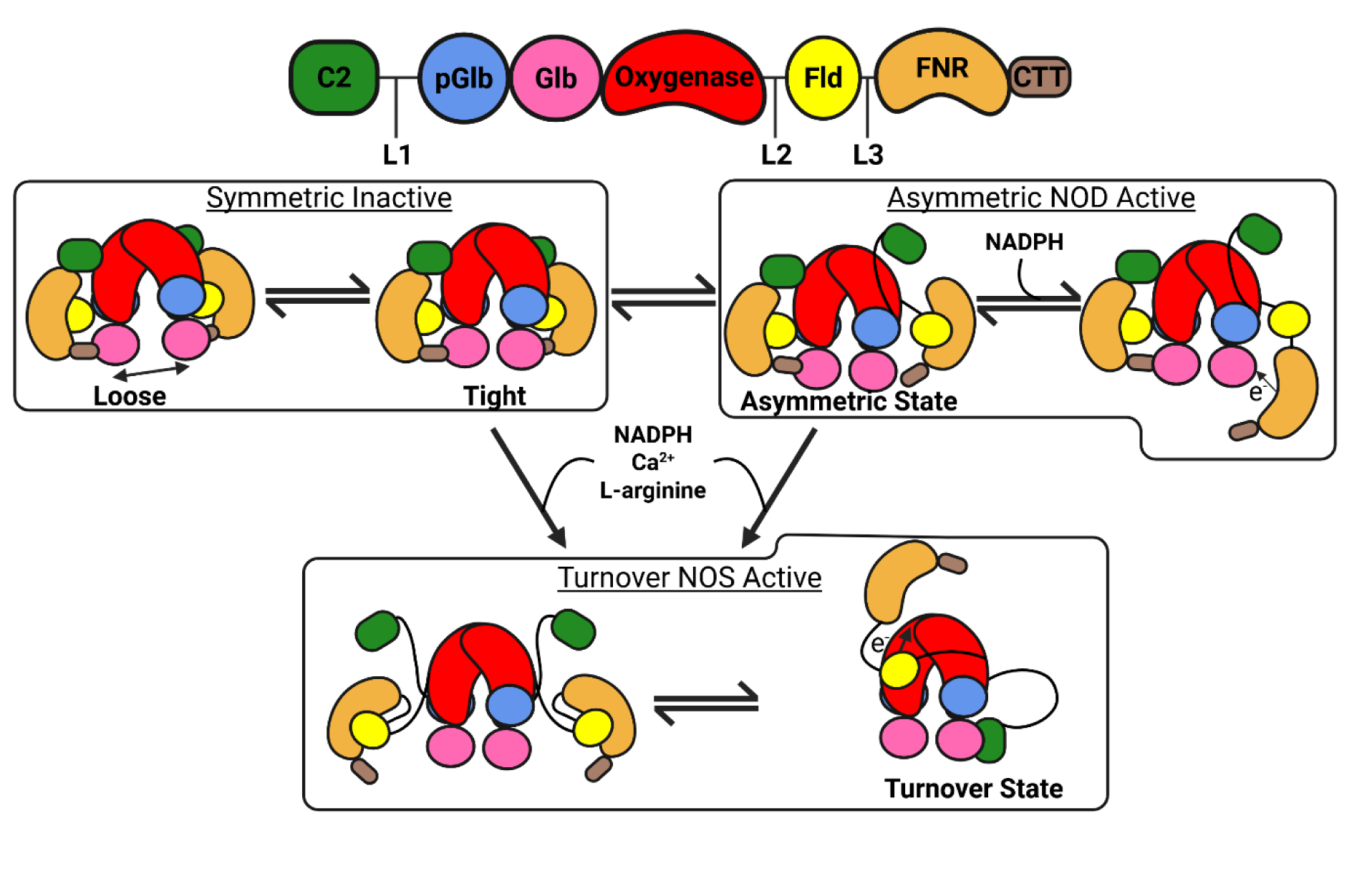
Model for NOS to NOD activity switching in syNOS. In the inactivated locked state syNOS_Oxy_ equilibrates between a loose and tight dimer, the latter stabilized by BH_4_. This state can transition into the asymmetric NOD active state which involves a release of a C2 domain from one subunit and increased mobility of NOS_Fld_ and NOS_FNR_, such that NOS_FNR_ can reduce the syNOS_Glb_ to activate NOD activity. SyNOS_Glb_ reduction coincides with a loosening of the CTT to allow hydride transfer from NADPH to FAD and movement of syNOS_FNR_. In the presence of Ca^2+^/NADPH/L-arginine syNOS rearranges to facilitate NOS activity. The displacement of syNOS_C2_ from NOS_Oxy_ and its binding to NOS_Glb_ relieves constraints on L2 and exposes the Trp715 site such that NOS_FNR_ can relocate to NOSOxy on the opposing subunit and reduce the NOS_Oxy_ heme for NO production.

Similar to mNOS, syNOS assumes a continuum of conformations that circumscribe defined enzymatic states. Hence to generate changes in physiologically relevant activities, the relative weighting of states within the conformational ensemble must be regulated and responsive to cellular context. It is striking that the syNOS_C2_ moves between a location that blocks syNOS_Oxy_ reduction to one that blocks syNOS_Glb_ reduction, thereby suggesting counter regulation of NO synthesis and NO oxidation. Such a mechanism may allow the enzyme to tune the balance of NO production versus NO detoxification, which would be of benefit for signal transduction, nitrogen metabolism and cellular homeostasis. Overall, these syNOS structural studies demonstrate how a multidomain reductase and its associated regulatory components can undergo large scale rearrangements to modulate reactivity in response to environmental conditions.

## Methods

### Molecular cloning

The native coding sequence for full-length syNOS (Uniprot ID: B4WU43) was PCR-amplified and cloned into a modified pet28-a construct between restriction sites NdeI and XhoI using NEBuilder HiFi DNA Assembly master mix (E2621L). The construct contained sequences for an N-terminal Twin-Strep Tag with a TEV cleavage site. To produce the hptG and gapA construct, htpG and gapA were amplified from K-12 *E. coli* and placed into the MCS-1 and the MCS2 of a tagless pAYC-Duet vector using NEBuilder HiFi DNA Assembly master mix (E2621L). Individual point mutants were constructed using Kinase-Ligase-DpnI (KLD) method using primers in Table S4 (New England Biosciences). All plasmid molecular cloning and plasmid material generation were carried out with NEB® 5-alpha competent *E. coli* (New England Biosciences). Successful transformants were validated through Sanger Sequencing or Nanopore sequencing (Eurofins Genomics).

### Protein expression and purification

SyNOS and/or variants were expressed in BL21(DE3) as previously described (*34*) with the modification of co-expression with hptG/gapA, which improved yield and reproducibility. Cell pellets were stored at –80 °C prior to purification. The pellet was thawed and resuspended in lysis buffer (50 mM HEPES (pH 7.5), 150 mM NaCl, 1 mM TCEP, 10% glycerol and 10 uM BH_4_), followed by high-speed centrifugation at 100 000 G and application on Strep-Tactin XT 4-Flow resin (IBA Life Sciences). Elution of the affinity-bound protein fraction was carried out in batch with 50 mM biotin supplemented lysis buffer. Post elution samples were concentrated using Viva-Spin columns (100 kDa cut –off), before application onto 10/300 Superose 6 Increase Column (Cytiva) using Gel Filtration Buffer 1 (GFB1, 25 mM HEPES (pH 7.5), 150 mM NaCl, 1 mM TCEP, 10% glycerol and 10 uM BH_4_).

### Size exclusion chromotography – Small angle X-ray scattering (SEC-SAXS)

All SEC-SAXS was performed at ID7A1 beamline at Cornell High Energy Synchrotron Source (CHESS). Photon flux was 2.3 x 10^12^ ph/s and scattering data was collected 0.005 Å^-1^ ≤ q ≤ 0.7 Å^-1^ on an EIGER 4M detector (Dectris). Samples were injected into an inline 3.2/150 Superdex 200 column in gel filtration buffer. The entire SEC-SAXS trace was captured using a 2s per exposure throughout the elution. The flow-cell format of the experiment should limit x-ray photoreduction of the sample, but it is possible that the NOS cofactors undergo some photoreduction during the experiment (*79*). All SEC-SAXS data processing was performed through RAW (*80*).

### Single turnover anaerobic globin reduction

Anerobic syNOS_Glb_ heme reduction assays were carried out using an Agilent 8453 UV-Vis spectrometer with an attached Hewlett Packard 89090A Peltier temperature controller for maintaining constant temperature at 20 °C and constant stirring at 400 rpm (Agilent & Hewlett Packard). Prior to reaction initiation with 1 mM NADPH, syNOS or variants (1 μM) were incubated anaerobically in the presence of the Glucose Oxidase/Catalase (GOD/CAT) oxygen scavenging system, as previously described (*81*). Samples were run in either in the presence of 1 mM Ca^2+^ or 1 mM EGTA, as noted. In the case of syNOS_FNR_ (1027–1468)/syNOS_Glb_ (337–469) reduction trials, syNOS_Glb_ was fixed at 2.2 μM and syNOS_FNR_ was varied between 1 μM to 21 μM with no EGTA/Ca^2+^ present. All data was processed using home-made scripts in R Studio (Posit PBC) with rates of syNOS_Glb_ reduction with to a single exponential to determine k_obs_. For the truncated domains k_cat_ and K_m_ were calculated by plotting k_obs_ for each syNOS_FNR_/syNOS_Glb_ pair against syNOS_FNR_ concentration using Prisim (GraphPad).

### Stopped-flow pre-steady state flavin reduction kinetics

Anerobic stopped-flow mixing of NADPH with syNOS_Red_ or syNOS_Fl_ in the absence of Ca^2+^ was used to characterize flavin reduction rates. After purification, syNOS_Red_ contains oxidized FAD and FMN. Attempts to pre-reduce the enzyme to FMNH• with sodium dithionate, DTT and ascorbate caused precipitation without an observable spectral shift to FMNH•. Thus, flavin reduction rates reflect conversion from fully oxidized syNOS_Red_. All stopped flow experiments were carried out anaerobically using a Stopped Flow SX20 instrument (Applied Photophysics). Samples were pre-incubated anaerobically in the presence of the GOD/CAT oxygen scavenging system (see above). Changes in absorbance were detected using a single wavelength Photomultiplier Tube detector. syNOS_FL_ (10 μM) / syNOS_Red_ (40 μM) were mixed with 4x excess NADPH. All data was processed using home-made scripts in R Studio (Posit PBC).

### CryoEM grid preparation and data collection – syNOS ligand free (Dataset 1 and 2)

A 4 μl volume of 1.0 mg ml^−1^ syNOS solution, in GFB1, was applied to glow-discharged grids (200 mesh Quantifoil Cu, R1.2/1.3; Electron Microscopy Sciences). After 10 s of incubation, the excess solution was blotted for 3.5 s with filter paper by a Vitrobot Mark IV and plunged into liquid ethane, for vitrification (ThermoFisher). CryoEM images of syNOS were collected on a Talos Arctica (ThermoFisher) operated at 200 keV at a nominal magnification of 63,000× with a Gatan GIF Quantum LS Imaging energy filter, using a Gatan K3 direct electron camera in super-resolution counting mode, corresponding to a pixel size of 1.31 Å. A total of 1214 images stacks were obtained with a defocus range of −0.8 to −2.0 μm by Serial EM (University of Colorado-Boulder). Each stack video was recorded for a total dose of 50 electrons per Å^2^.

### CryoEM grid preparation and data collection – Ca^2+^ and L-arginine conditions (Dataset 3)

A 4 μl volume of 1.0 mg ml^−1^ syNOS in the presence of 1 mM Ca^2+^ and 1 mM L-arg solution was applied to glow-discharged grids (300 mesh Quantifoil Au, R1.2/1.3). After 10s of incubation, the excess solution was blotted for 2.5 s with filter paper by a Vitrobot Mark IV and plunged into liquid ethane for vitrification (ThermoFisher). CryoEM images of syNOS were collected on a Talos Krios (ThermoFisher) operated at 300 keV at a nominal magnification of 165,000× with a Gatan GIF Quantum LS Imaging energy filter, using a Gatan K3 direct electron camera in super-resolution counting mode, corresponding to a pixel size of 0.422 Å. A total of 5019 images stacks were obtained with a defocus range of −0.6 to −2.0 μm by Leginon (*82*). Each stack video was recorded for a total dose of 50.5 electrons per Å^2^.

### CryoEM grid preparation and data collection – turnover conditions (Dataset 5)

A 4 μl volume of 1.0 mg ml^−1^ syNOS-ligand free solution in GFB1 was applied to glow-discharged grids (200 mesh Quantifoil Cu, R1.2/1.3). After 10s of incubation, the excess solution was blotted for 3.5 s with filter paper by a Vitrobot Mark IV and plunged into liquid ethane for vitrification (ThermoFisher). CryoEM images of syNOS were collected on a Talos Krios (ThermoFisher) operated at 300 keV at a nominal magnification of × 165,000 with a Gatan GIF Quantum LS Imaging energy filter, using a Gatan K3 direct electron camera in super-resolution counting mode, corresponding to a pixel size of 0.730 Å. A total of 5001 images stacks were obtained with a defocus rage of −0.6 to −2.0 μm by Leginon (*82*). Each stack video was recorded for a total dose of 44.9 electrons per Å^2^.

### CryoEM Image processing and structural reconstruction

All cryoEM density refinement was performed using CryoSPARC (*50*). In brief, micrograpghs were motion corrected using patch motion followed by CTF estimation using patch CTF. Particles were picked using CRYOLO and run through 2D classification and initial density reconstructions. Candidate densities were then selected for further refinement, including junk classes to eliminate contaminating/damaged particles. For dataset dependant details please refer to Figs. S2-S6. as illustrated by the procedures illustrated in Figs. S2-S6.

For syNOS-Asym (DS4, EMD-72110), non-uniform refinement obscured the smaller subunit. To improve the density quality we locally refined each subunit. ChimeraX was used to create the two masks, centered at each subunit, for the local refinement(Fig. S5). Gaussian noise with a standard deviation of 3 was applied to EMD 72110, producing a new density that obscured secondary structure features. The Segger tool (*83*) was used to segment the map into each subunit. composite map (EMD-72116) produced from DS4, The electron density was resampled into the appropriate box size and imported into CryoSPARC where masks were created using the 3D Volume tool and subsequently used for local refinement. The refined maps were aligned and merged using the vop maximum command in ChimeraX to produce the composite map.

An Alphafold3 prediction of a syNOS homodimer (residues 1-1468), was used as the input model and docked into the appropriate electron density using ChimeraX (*50*, *84*, *85*). Ligand and poor fitting regions were built and real-space refined using with Coot (*85*). The resulting models were passed through ISOLDE (*86*) to approach thermodynamically rational rotamers/conformations. The outputs from these initial models were subjected to a series of real-space refinement with PHENIX (*87*) and conformational outlier residues were then adjusted manually using Coot (*85*).

### Cytochrome c reduction assay

Cytochrome c (Cc) was used as an electron acceptor to measure the efficiency of electron flow from NOS_Red_. Cc accepts an electron from the fully reduced FMNH_2_ and not from FADH_2_/FADH^-^/FAD^•^/ FMNH^•^. Cc reduction assays were carried out as previously described (*88*) using an Agilent 8453 UV-Vis Spectrometer with an attached Hewlett Packard 89090A Peltier Temperature Controller for constant temperature at 20 °C and constant stirring at 400 rpm (Agilent & Hewlett Packard). All reported reactions were carried out in CC buffer (100 mM Tris (pH 7.5) and 200 mM NaCl). Cc was pre-oxidized with 1 mM potassium ferrocyanide and then buffer exchanged into CC buffer. Reactions were conducted with 10 uM Cc and 3 nM of the sample of interest. Reactions were initiated with 1 mM NADPH and were run either in the presence of 1 mM Ca^2+^ or 1 mM EGTA as noted. All data was processed using home-made scripts in R Studio (Posit PBC).

### Nitric oxide synthase activity assay

All activity assays were carried out for 30 minutes in 100 mM Tris, (pH 7.5) 200 mM NaCl, 1 mM DTT and 1 mM L-arginine and initiated with 1 mM NADPH (25 °C). Each sample was evaluated in the presence of 1 mM Ca^2+^ or 1 mM EGTA. Post reaction, samples were boiled at 70 °C to halt NOS activity. NADPH was then precipitated using the ZnCl_2_/NaCO_3_ method and subsequently centrifuged at 15000 G at room temperature. The supernatant was then quantified for total NO_2_^-^ and NO_3_ ^-^ content through the Griess assay as implemented in the Nitrate/Nitrite Colorimetric Assay Kit (Cayman Chemicals).

### Oxygen consumption assay to probe Nitric Oxide Dioxygenase activity

The Oxygraph+ system (Hansatech Instruments) was used to measure oxygen consumption of syNOS in the presence of the NO donor, 3-(2-Hydroxy-1-methyl-2-nitrosohydrazino)-N-methyl-1-propanamine (NOC-7). 1 μM of enzyme (syNOS_Fl_ or syNOS_ΔC2_) was incubated in buffer containing 50 mM Tris and 150 mM NaCl at pH 7.5, with 10 μM NOC-7 for 5 minutes. The reaction was initiated with 1 mM NADPH. Each sample was evaluated in the presence of 1 mM Ca^2+^ or 1 mM EGTA. Three separate controls were performed to rule out superoxide or non-enzymatic NO oxidation: NOC-7 + NADPH, NOC-7 + syNOS_FL_ and syNOS + NADPH.

### Isothermal calorimetry

Thermodynamic binding properties between Ca^2+^ and syNOS_C2_ (1–138) were measured on a Nano ITC Low Volume isothermal titration calorimeter (TA instruments). SyNOS_C2_ was prepared in a 50 mM CHES and 150 mM NaCl buffer at pH 9.0 at 0.072 mM with 200 μL in the sample cell. Lower pHs also gave indication of Ca^2+^ binding, but led to aggregation of the C2 domain that prevented quantitative assessment. CaCl_2_ was prepared in the same buffer at a concentration of 0.250 mM; 2.5uL was titrated every 250 s for a total of 30 injections. Thermal data were analyzed using an independent binding model with the NanoAnalyze software.

### Reduction potential determination

Measurements were performed using the isolated domains syNOS_Glb_ (337–469), syNOS_FMN_ (856–1027) and syNOS_FNR_ (1027–1468), which were expressed and purified separately. Reduction potential measurements were performed using a dye calibration method (*89*, *90*), wherein reduction was driven by a deazaflavin/EDTA photochemical system in a 50 mM NaPO_3_, 10% Glycerol at pH 7.0, with the following modifications: i) 9 μm 5-deazariobflavin and 5 mM EDTA was used as the chemical reductant ii) reactions were equilibrated anaerobically in a coy chamber while supplemented with GOD/CAT oxygen scavenging system. All measurements were carried out using an Agilent 8453 UV-Vis spectrometer with an attached Hewlett Packard 89090A Peltier Temperature Controller for maintaining constant temperature at 20 °C and constant stirring at 400 rpm (Agilent & Hewlett Packard), with the reaction being photoinitiated with a 15 mW 300 nm laser (ThorLabs) placed above the clear septum capping the cuvette.

Calibrating dyes per domain were selected based on literature values of the published redox potentials (Table S5). Note that the quoted reduction potentials for syNOS_Fld_ and syNOS_FNR_ will be the average of the single electron couples between semiquinone and oxidized, and hydroquinone and semiquinone.

### DSBU crosslinking, in-gel protein digestion and extraction

Purified syNOS that correspond to the dimer fraction from SEC was split into two conditions: 1) inactive – GFB1, 2) turnover – GFB1 + 1mM Ca^2+^ + 1 mM NADPH + 100 μM BH_4_. Both conditions were then incubated with 1mM (100 molar excess) disuccinimidyl dibutyric urea (DSBU, ThermoFisher) for 30 minutes at room temperature prior to quenching with 20 mM Tris pH 8.0. Samples were then centrifuged and run on an SDS-PAGE gel using lameli sample buffer.

Bands corresponding to dimers were excised, digested and extracted as previously described (*91*), apart from the proteases of choice being Lys-C/Trypsin, prior to LC-MS/MS analysis.

### Crosslinks identification by nanoLC-MS/MS

The digested product was characterized by NanoLC-MS/MS analysis at the Cornell Biotechnology Resource Center. The analysis was carried out using an Orbitrap FusionTM TribridTM (Thermo-Fisher Scientific, San Jose, CA) mass spectrometer equipped with a nanospray Flex Ion Source, and coupled with a Dionex UltiMate 3000 RSLCnano system (Thermo, Sunnyvale, CA). Each sample was loaded onto a nano Viper PepMap C18 trapping column (5 µm, 100 µm’ 20 mm, 100 Å, Thermo Fisher Scientific) at 20 μL/min flow rate for rapid sample loading. After 3 minutes, the valve switched to allow peptides to be separated on an Acclaim PepMap C18 nano column (2 µm, 75 µm x 25 cm, Thermo Fisher Scientific) at 35 °C in a 93 min gradient of 5% to 40% then 93-98 min of 40%-60% buffer B (95% ACN with 0.1% formic acid) at 300 nL/min. The Orbitrap Fusion was operated in positive ion mode with nano spray voltage set at 1.5 kV and source temperature at 275 °C. External calibrations for Fourier transform, ion-trap and quadrupole mass analyzers were performed prior to the analysis. Samples were analyzed using the HCD-MS2 workflow, in which MS scan range was set to 300-1800 m/z and the resolution was set to 60,000. Precursor ions with charge states 2-8 were selected for HCD MS2 acquisitions in Orbitrap analyzer with a resolution of 30,000 with collision energy of 25% and 30%, normalized AGC of 200%. The precursor isolation width was 1.6 m/z and the maximum injection time was 70 ms. All data were acquired under Xcalibur 4.4 operation software (Thermo-Fisher Scientific).

### Crosslink identification and visualization

Raw spectra were searched using Proteome Discoverer 2.4 (Thermo-Fisher Scientific, San Jose, CA) with XlinkX v2.0 algorithm for identification of crosslinked peptides. The search parameters were as follow: four missed cleavages for either double digestion or triple digestion with fixed carbamidomethyl modification of cysteine, variable modifications of methionine oxidation, asparagine/glutamine deamidation, protein N-terminal acetylation, serine/threonine/tyrosine phosphorylation and lysine di-glycine tag. Only lysine-to-lysine crosslinks were considered. The peptide mass tolerance was 10 ppm, and MS2 fragment mass tolerance was 0.6 Da. A custom database (246 sequences) containing syNOS, common *E.coli* chaperones and common contaminates was used for the search, with > 1% FDR for report of crosslink results. Identified crosslinked peptides were filtered for Max. XlinkX Score > 40, a confidence score that reflects how well peptide spectrum matching, peptide fragmentation and crosslink doublet identification. Spectra were manually analyzed to validate potential crosslinks. The crosslinked peptides were analyzed using XMAS (*93*).

## Acknowledgements

We thank Angela Picciano for helpful discussions. We thank the National Center for CryoEM Access and Training (NCCAT) and the Cornell Center for Material Science (CCMR) for access to data collection facilities. We thank Yajie Xu and Dr Mariena Silvestry Ramos for assistance in CryoEM training and data collection. We thank the Proteomics and Metabolomics Facility of Cornell University for providing the mass spectrometry data and NIH SIG grant 1S10 OD017992-01 support for the Orbitrap Fusion mass spectrometer. The Center for High-Energy X-ray Sciences at CHESS (CHEXS) is supported by the NSF award DMR-2342336, and the MacCHESS resource is supported by NIGMS award 1-P30-GM124166. All figures were created or formated using Biorender.com.

## Funding

This work was supported by grant MCB-2129728 from the National Science Foundation to BRC.

## Author contributions

D.N. and B.R.C. designed research, conducted research, analyzed data and wrote the manuscript. Competing interests: The authors declare that they have no competing interests. Data and materials availability: The cryoEM data and structures are available at the Protein Data Bank with accession codes: 9Q05, 9Q06, 9Q0Y, 9Q0X, and 9Q15. All other data needed to evaluate the conclusions in the paper are present in the paper or the Supplementary Materials.

## Supplementary Movies

Movie S1: 3D Flexible Refinement of syNOS dimer shows a ‘breathing’ motion

## Supplementary Tables

Table S1: Description of datasets, conditions and refined classes.

Table S2: Identified and validated DSBU crosslinks

Table S3: Cryo-EM data collection, refinement, and validation statistics.

Table S4: DNA primers used in the study with the cloning approach.

Table S5: Reduction potentials (E^0^’) for syNOS_Glb_, syNOS_Fld_ and syNOS_FNR_ vs. the Normal Hydrogen Electrode (NHE)

## Supplementary Figures

Figure S1: syNOS purification

Figure S2: Dataset 1 processing scheme

Figure S3: Dataset 2 processing scheme

Figure S4: Merged data for NOS-Sym state, boxed class used for 3DVA and 3DFlex

Figure S5: Dataset 3 processing scheme and validation.

Figure S6: Dataset 4 processing scheme and validation

Figure S7: Dataset 5 processing scheme and validation

Figure S8: syNOS linker intertwining

Figure S9: syNOS tight and loose NOS_Oxy_ states

Figure S10: Identified cross-links plotted onto dimer interface Oxy+pGlb and the syNOS FNR+CTT

Figure S11: Phylogenetic sequence clustering in the context of syNOS_Red_.

Figure S12: C-terminal tail (CTT) sequence alignment of syNOS, mNOSs and diatom NOS.

Figure S13: Density validation of key interactions

Figure S14: SEC-SAXS of syNOS and variants.=

Figure S15: Steady state cytochrome c reductase activities of WT syNOS and variants

Figure S16: Flavin reduction by NADPH

Figure S17: Isothermal calorimetric titration of Ca^2+^ with syNOS_C2_

Figure S18: syNOS NOD activity and heme reduction rates

Figure S19: Output state predicted by AlphaFold3

## Supplemental Movies

**Movie S1.** 3D Variability analysis of one subunit of the syNOS asymmetric dimer showing the breathing motion of NOS_Red_ as a unit relative to motion about the NOS_Oxy_ interface.

## Supplementary Tables

**Table S1.**
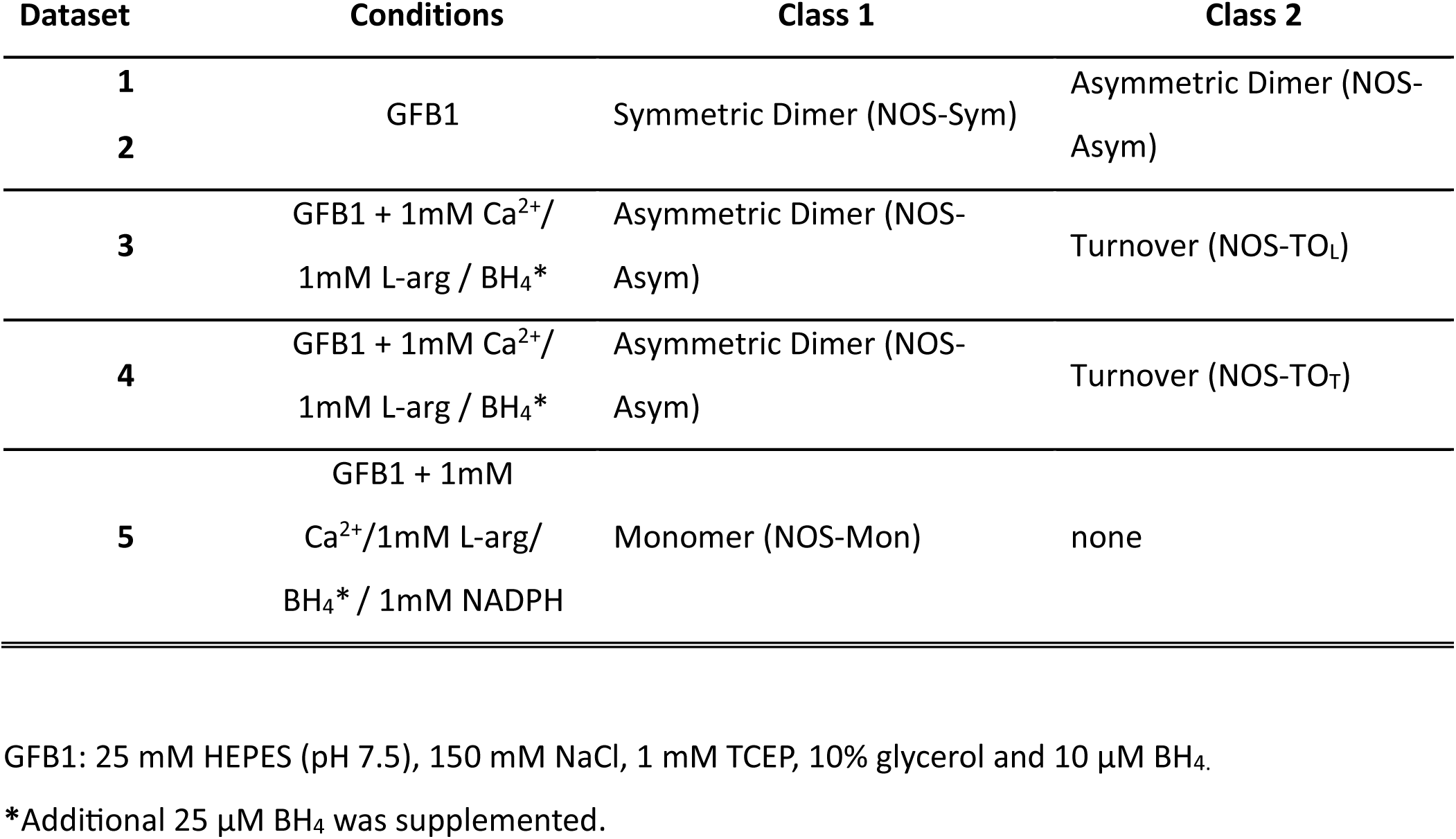
Description of datasets, conditions and refined classes.

**Table S2.**
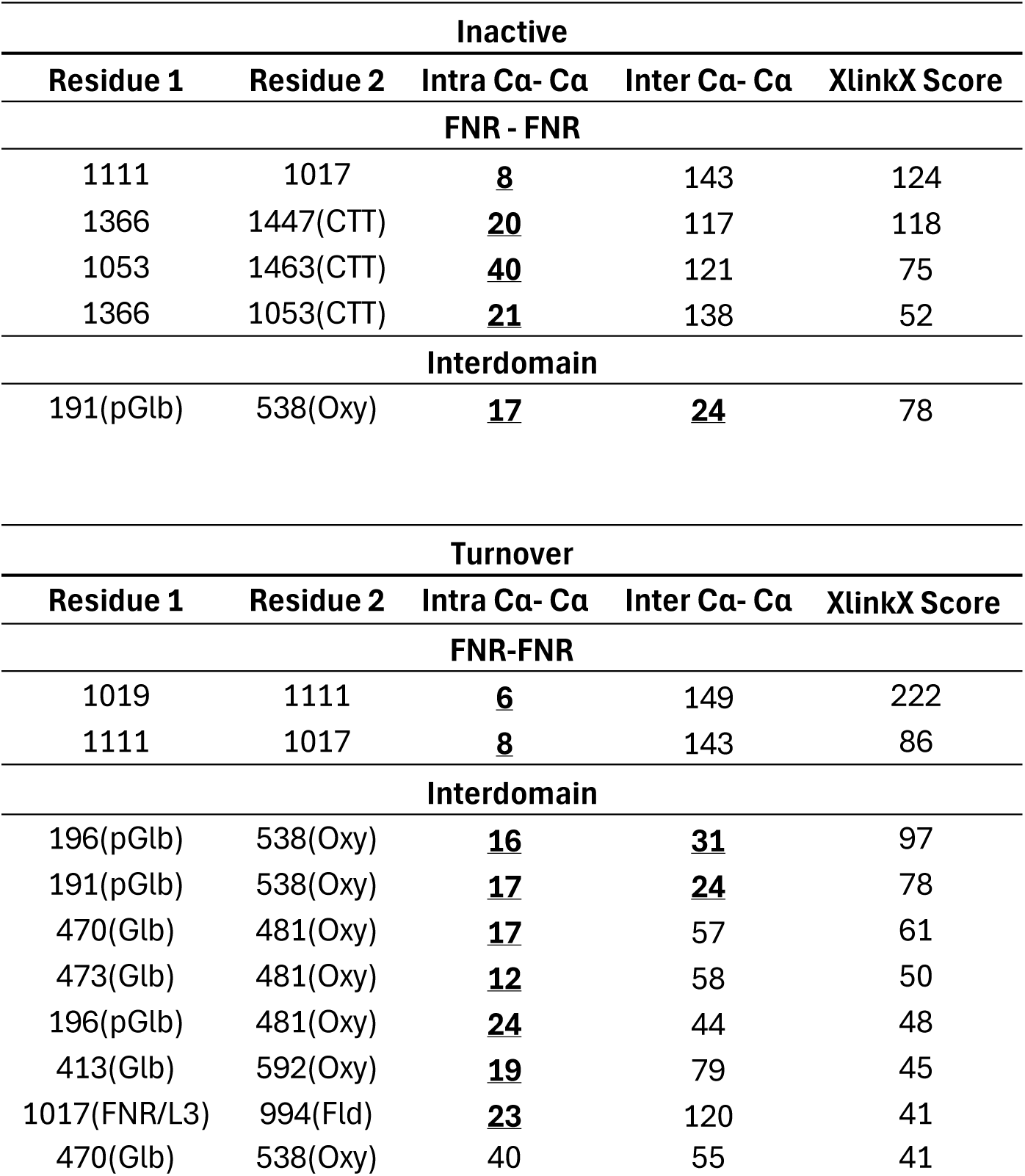
Identified and validated DSBU crosslinks. Reasonable distances based on the cryo-EM structures are bolded and underlined. PDB:9Q15 was used to estimate distances.

**Table S3.**
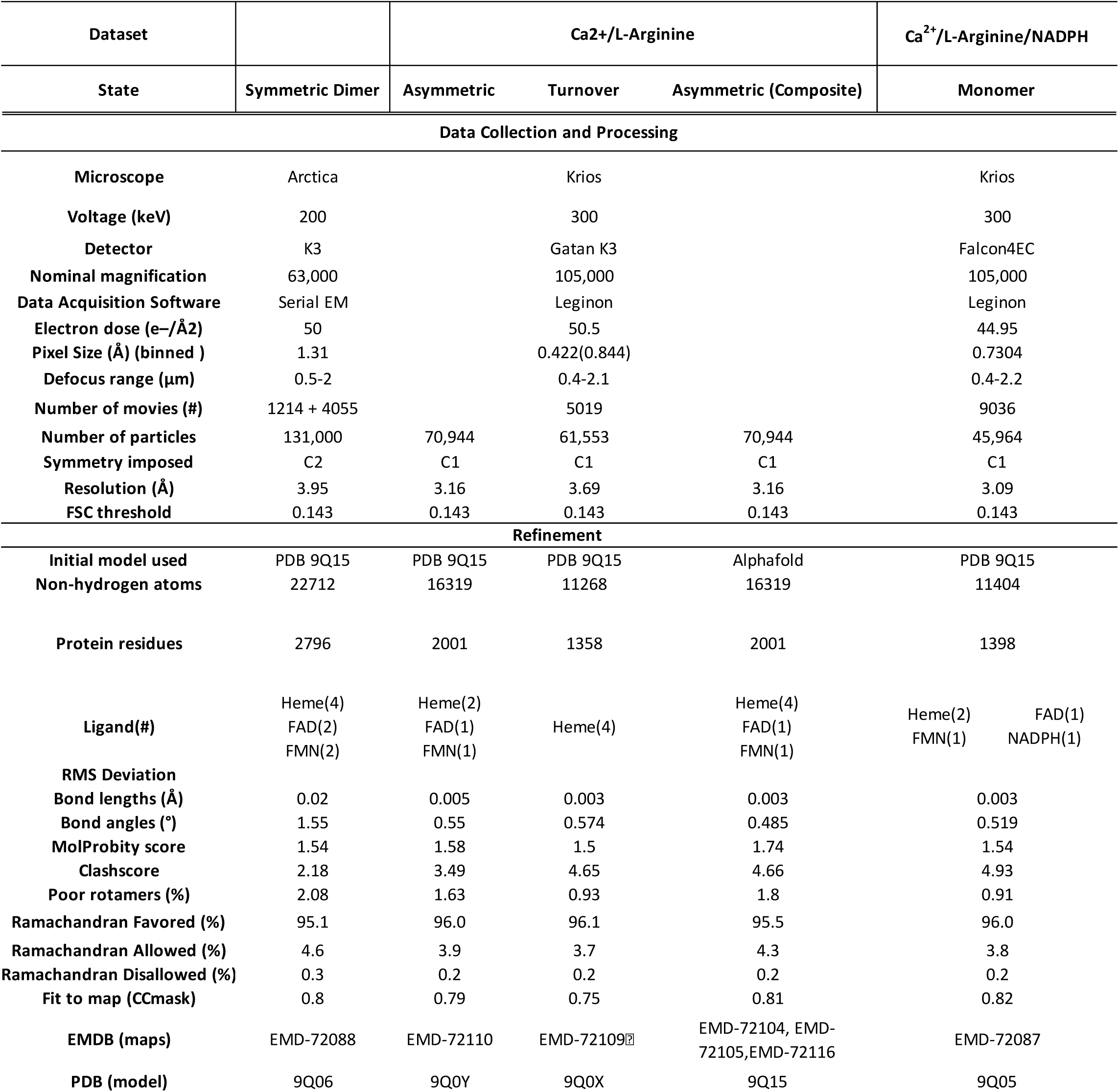
Cryo-EM data collection, refinement, and validation statistics.

**Table S4.**
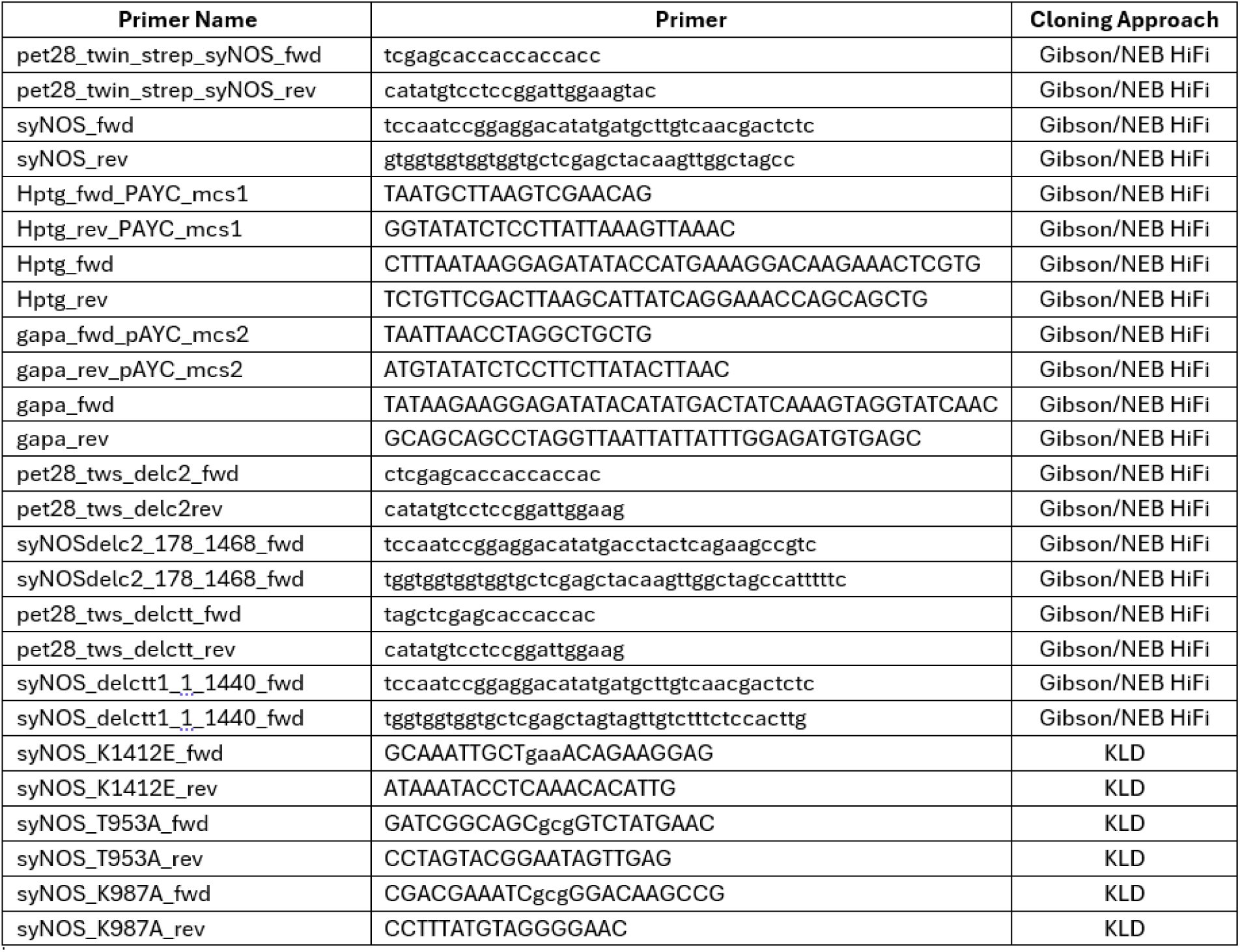
Primers used in the study with the cloning approach (left).

**Table S5.**
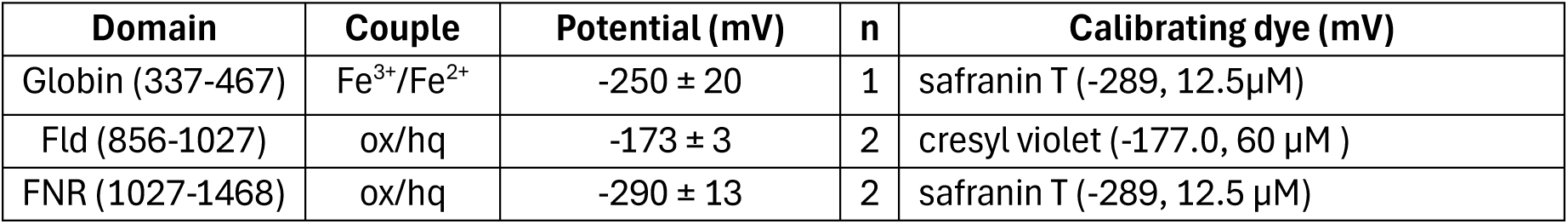
Reduction potentials (E^0’^) for syNOS_Glb_, syNOS_Fld_ and syNOS_FNR_ vs. the Normal Hydrogen Electrode (NHE). For syNOS_Glb_ the single electron reduction (n_e_=1) from Fe^3+^ to Fe^2+^ was measured whereas for the flavin-containing domains a midpoint potential for the 2-electron reduction (n_e_=2) from oxidized (ox) to hydroquinone (hq) was measured as no spectral signature for the semiquinone was apparent in the titrations. (n = 3 for all measurements)

## Supplementary Figures

**Figure S1.**
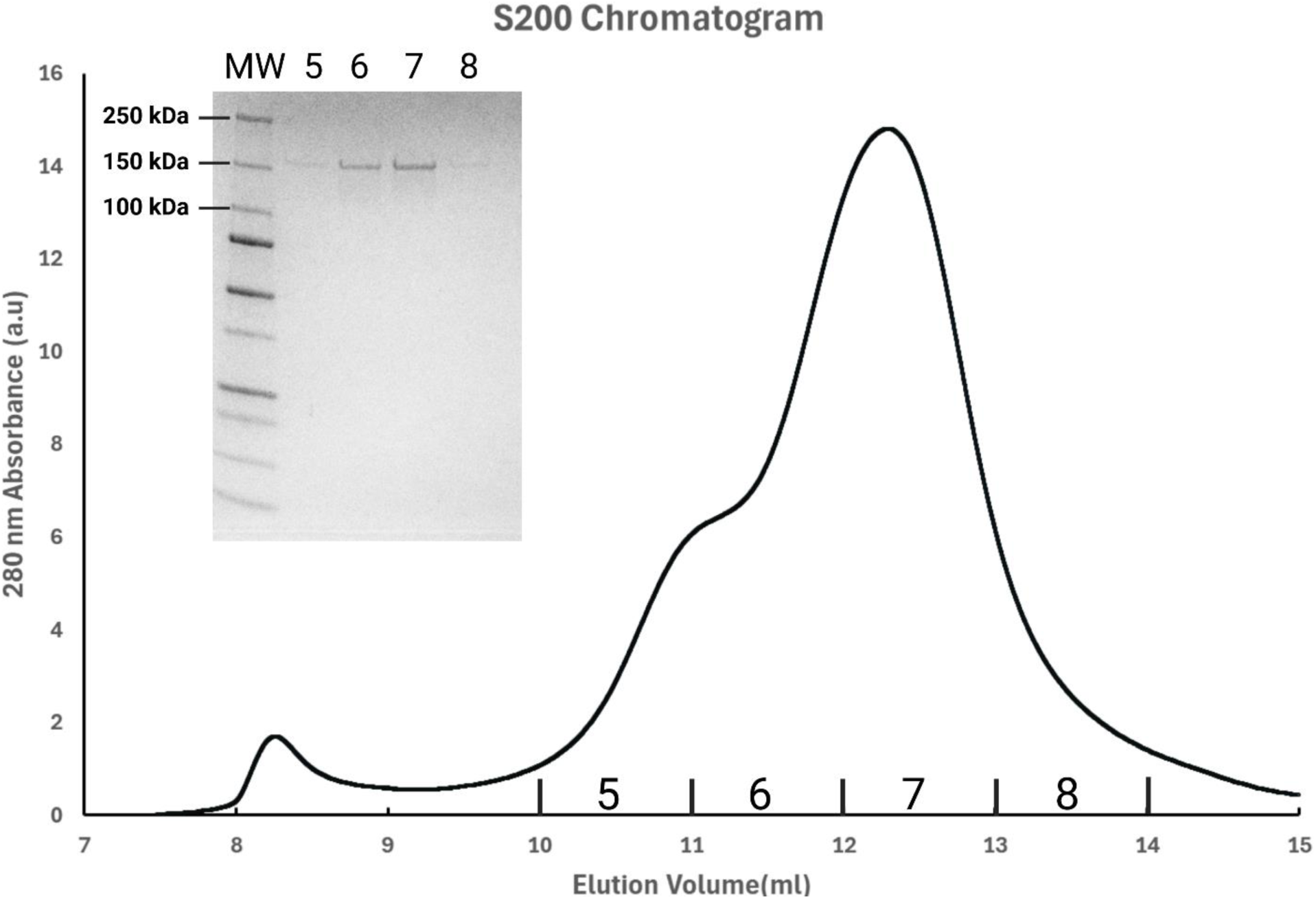
syNOS Purification. Representative chromatogram from a Superdex 200 increase 10/30 size exclusion chromatography column with an arrow denoting the syNOS dimer peak. The inset corresponds to a Coomassie blue stained 4-12% SDS-PAGE gel of fractions 5,6,7 and 8.

**Figure S2.**
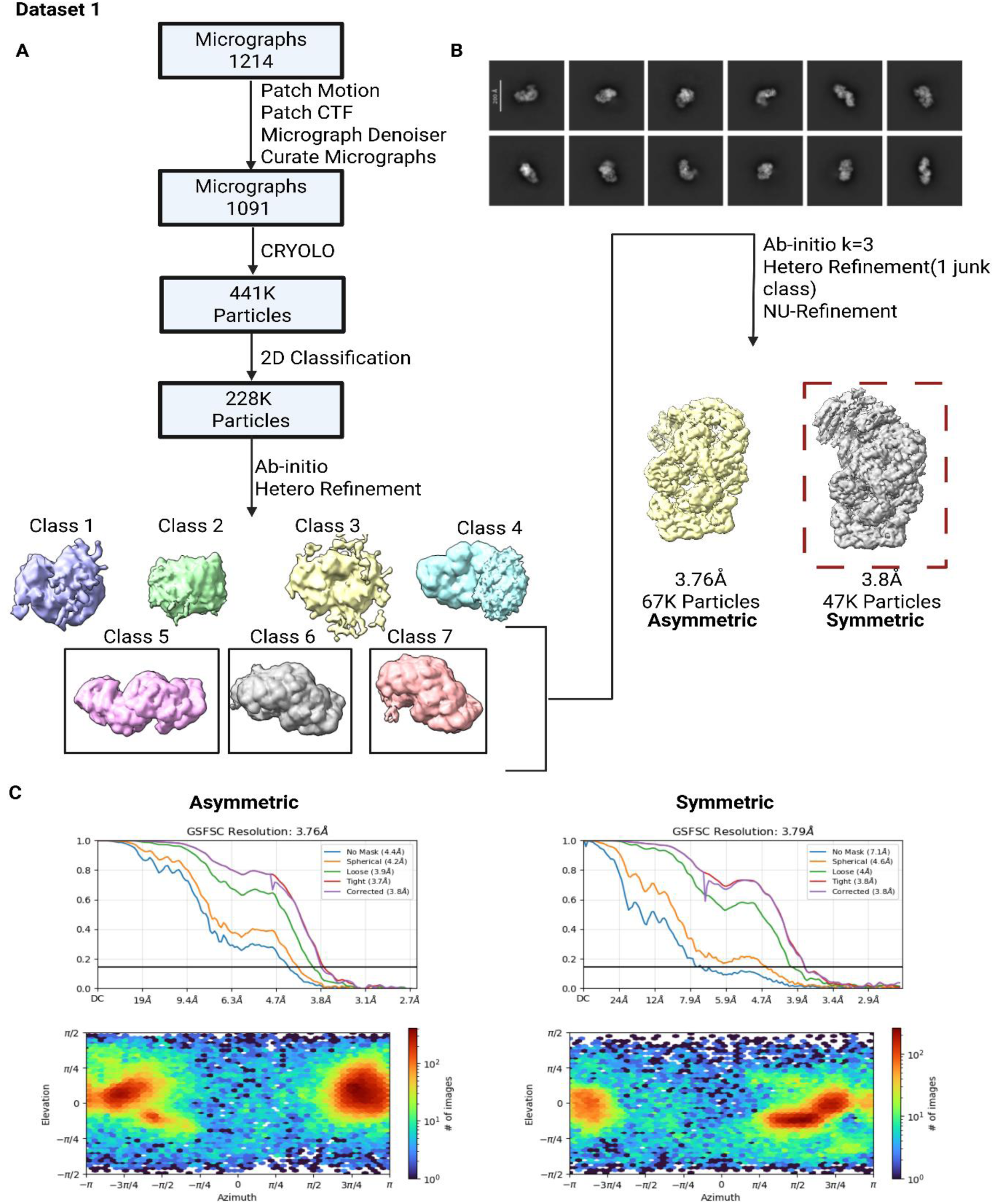
Dataset 1 processing scheme. (A) CryoEM processing workflow for dataset 1: syNOS ligand free. NOS-Sym class used for 3DVA and 3DFlex boxed (dashed-red). (B) Representative 2D Classes. CTF, contrast transfer function; NU, non-uniform refinement. (C) Overall Particle and Gold-standard Fourier Shell Correlation (GSFSC) curves for NOS-Asym and NOS-Sym.

**Figure S3.**
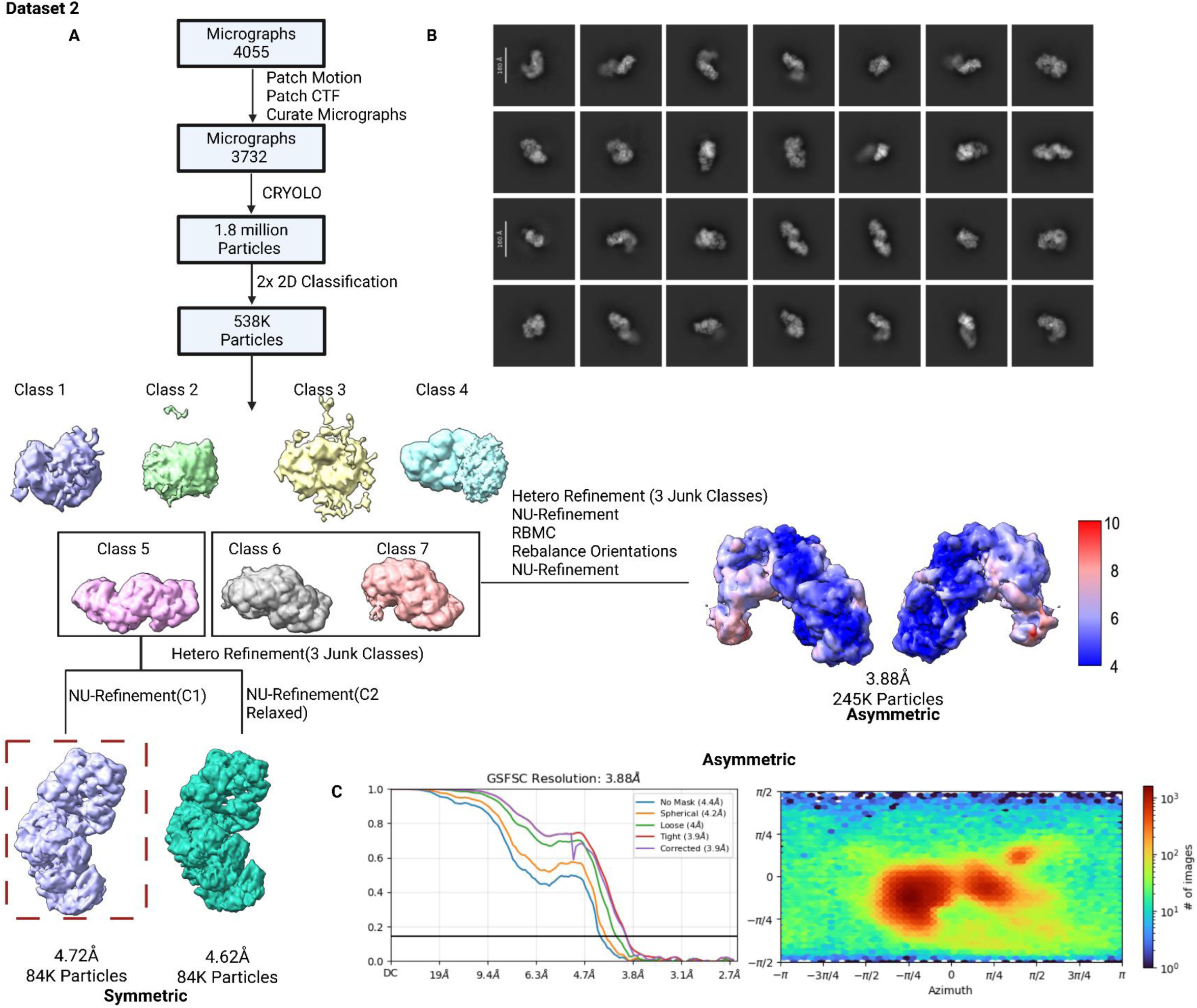
Dataset 2 processing scheme. (A) CryoEM processing workflow for dataset 2: syNOS ligand free. NOS-Sym class used for 3DVA and 3DFlex boxed (dashed-red). (B) Representative 2D Classes. (C) GSFSC curves and positional distribution of particles.

**Figure S4.**
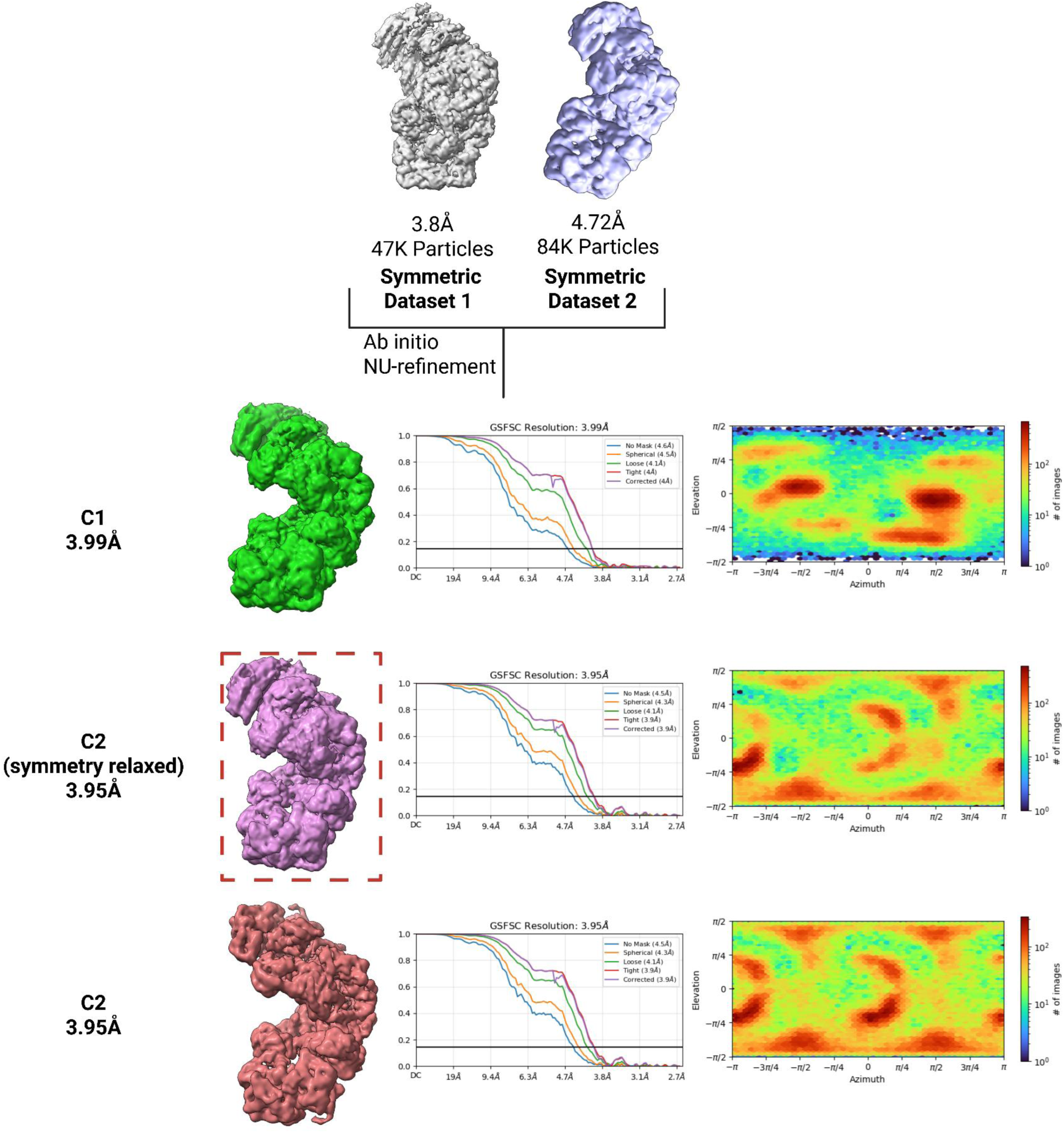
Merged data for NOS-Sym state, boxed class used for 3DVA and 3DFlex.

**Figure S5.**
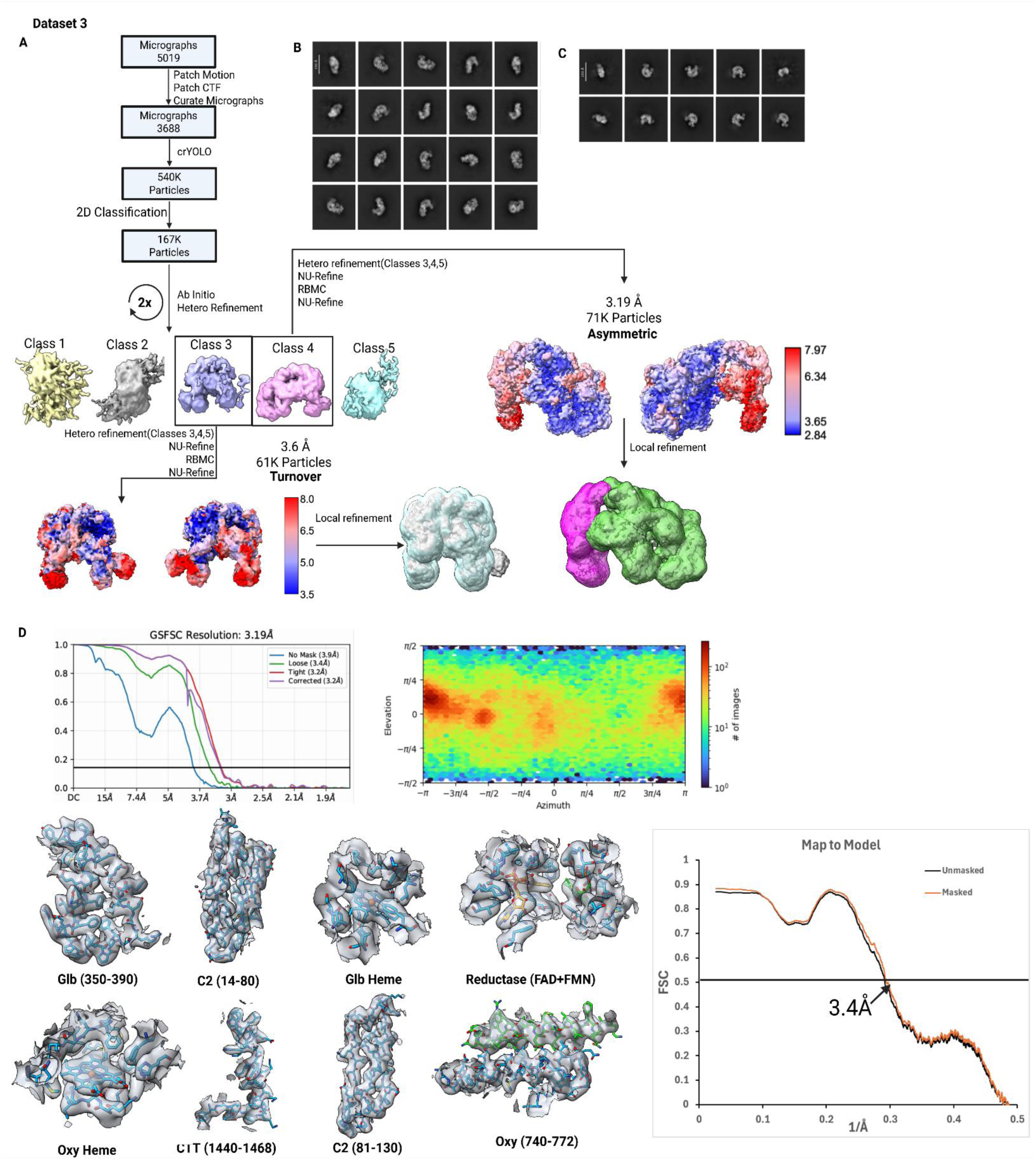
Dataset 3 processing scheme and validation. (A) CryoEM processing workflow for dataset 3: syNOS in the presence of Ca^2+^/L-arginine (B) Representative 2D Classes for NOS-Asym state (C) Representative 2D classes for NOS turnover state (NOS-TO_L_) (D) Asymmetric states GSFSC curves, orientation validation, model density fit and map to model Fourier Shell Correlation (FSC).

**Figure S6.**
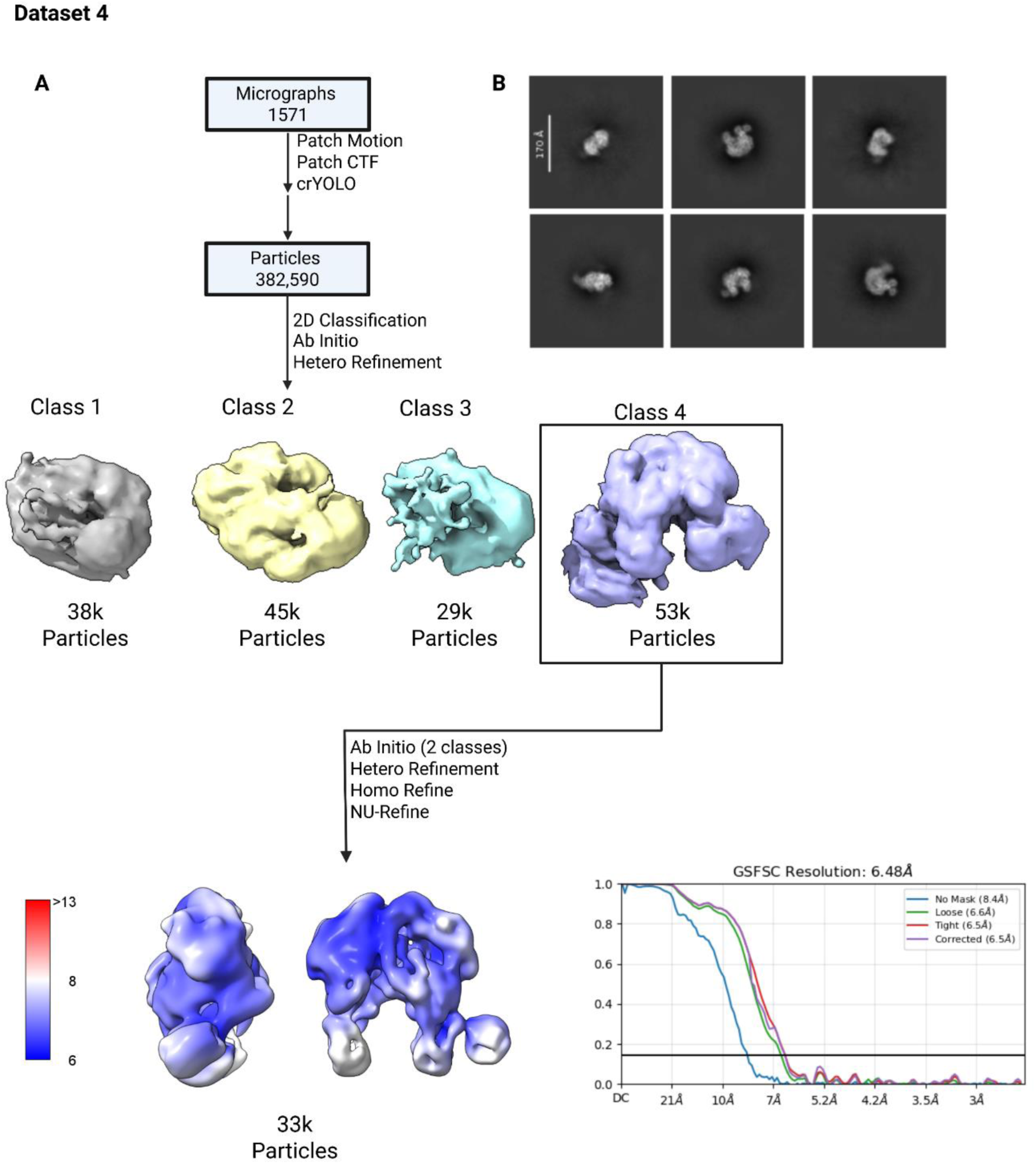
Dataset 4 processing scheme and validation. (A) CryoEM processing workflow for dataset 4, syNOS with NADPH/Ca^2+^ including GSFSC curves. (B) Representative 2D classes for NOS-TO_T_.

**Figure S7.**
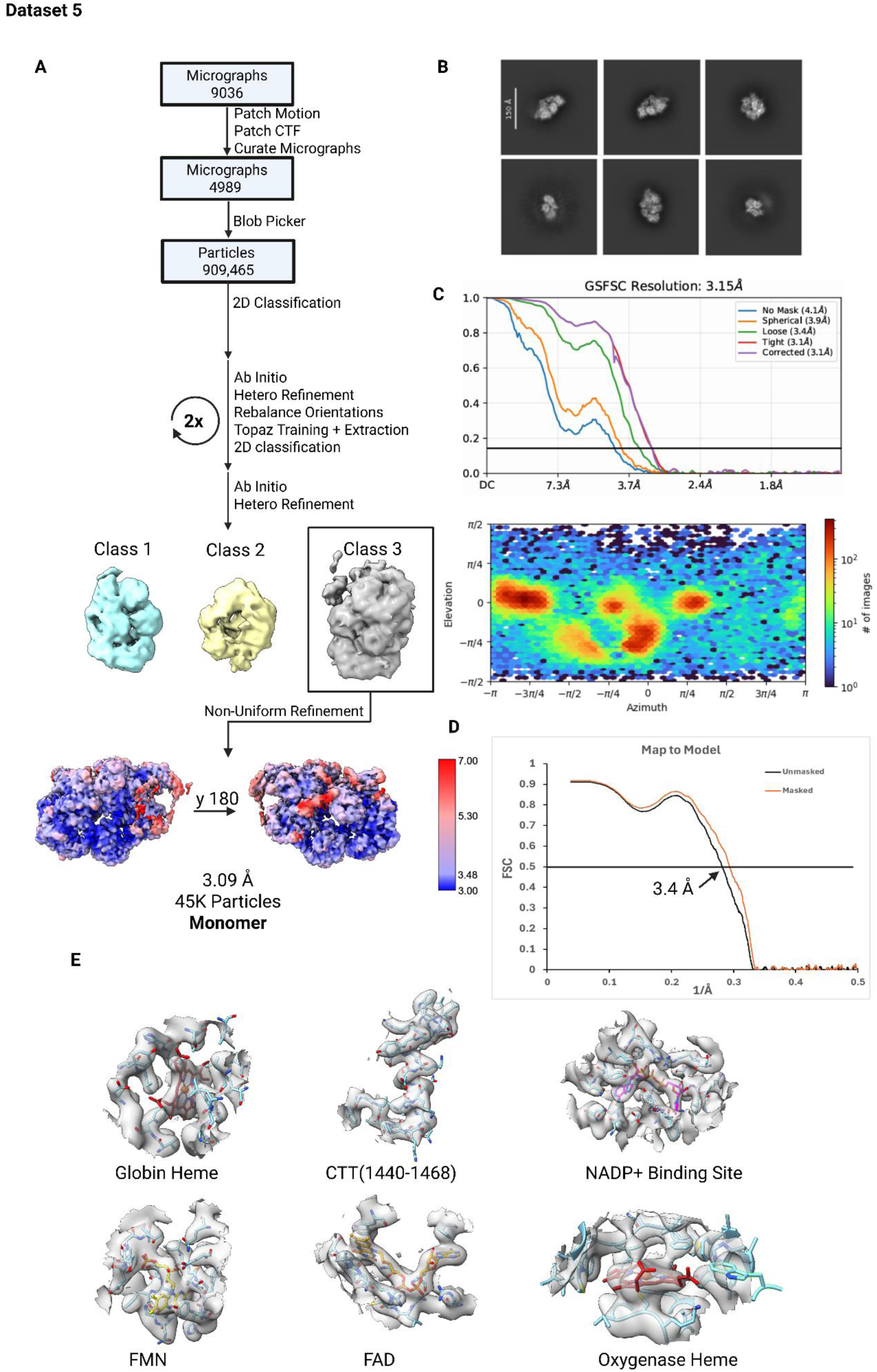
Dataset 5 processing scheme and validation. (A) CryoEM processing workflow for dataset 5: syNOS with L-arginine, NADPH/Ca^2+^/NADPH (B) Representative 2D classes for NOS-Mon. (C) GSFSC curves, orientation validation and (D) Map to Model FSC. (E) Density fit of model in map.

**Figure S8.**
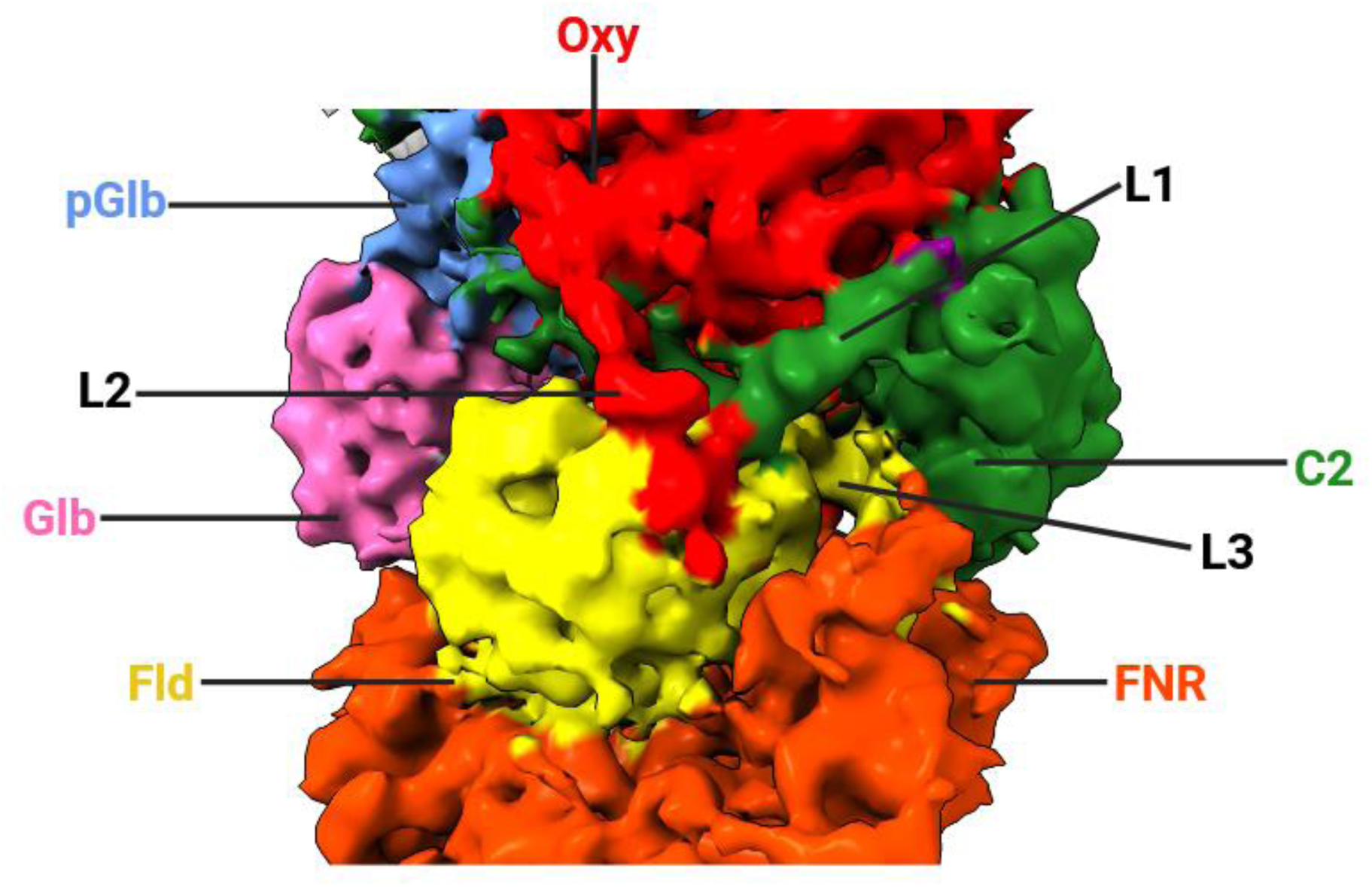
SyNOS linker intertwining. Intermediate resolution homodimer cryoEM density with all three linkers colored the same as their preceding domains. Note how L1 threads beneath L2 and on top of L3. Movement of the C2 domain (green) will release the Fld module for NOS_Oxy_ reduction.

**Figure S9.**
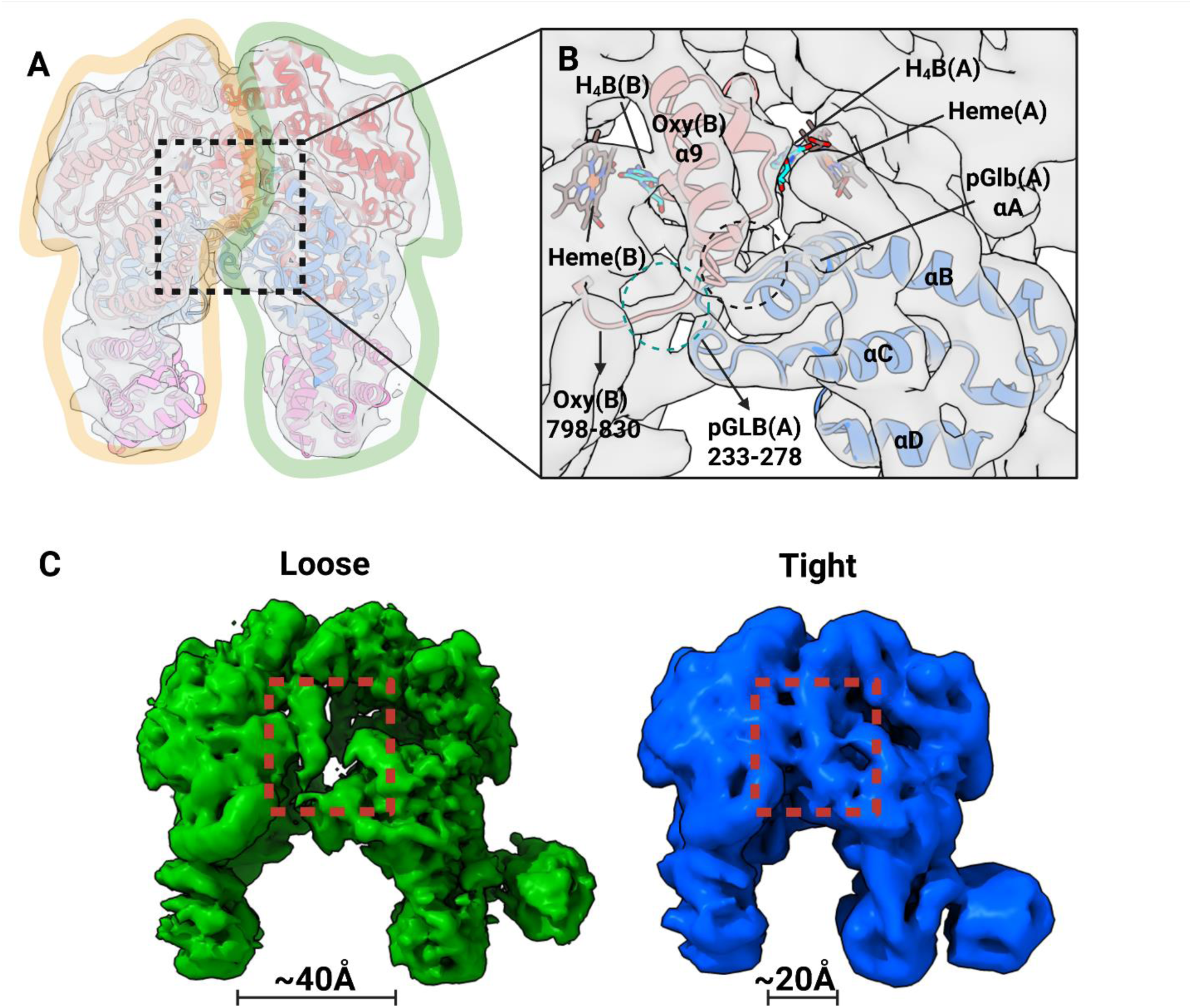
SyNOS tight and loose NOS_Oxy_ states. (A) syNOS cryoEM density (dataset 4) with the NOS_Oxy_ dimer interface outlined (orange and green) to show each subunit. (B) Close up of the dimeric interface showing the close association of NOS_Oxy_ α9 to an intersubunit pGlb C/D loop. Although at lower resolution, strong electron density for the core the NOS-TO_T_ interface includes the helical lariats and BH_4_. (C) Loose (green) and tight (blue) NOS_Oxy_ dimers differ by a large 20 Å breathing motion normal to the dimer interface that also affects the positioning of the C2 domain (lower right peripheral electron density).

**Figure S10.**
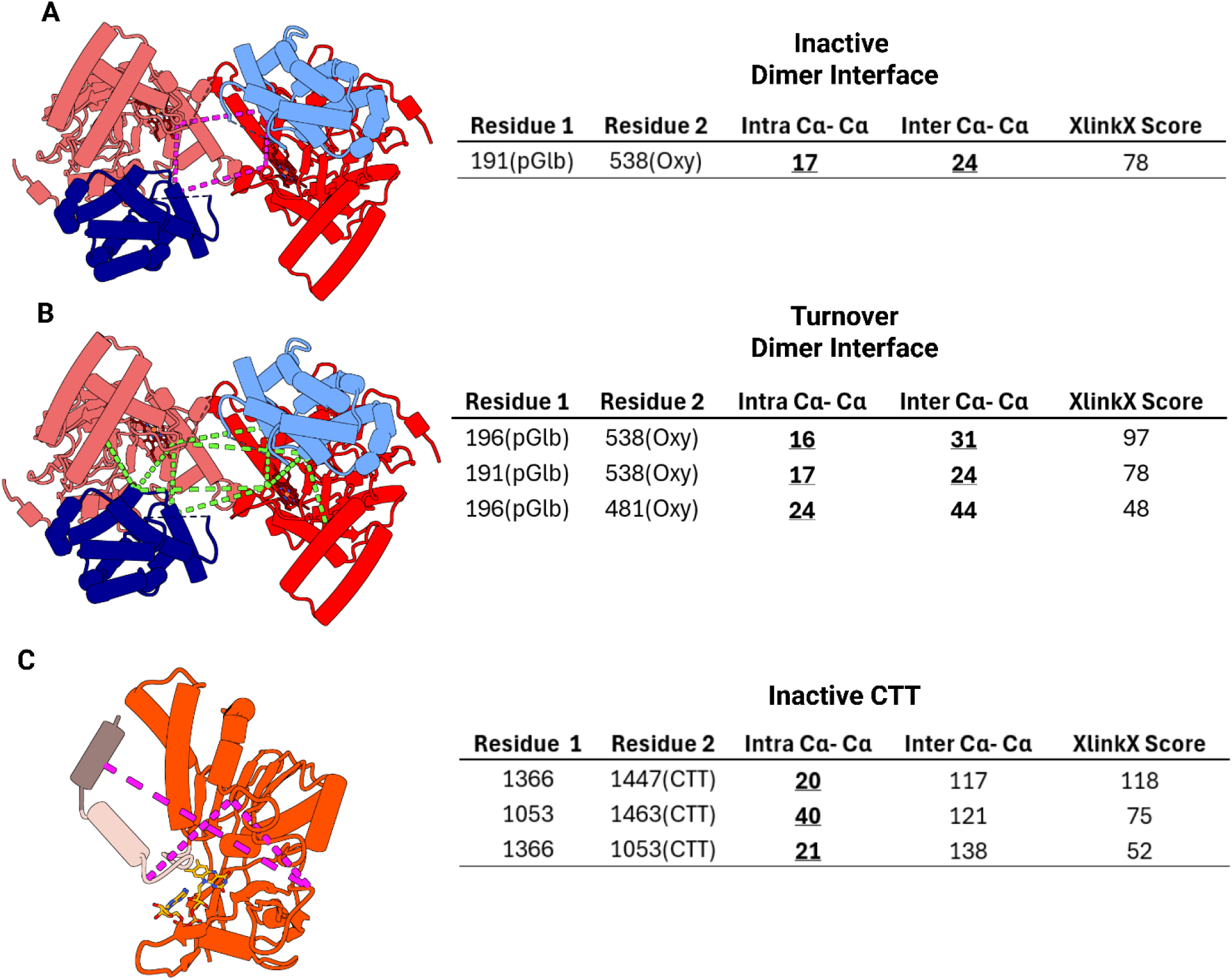
DSBU crosslinks plotted onto the dimer interface of syNOS Oxy + pGlb and the syNOS FNR + CTT. (A) syNOS inactive dimer interface, subunit 1 Oxy (red) + pGlb (blue) and subunit 2 Oxy (bright red) + pGlb (navy), with magenta crosslinks corresponding to table (left). (B) syNOS inactive dimer interface, protomer 1 Oxy (red) + pGlb (blue) and subunit 2 Oxy (bright red) + pGlb (navy), with green crosslinks corresponding to table (left). (C) syNOS inactive CTT with magenta crosslinks corresponding to table (left). Crosslinks within range based on the structures are underlined and bolded.

**Figure S11.**
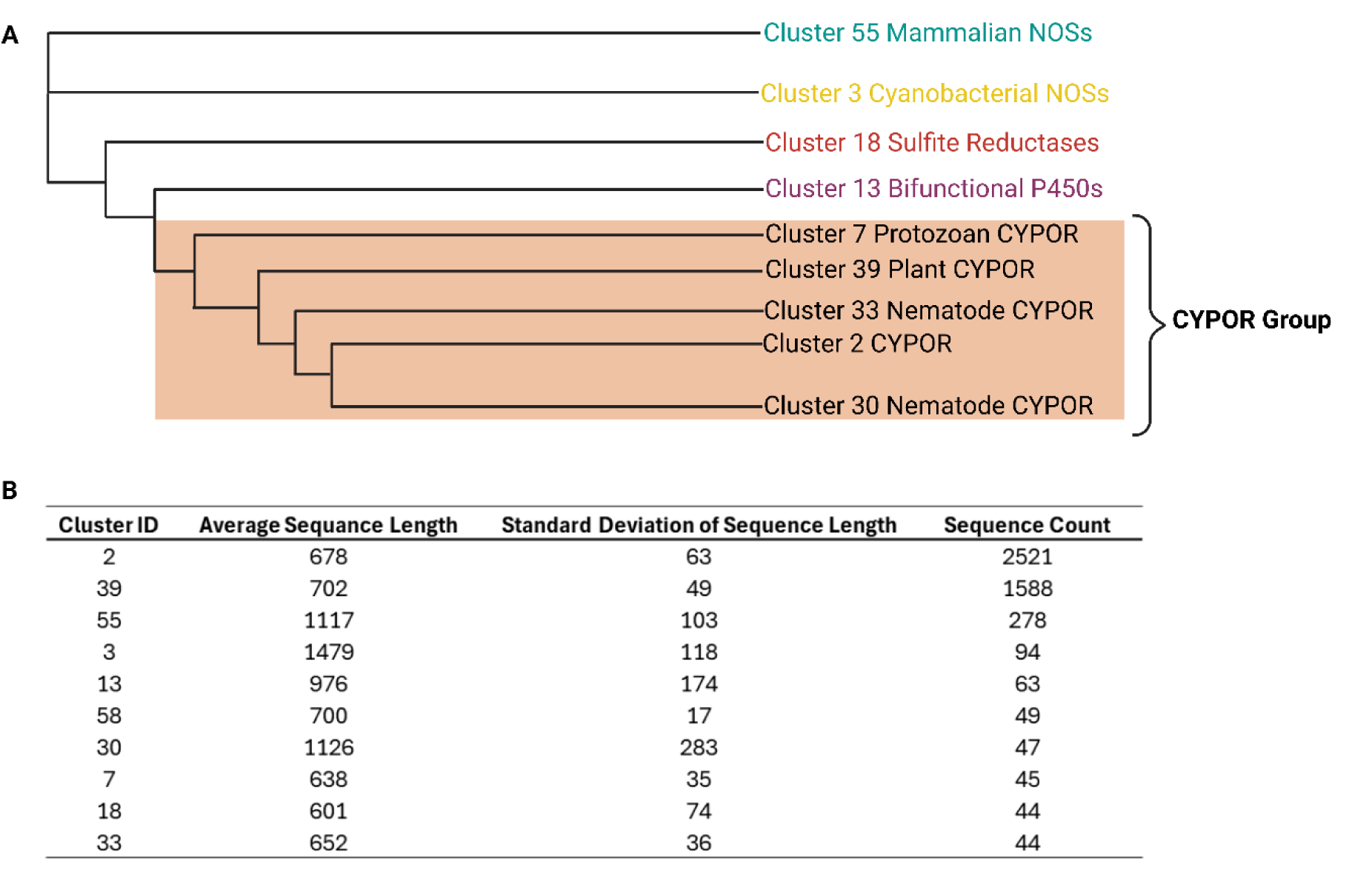
Phylogenetic sequence clustering in the context of syNOS_Red_(residues 853-1468). Representative clusters were subjected to PhyML with clusters above 40 sequences represented in the tree. (B) Cluster composition by sequence length and count.

**Figure S12.**
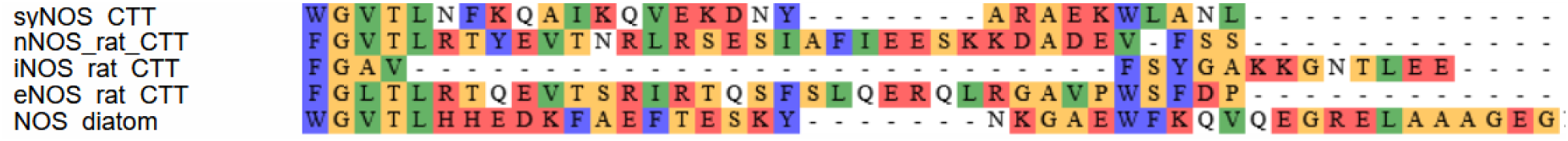
C-terminal tail (CTT) sequence alignment of syNOS, mNOSs and diatom NOS.

**Figure S13.**
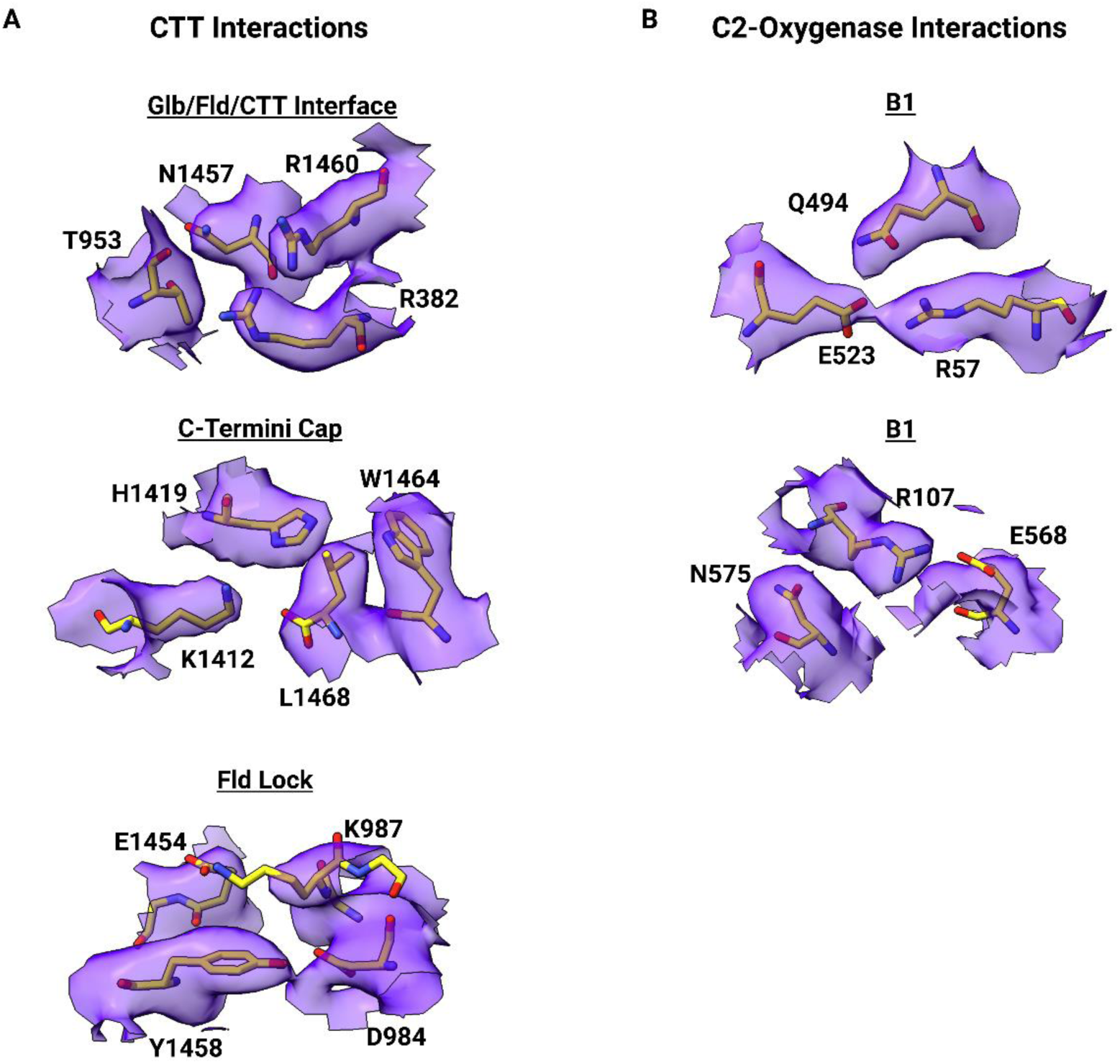
Cryo-EM density validation of key interactions. (A) Density fit of syNOS-Asym (9Q15), key interactions shown in Fig 3 C-E (B) Density fit of syNOS-Asym (9Q15), key interactions shown in Fig 4. B1-B2.

**Figure S14.**
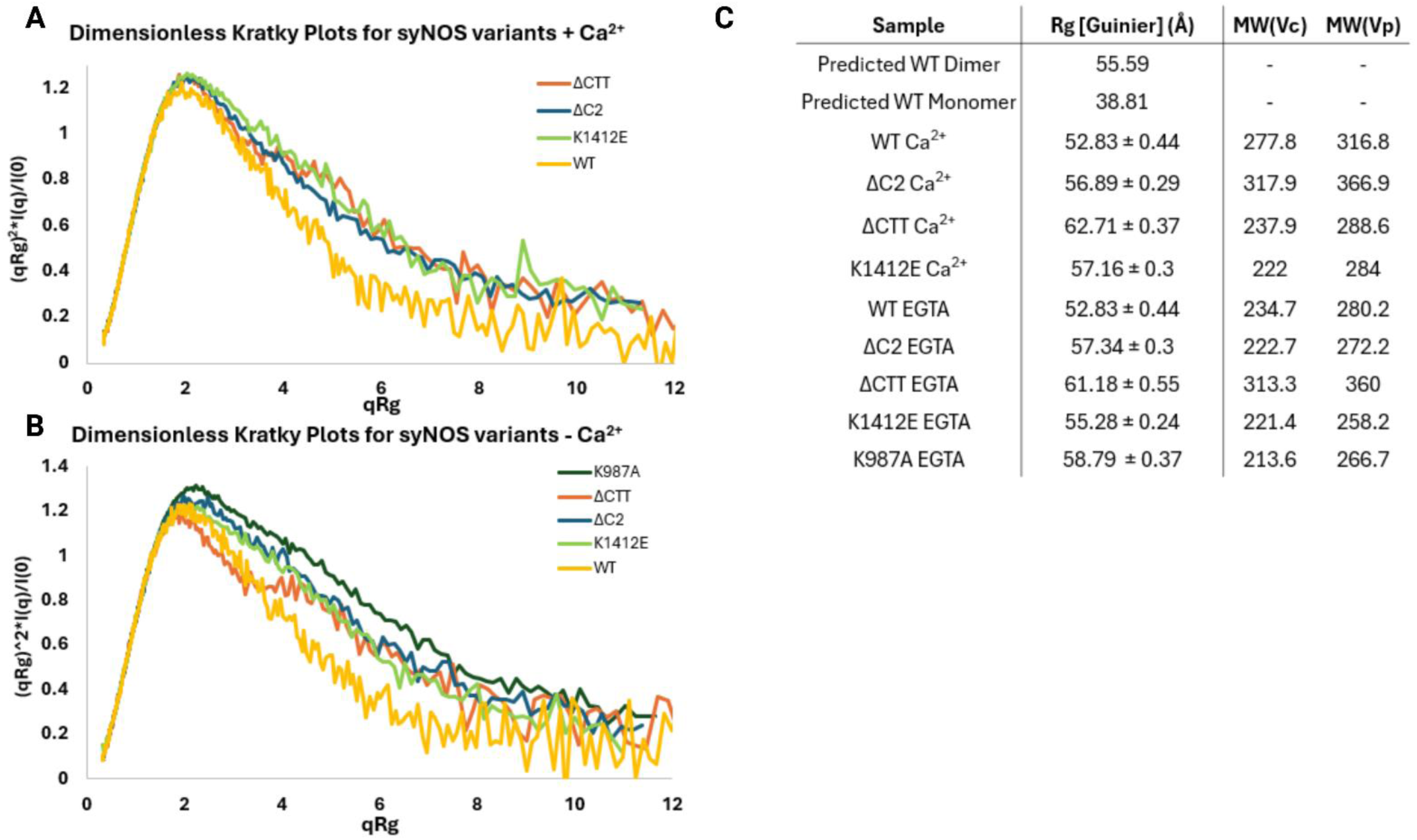
SEC-SAXS of syNOS and variants. (A) Dimensionless Kratky plots, binned by 4 qRg values, for samples in the presence and absence of Ca^2+^. (B) Table of predicted and experientially characterized radius of gyrations (Rgs) and molecular weights calculated either by volumes of correlation (V_C_) or Porod volumes (V_P_). (Samples were measured once)

**Figure S15.**
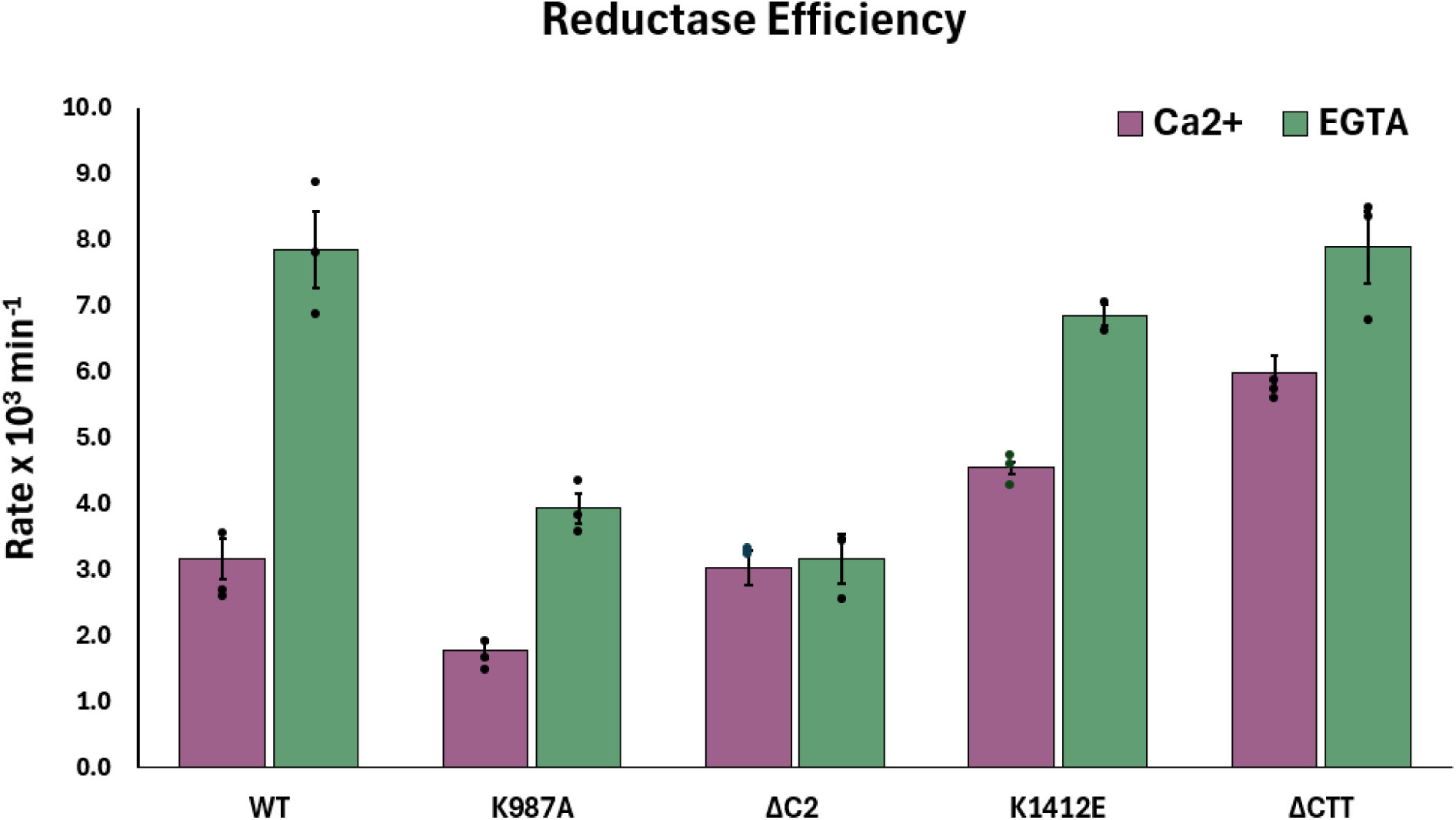
Steady state cytochrome c reductase activities of WT syNOS and variants expressed as turnover numbers. (Cc reduced (NOS subunit)^-1^ min^-1^) with standard errors shown as error bars (see Methods). Statistical comparisons between groups were performed with a 2-way ANOVA with post hoc analysis using Tukey’s multiple comparisons test. Significant differences are indicated as follows: ns (not significant), * (p ≤ 0.05), ** (p ≤ 0.01), *** (p ≤ 0.001), **** (p ≤ 0.0001)..Ca:WT vs. Ca:K987A (ns); Ca:WT vs. Ca:ΔC2 (ns); Ca:WT vs. Ca:K1412E (ns); Ca:WT vs. Ca:ΔCTT (ns); Ca:WT vs. EGTA:WT (****); Ca:WT vs. EGTA:K987A (ns); Ca:WT vs. EGTA:ΔC2 (ns); Ca:WT vs. EGTA:K1412E (****); Ca:WT vs. EGTA:ΔCTT (****);Ca:K987A vs. Ca:ΔC2 (ns); Ca:K987A vs. Ca:K1412E (***); Ca:K987A vs. Ca:ΔCTT (****); Ca:K987A vs. EGTA:WT (**); Ca:K987A vs. EGTA:K987A (**); Ca:K987A vs. EGTA:ΔC2 (ns); Ca:K987A vs. EGTA:K1412E (****); Ca:K987A vs. EGTA:ΔCTT (****); Ca:ΔC2 vs. Ca:K1412E (ns); Ca:ΔC2 vs. Ca:ΔCTT (**); Ca:ΔC2 vs. EGTA:WT (****); Ca:ΔC2 vs. EGTA:K987A (ns); Ca:ΔC2 vs. EGTA:ΔC2 (ns); Ca:ΔC2 vs. EGTA:K1412E (****); Ca:ΔC2 vs. EGTA:ΔCTT (****);Ca:K1412E vs. Ca:ΔCTT (ns); Ca:K1412E vs. EGTA:WT (****); Ca:K1412E vs. EGTA:K987A (ns); Ca:K1412E vs. EGTA:ΔC2 (ns); Ca:K1412E vs. EGTA:K1412E (**); Ca:K1412E vs. EGTA:ΔCTT (****); Ca:ΔCTT vs. EGTA:WT (**); Ca:ΔCTT vs. EGTA:K987A (*); Ca:ΔCTT vs. EGTA:ΔC2 (***); Ca:ΔCTT vs. EGTA:K1412E (ns); Ca:ΔCTT vs. EGTA:ΔCTT (**); EGTA:WT vs. EGTA:K987A (****); EGTA:WT vs. EGTA:ΔC2 (****); EGTA:WT vs. EGTA:K1412E (ns); EGTA:WT vs. EGTA:ΔCTT (ns); EGTA:K987A vs. EGTA:ΔC2 (ns); EGTA:K987A vs. EGTA:K1412E (***); EGTA:K987A vs. EGTA:ΔCTT (***); EGTA:ΔC2 vs. EGTA:K1412E (****); EGTA:ΔC2 vs. EGTA:ΔCTT (****); EGTA:K1412E vs. EGTA:ΔCTT (ns). (n=3, except for K1412E +EGTA; n = 2).

**Figure S16.**
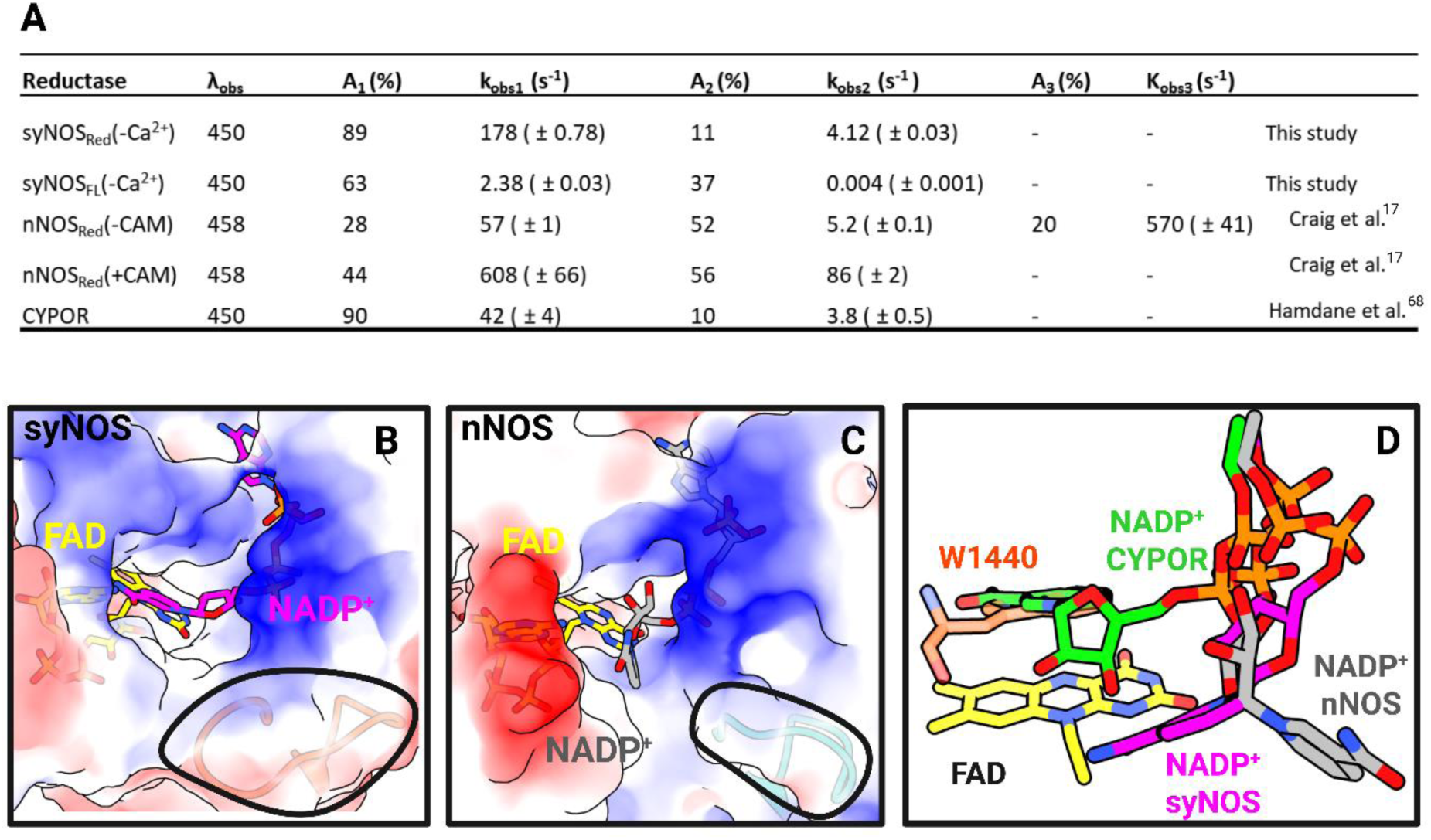
Flavin reduction by NADPH. (A) Table of flavin reduction rate constants for syNOS, nNOS and CYPOR(n = 4). (B) Surface representation and model of NADP^+^ (magenta) bound in the pocket of syNOS-Mon with loop 1050-1062 (orange-red and circled in black) in comparison to (C) nNOS bound to NADP^+^ (grey) with corresponding loop 1002-1012 (aqua) circled Iin black. (D) Overlay of NADP^+^ bound conformations of syNOS, nNOS and CYPOR W677X (1–676) (PDB: 1JA0) (green) over a consensus position of FNR-bound FAD (yellow). syNOS CTT residue 1440 blocks access of the nicotinamide ring to the isoalloxazine ring.

**Figure S17.**
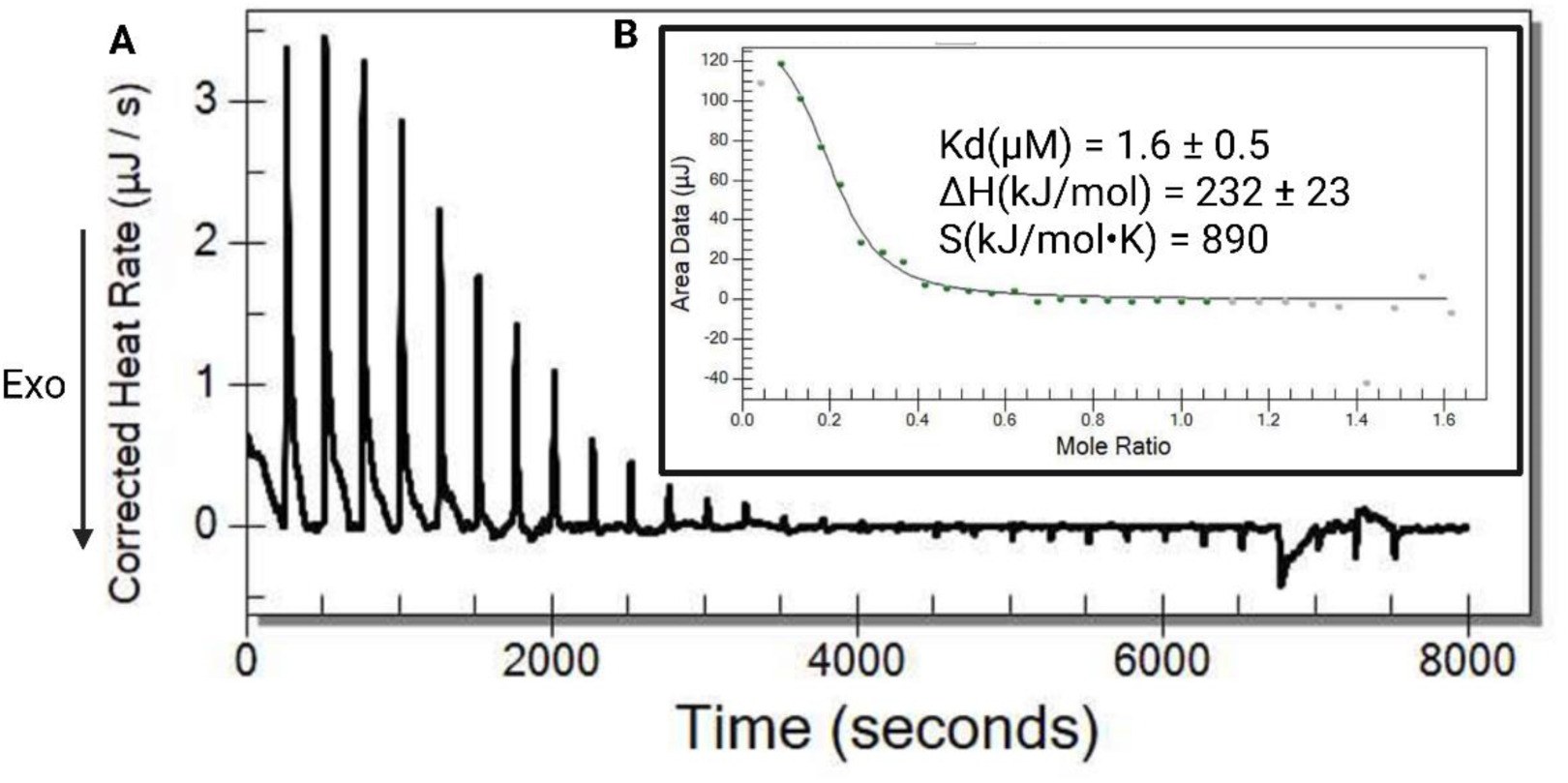
Isothermal calorimetric titration of Ca^2+^ with syNOS_C2_ (residues 1-138). (A) Endothermic, baseline corrected isotherm of Ca^2+^ titrated into syNOS_C2_. (B) Binding model fit with the accompanying thermodynamic parameters. (Samples were measured once)

**Figure S18.**
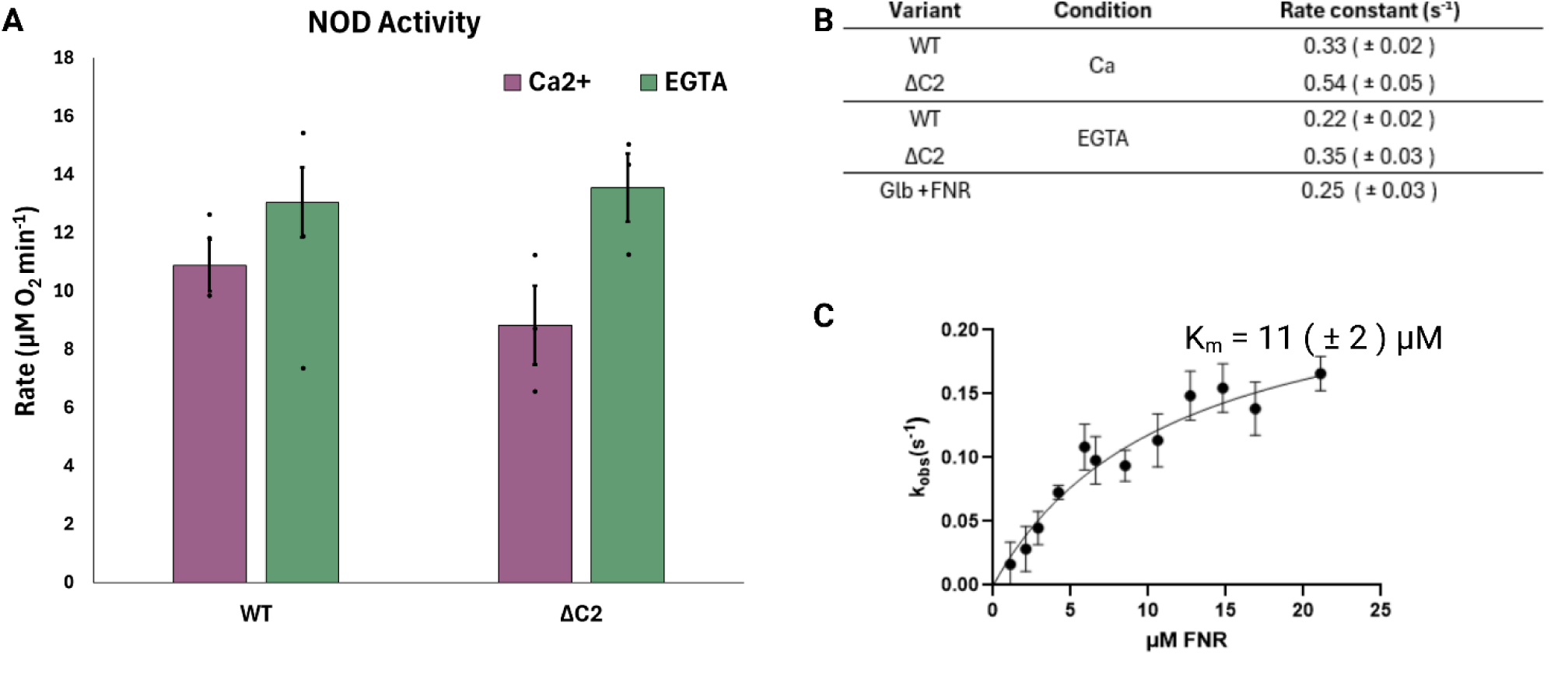
SyNOS globin-based NOD activity and heme reduction rates. (A) NOD activity was measured by O2 consumption in the presence of the NO-donor NOC-7 and NADPH for WT and syNOS_DC2_ (n=3, except for syNOS_DC2_ + Ca^2+^; n = 2). (B) Heme reduction rate constants of syNOS_Glb_ in the presence or absence of Ca^2+^ for WT or syNOS_DC2_ and the truncated syNOS_Glb_/syNOS_FNR_ couple in the absence of Ca^2+^, with standard error represented. Statistical comparisons between groups were performed with a 2-way ANOVA with post hoc analysis using Tukey’s multiple comparisons test, all comparisons were non-significant. (C) Michelis-Menten plot of truncated syNOS_Glb_ reduced by varied concentrations of truncated syNOS_FNR_ until saturation, with standard deviation represented. (Each data point reflects at least n = 3.)

**Figure S19.**
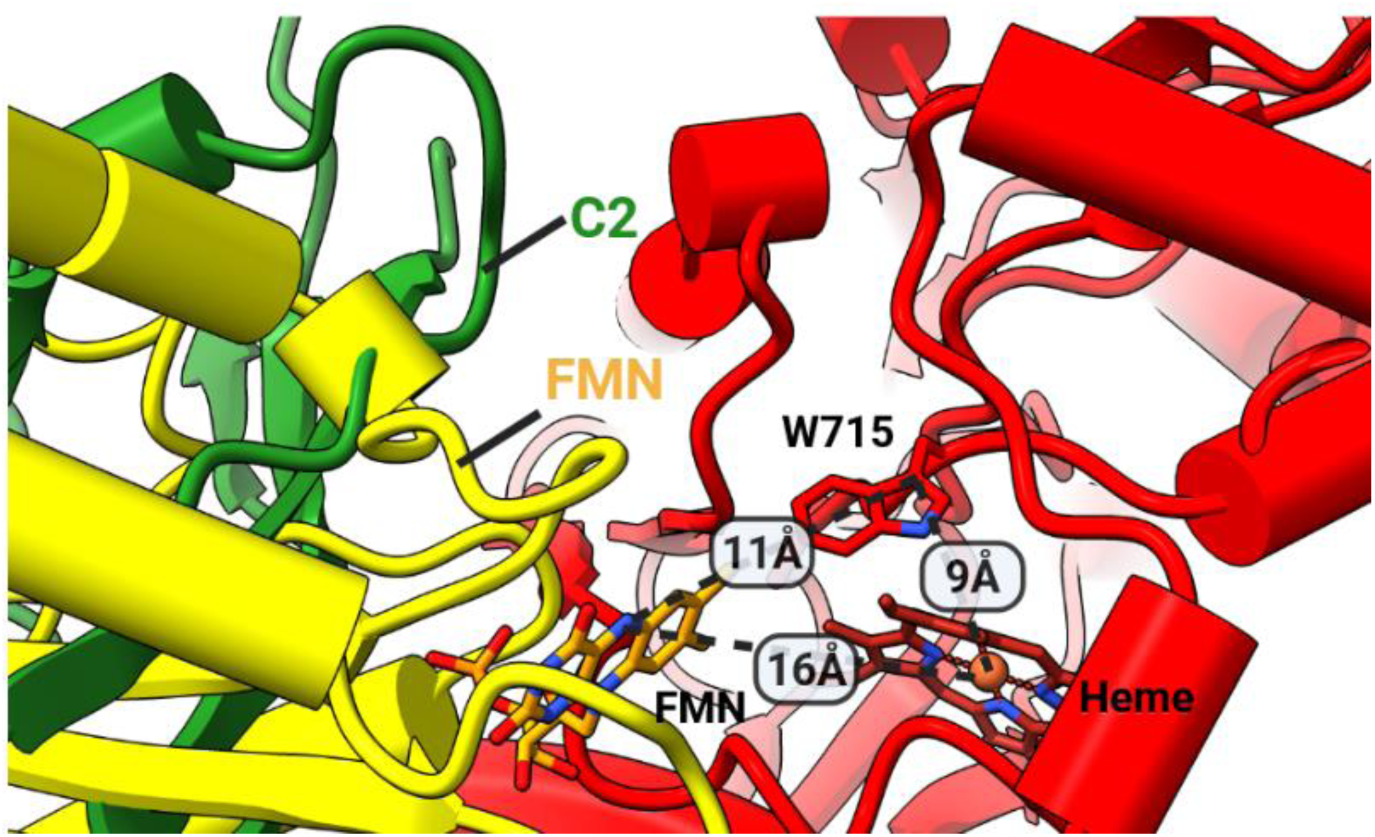
Output state predicted by AlphaFold3 with FMN-to-Trp715-to-Heme distances shown. Overlay of the C2 in the locked state (green) with the output Fld conformation (yellow) illustrates that the Fld is sterically blocked by the C2 domain in the NOS-Asym state. The AF3 prediction was made with a minimal unit of two truncated syNOS chains: 1) residues 473-863 (Oxy) and 2) residues 473-1003 (Oxy + Fld). Including additional domains did not predict interaction between syNOS_Fld_ and syNOS_Oxy_.

## Notes

### Competing Interest Statement

The authors have declared no competing interest.

### Summary of Updates

Revisions to suit peer review. Additional experiments were conducted and portions of the text was cleaned up.

