## Supplemental Tables and Figures for "Structure and dynamics of a muti-domain nitric oxide synthase regulated by a C2 domain"

**Supplementary Movies**

Movie S1: 3D Flexible Refinement of syNOS dimer shows a 'breathing' motion

**Supplementary Tables**

Table S1: Description of datasets, conditions and refined classes.

Table S2: Identified and validated DSBU crosslinks

Table S3: Cryo-EM data collection, refinement, and validation statistics.

Table S4: DNA primers used in the study with the cloning approach.

Table S5: Reduction potentials ($E^{0\prime}$) for $syNOS_{Glb}$, $syNOS_{Fld}$ and $syNOS_{FNR}$ vs. the Normal Hydrogen Electrode (NHE)

**Supplementary Figures**

Figure S1: syNOS purification

Figure S2: Dataset 1 processing scheme

Figure S3: Dataset 2 processing scheme

Figure S4: Merged data for NOS-Sym state, boxed class used for 3DVA and 3DFlex

Figure S5: Dataset 3 processing scheme and validation.

Figure S6: Dataset 4 processing scheme and validation

Figure S7: Dataset 5 processing scheme and validation

Figure S8: syNOS linker intertwining

Figure S9: syNOS tight and loose $NOS_{Oxy}$ states

Figure S10: Identified cross-links plotted onto dimer interface Oxy+pGlb and the syNOS FNR+CTT

Figure S11: Phylogenetic sequence clustering in the context of $syNOS_{Red}$.

Figure S12: C-terminal tail (CTT) sequence alignment of syNOS, mNOSs and diatom NOS.

Figure S13: Density validation of key interactions

Figure S14: SEC-SAXS of syNOS and variants.=

Figure S15: Steady state cytochrome c reductase activities of WT syNOS and variants

Figure S16: Flavin reduction by NADPH

Figure S17: Isothermal calorimetric titration of $Ca^{2+}$ with $syNOS_{C2}$

Figure S18: syNOS NOD activity and heme reduction rates
Figure S19: Output state predicted by AlphaFold3

#### Supplemental Movies

**Movie S1.** 3D Variability analysis of one subunit of the syNOS asymmetric dimer showing the breathing motion of $NOS_{Red}$ as a unit relative to motion about the $NOS_{Oxy}$ interface.

#### Supplementary Tables

| Dataset | Conditions | Class 1 | Class 2 |
|---|---|---|---|
| **1**<br>**2** | GFB1 | Symmetric Dimer (NOS-Sym) | Asymmetric Dimer (NOS-Asym) |
| **3** | GFB1 + 1mM $Ca^{2+}$/ 1mM L-arg / $BH_4$* | Asymmetric Dimer (NOS-Asym) | Turnover (NOS-$TO_L$) |
| **4** | GFB1 + 1mM $Ca^{2+}$/ 1mM L-arg / $BH_4$* | Asymmetric Dimer (NOS-Asym) | Turnover (NOS-$TO_T$) |
| **5** | GFB1 + 1mM $Ca^{2+}$/1mM L-arg/ $BH_4$* / 1mM NADPH | Monomer (NOS-Mon) | none |

GFB1: 25 mM HEPES (pH 7.5), 150 mM NaCl, 1 mM TCEP, 10% glycerol and 10 µM $BH_4$.

***Additional 25 µM $BH_4$ was supplemented.

**Table S1.** Description of datasets, conditions and refined classes.

| Inactive | | | | |
|---|---|---|---|---|
| **Residue 1** | **Residue 2** | **Intra Cα- Cα** | **Inter Cα- Cα** | **XlinkX Score** |
| | | **FNR - FNR** | | |
| 1111 | 1017 | **<u>8</u>** | 143 | 124 |
| 1366 | 1447(CTT) | **<u>20</u>** | 117 | 118 |
| 1053 | 1463(CTT) | **<u>40</u>** | 121 | 75 |
| 1366 | 1053(CTT) | **<u>21</u>** | 138 | 52 |
| | | **Interdomain** | | |
| 191(pGlb) | 538(Oxy) | **<u>17</u>** | **<u>24</u>** | 78 |

| Turnover | | | | |
|---|---|---|---|---|
| **Residue 1** | **Residue 2** | **Intra Cα- Cα** | **Inter Cα- Cα** | **XlinkX Score** |
| | | **FNR-FNR** | | |
| 1019 | 1111 | **<u>6</u>** | 149 | 222 |
| 1111 | 1017 | **<u>8</u>** | 143 | 86 |
| | | **Interdomain** | | |
| 196(pGlb) | 538(Oxy) | **<u>16</u>** | **<u>31</u>** | 97 |
| 191(pGlb) | 538(Oxy) | **<u>17</u>** | **<u>24</u>** | 78 |
| 470(Glb) | 481(Oxy) | **<u>17</u>** | 57 | 61 |
| 473(Glb) | 481(Oxy) | **<u>12</u>** | 58 | 50 |
| 196(pGlb) | 481(Oxy) | **<u>24</u>** | 44 | 48 |
| 413(Glb) | 592(Oxy) | **<u>19</u>** | 79 | 45 |
| 1017(FNR/L3) | 994(Fld) | **<u>23</u>** | 120 | 41 |
| 470(Glb) | 538(Oxy) | 40 | 55 | 41 |

**Table S2.** Identified and validated DSBU crosslinks. Reasonable distances based on the cryo-EM structures are bolded and underlined. PDB:9Q15 was used to estimate distances.

| Dataset | | Ca2+/L-Arginine | | | $Ca^{2+}$/L-Arginine/NADPH |
|---|---|---|---|---|---|
| **State** | **Symmetric Dimer** | **Asymmetric** | **Turnover** | **Asymmetric (Composite)** | **Monomer** |
| | | | **Data Collection and Processing** | | |
| **Microscope** | Arctica | | Krios | | Krios |
| **Voltage (keV)** | 200 | | 300 | | 300 |
| **Detector** | K3 | | Gatan K3 | | Falcon4EC |
| **Nominal magnification** | 63,000 | | 105,000 | | 105,000 |
| **Data Acquisition Software** | Serial EM | | Leginon | | Leginon |
| **Electron dose (e–/Å2)** | 50 | | 50.5 | | 44.95 |
| **Pixel Size (Å) (binned )** | 1.31 | | 0.422(0.844) | | 0.7304 |
| **Defocus range (µm)** | 0.5-2 | | 0.4-2.1 | | 0.4-2.2 |
| **Number of movies (#)** | 1214 + 4055 | | 5019 | | 9036 |
| **Number of particles** | 131,000 | 70,944 | 61,553 | 70,944 | 45,964 |
| **Symmetry imposed** | C2 | C1 | C1 | C1 | C1 |
| **Resolution (Å)** | 3.95 | 3.16 | 3.69 | 3.16 | 3.09 |
| **FSC threshold** | 0.143 | 0.143 | 0.143 | 0.143 | 0.143 |
| | | | **Refinement** | | |
| **Initial model used** | PDB 9Q15 | PDB 9Q15 | PDB 9Q15 | Alphafold | PDB 9Q15 |
| **Non-hydrogen atoms** | 22712 | 16319 | 11268 | 16319 | 11404 |
| **Protein residues** | 2796 | 2001 | 1358 | 2001 | 1398 |
| **Ligand(#)** | Heme(4) FAD(2) FMN(2) | Heme(2) FAD(1) FMN(1) | Heme(4) | Heme(4) FAD(1) FMN(1) | Heme(2) FAD(1) FMN(1) NADPH(1) |
| **RMS Deviation** | | | | | |
| **Bond lengths (Å)** | 0.02 | 0.005 | 0.003 | 0.003 | 0.003 |
| **Bond angles (°)** | 1.55 | 0.55 | 0.574 | 0.485 | 0.519 |
| **MolProbity score** | 1.54 | 1.58 | 1.5 | 1.74 | 1.54 |
| **Clashscore** | 2.18 | 3.49 | 4.65 | 4.66 | 4.93 |
| **Poor rotamers (%)** | 2.08 | 1.63 | 0.93 | 1.8 | 0.91 |
| **Ramachandran Favored (%)** | 95.1 | 96.0 | 96.1 | 95.5 | 96.0 |
| **Ramachandran Allowed (%)** | 4.6 | 3.9 | 3.7 | 4.3 | 3.8 |
| **Ramachandran Disallowed (%)** | 0.3 | 0.2 | 0.2 | 0.2 | 0.2 |
| **Fit to map (CCmask)** | 0.8 | 0.79 | 0.75 | 0.81 | 0.82 |
| **EMDB (maps)** | EMD-72088 | EMD-72110 | EMD-72109 | EMD-72104, EMD-72105,EMD-72116 | EMD-72087 |
| **PDB (model)** | 9Q06 | 9Q0Y | 9Q0X | 9Q15 | 9Q05 |

**Table S3.** Cryo-EM data collection, refinement, and validation statistics.

| Primer Name | Primer | Cloning Approach |
|---|---|---|
| pet28_twin_strep_syNOS_fwd | tcgagcaccaccaccacc | Gibson/NEB HiFi |
| pet28_twin_strep_syNOS_rev | catatgtcctccggattggaagtac | Gibson/NEB HiFi |
| syNOS_fwd | tccaatccggaggacatatgatgcttgtcaacgactctc | Gibson/NEB HiFi |
| syNOS_rev | gtggtggtggtggtgctcgagctacaagttggctagcc | Gibson/NEB HiFi |
| Hptg_fwd_PAYC_mcs1 | TAATGCTTAAGTCGAACAG | Gibson/NEB HiFi |
| Hptg_rev_PAYC_mcs1 | GGTATATCTCCTTATTAAAGTTAAAC | Gibson/NEB HiFi |
| Hptg_fwd | CTTTAATAAGGAGATATACCATGAAAGGACAAGAAACTCGTG | Gibson/NEB HiFi |
| Hptg_rev | TCTGTTCGACTTAAGCATTATCAGGAAACCAGCAGCTG | Gibson/NEB HiFi |
| gapa_fwd_pAYC_mcs2 | TAATTAACCTAGGCTGCTG | Gibson/NEB HiFi |
| gapa_rev_pAYC_mcs2 | ATGTATATCTCCTTCTTATACTTAAC | Gibson/NEB HiFi |
| gapa_fwd | TATAAGAAGGAGATATACATATGACTATCAAAGTAGGTATCAAC | Gibson/NEB HiFi |
| gapa_rev | GCAGCAGCCTAGGTTAATTATTATTTGGAGATGTGAGC | Gibson/NEB HiFi |
| pet28_tws_delc2_fwd | ctcgagcaccaccaccac | Gibson/NEB HiFi |
| pet28_tws_delc2rev | catatgtcctccggattggaag | Gibson/NEB HiFi |
| syNOSdelc2_178_1468_fwd | tccaatccggaggacatatgacctactcagaagccgtc | Gibson/NEB HiFi |
| syNOSdelc2_178_1468_fwd | tggtggtggtggtgctcgagctacaagttggctagccatttttc | Gibson/NEB HiFi |
| pet28_tws_delctt_fwd | tagctcgagcaccaccac | Gibson/NEB HiFi |
| pet28_tws_delctt_rev | catatgtcctccggattggaag | Gibson/NEB HiFi |
| syNOS_delctt1_1_1440_fwd | tccaatccggaggacatatgatgcttgtcaacgactctc | Gibson/NEB HiFi |
| syNOS_delctt1_1_1440_fwd | tggtggtggtgctcgagctagtagttgtctttctccacttg | Gibson/NEB HiFi |
| syNOS_K1412E_fwd | GCAAATTGCTgaaACAGAAGGAG | KLD |
| syNOS_K1412E_rev | ATAAATACCTCAAACACATTG | KLD |
| syNOS_T953A_fwd | GATCGGCAGCgcgGTCTATGAAC | KLD |
| syNOS_T953A_rev | CCTAGTACGGAATAGTTGAG | KLD |
| syNOS_K987A_fwd | CGACGAAATCgcgGGACAAGCCG | KLD |
| syNOS_K987A_rev | CCTTTATGTAGGGGAAC | KLD |

**Table S4**. Primers used in the study with the cloning approach (left).

| Domain | Couple | Potential (mV) | n | Calibrating dye (mV) |
|---|---|---|---|---|
| Globin (337-467) | $Fe^{3+}/Fe^{2+}$ | -250 ± 20 | 1 | safranin T (-289, 12.5μM) |
| Fld (856-1027) | ox/hq | -173 ± 3 | 2 | cresyl violet (-177.0, 60 μM ) |
| FNR (1027-1468) | ox/hq | -290 ± 13 | 2 | safranin T (-289, 12.5 μM) |

**Table S5.** Reduction potentials ($E^{0'}$) for $syNOS_{Glb}$, $syNOS_{Fld}$ and $syNOS_{FNR}$ vs. the Normal Hydrogen Electrode (NHE). For $syNOS_{Glb}$ the single electron reduction ($n_e$=1) from $Fe^{3+}$ to $Fe^{2+}$ was measured whereas for the flavin-containing domains a midpoint potential for the 2-electron reduction ($n_e$=2) from oxidized (ox) to hydroquinone (hq) was measured as no spectral signature for the semiquinone was apparent in the titrations. (n = 3 for all measurements)

#### Supplementary Figures

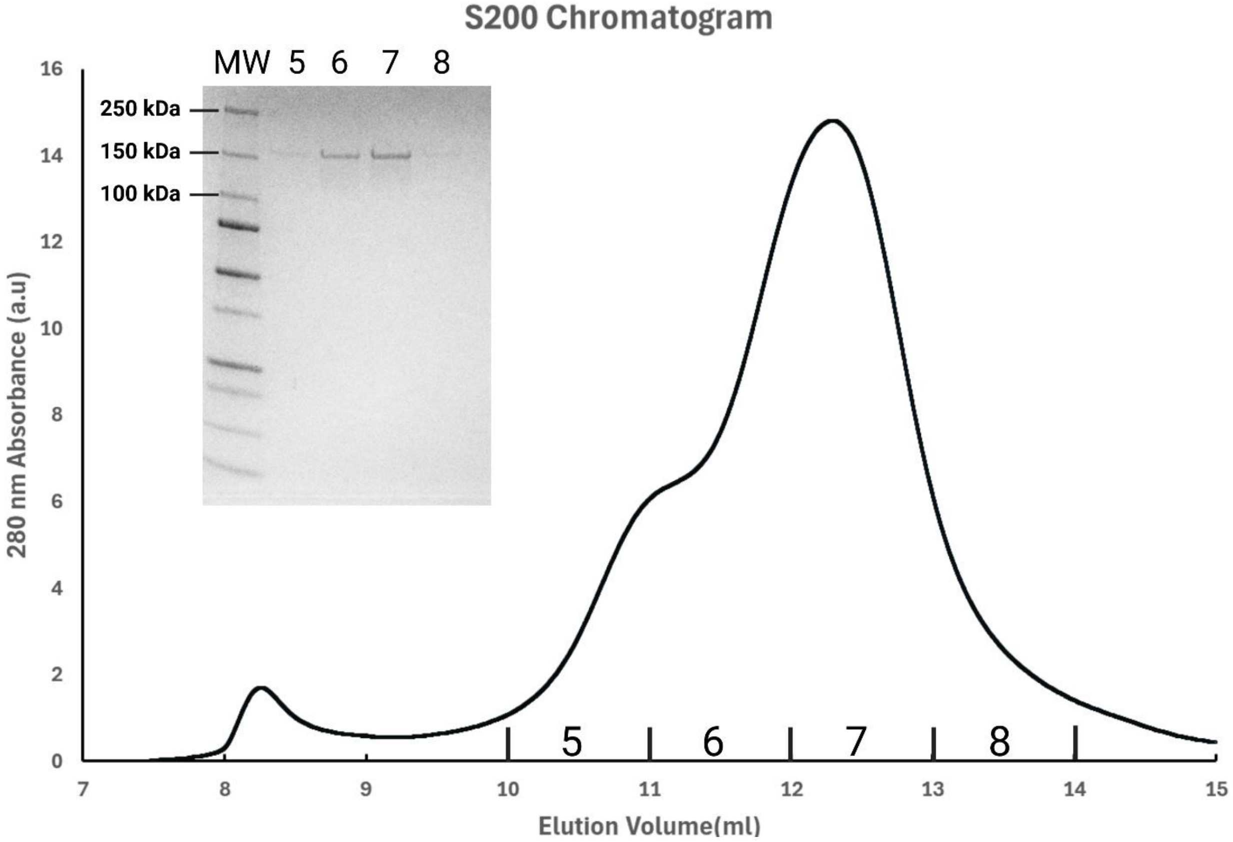


**Figure S1.** syNOS Purification. Representative chromatogram from a Superdex 200 increase 10/30 size exclusion chromatography column with an arrow denoting the syNOS dimer peak. The inset corresponds to a Coomassie blue stained 4-12% SDS-PAGE gel of fractions 5,6,7 and 8.

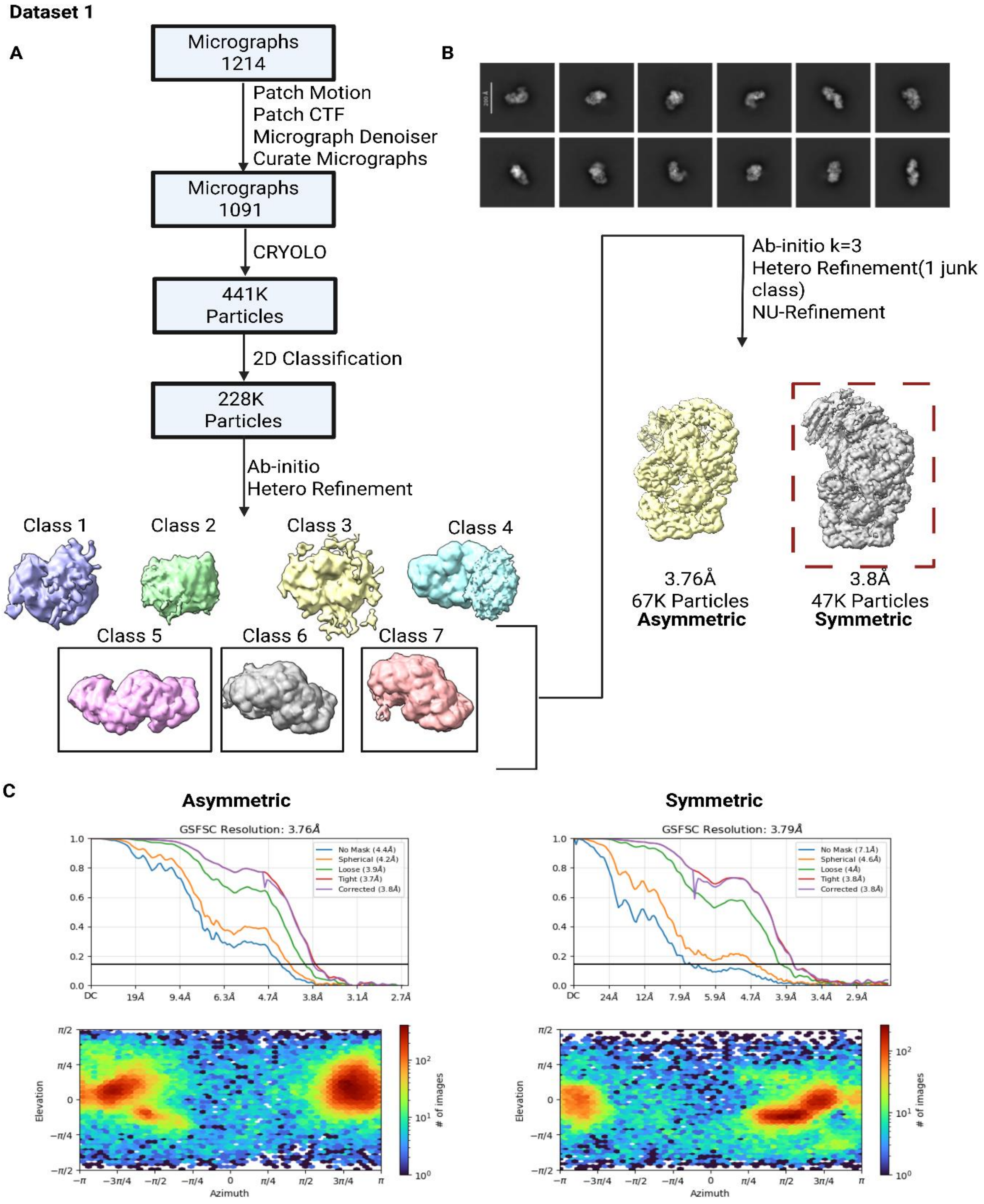


**Figure S2.** Dataset 1 processing scheme. (A) CryoEM processing workflow for dataset 1: syNOS ligand free. NOS-Sym class used for 3DVA and 3DFlex boxed (dashed-red). (B) Representative

2D Classes. CTF, contrast transfer function; NU, non-uniform refinement. (C) Overall Particle and Gold-standard Fourier Shell Correlation (GSFSC) curves for NOS-Asym and NOS-Sym.

**Dataset 2**

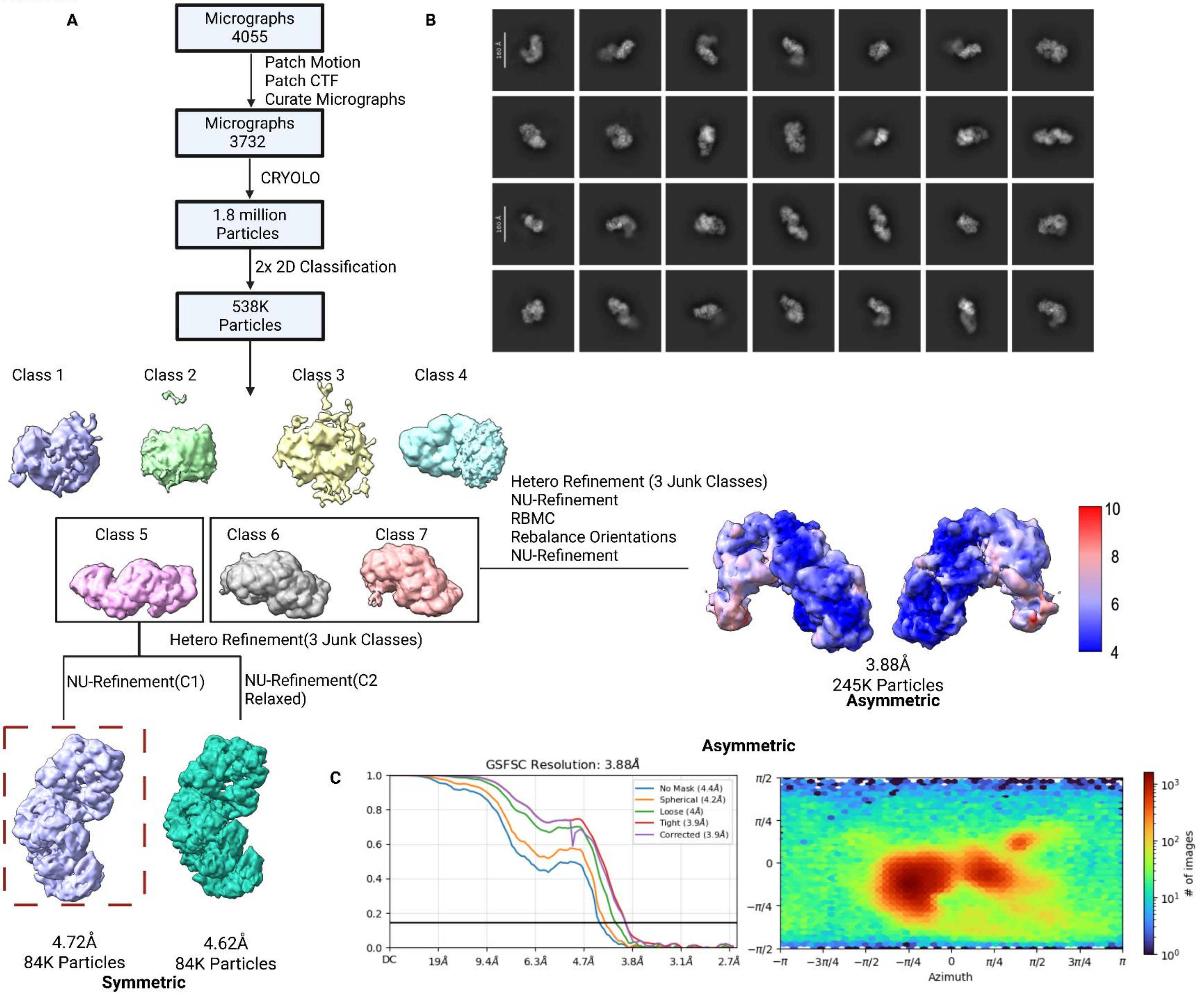

**Figure S3.** Dataset 2 processing scheme. (A) CryoEM processing workflow for dataset 2: syNOS ligand free. NOS-Sym class used for 3DVA and 3DFlex boxed (dashed-red). (B) Representative 2D Classes. (C) GSFSC curves and positional distribution of particles.

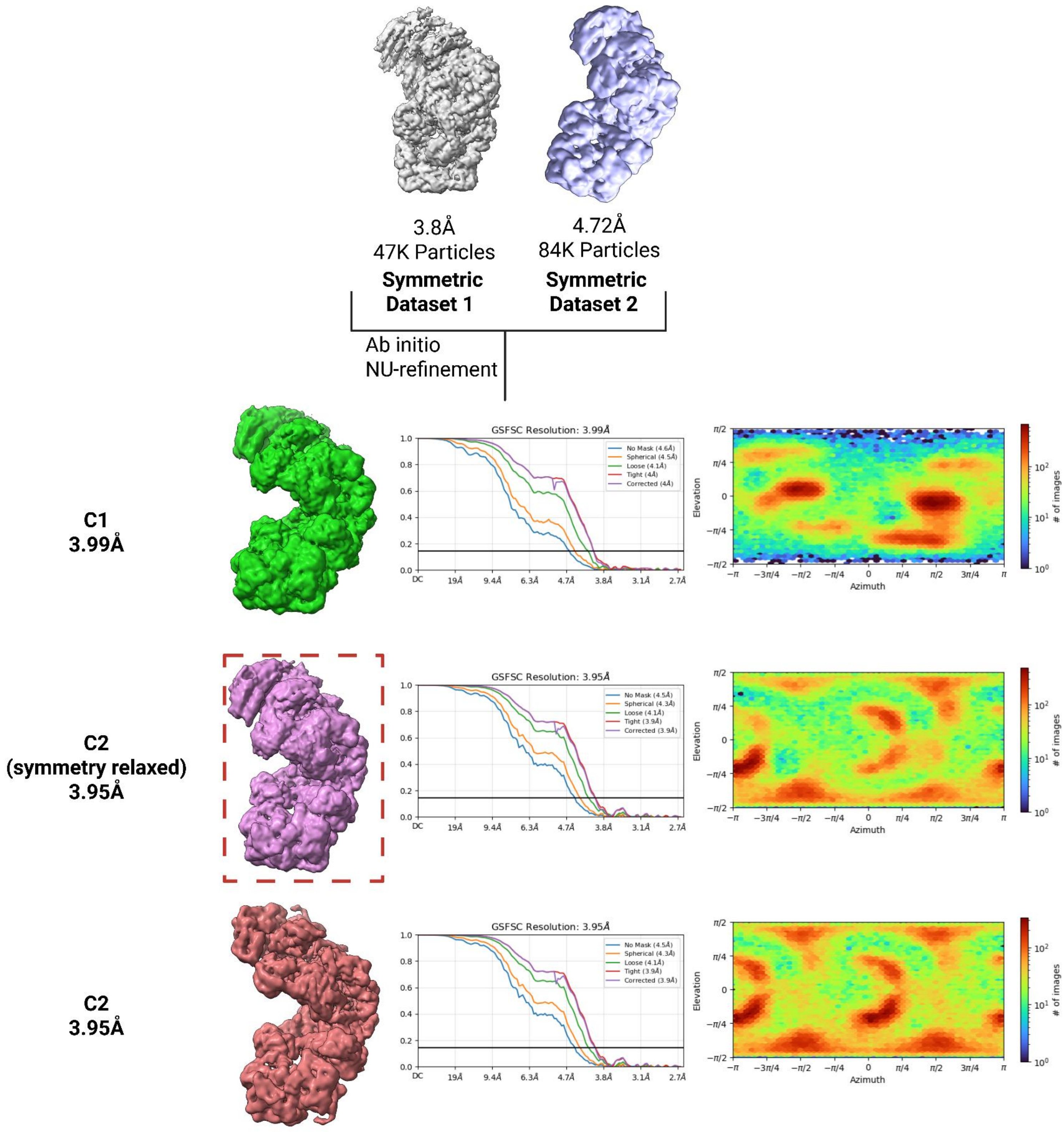


**Figure S4.** Merged data for NOS-Sym state, boxed class used for 3DVA and 3DFlex.

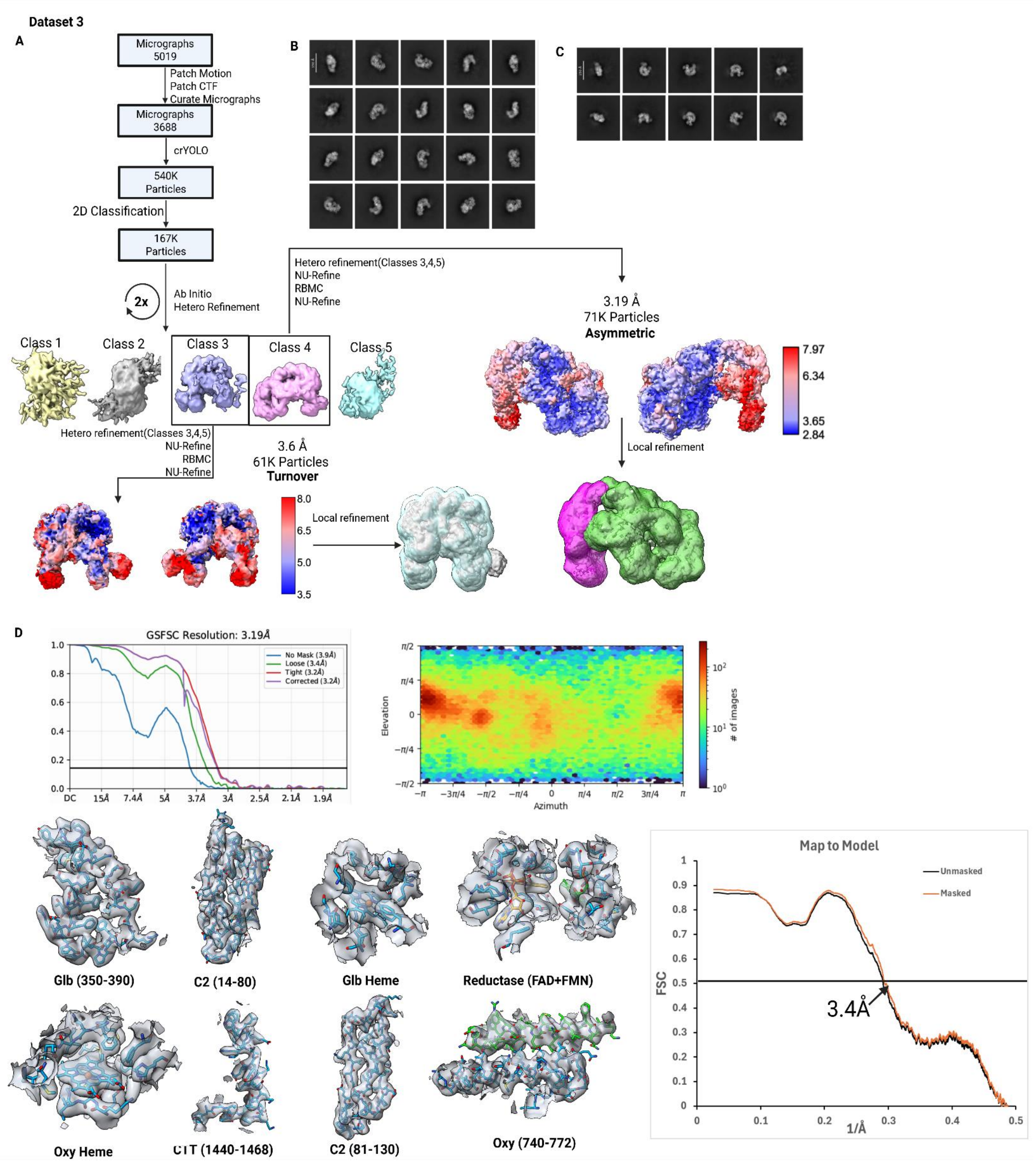


**Figure S5.** Dataset 3 processing scheme and validation. (A) CryoEM processing workflow for dataset 3: syNOS in the presence of $Ca^{2+}$/L-arginine (B) Representative 2D Classes for NOS-Asym state (C) Representative 2D classes for NOS turnover state (NOS-$TO_L$) (D) Asymmetric states GSFSC curves, orientation validation, model density fit and map to model Fourier Shell Correlation (FSC).

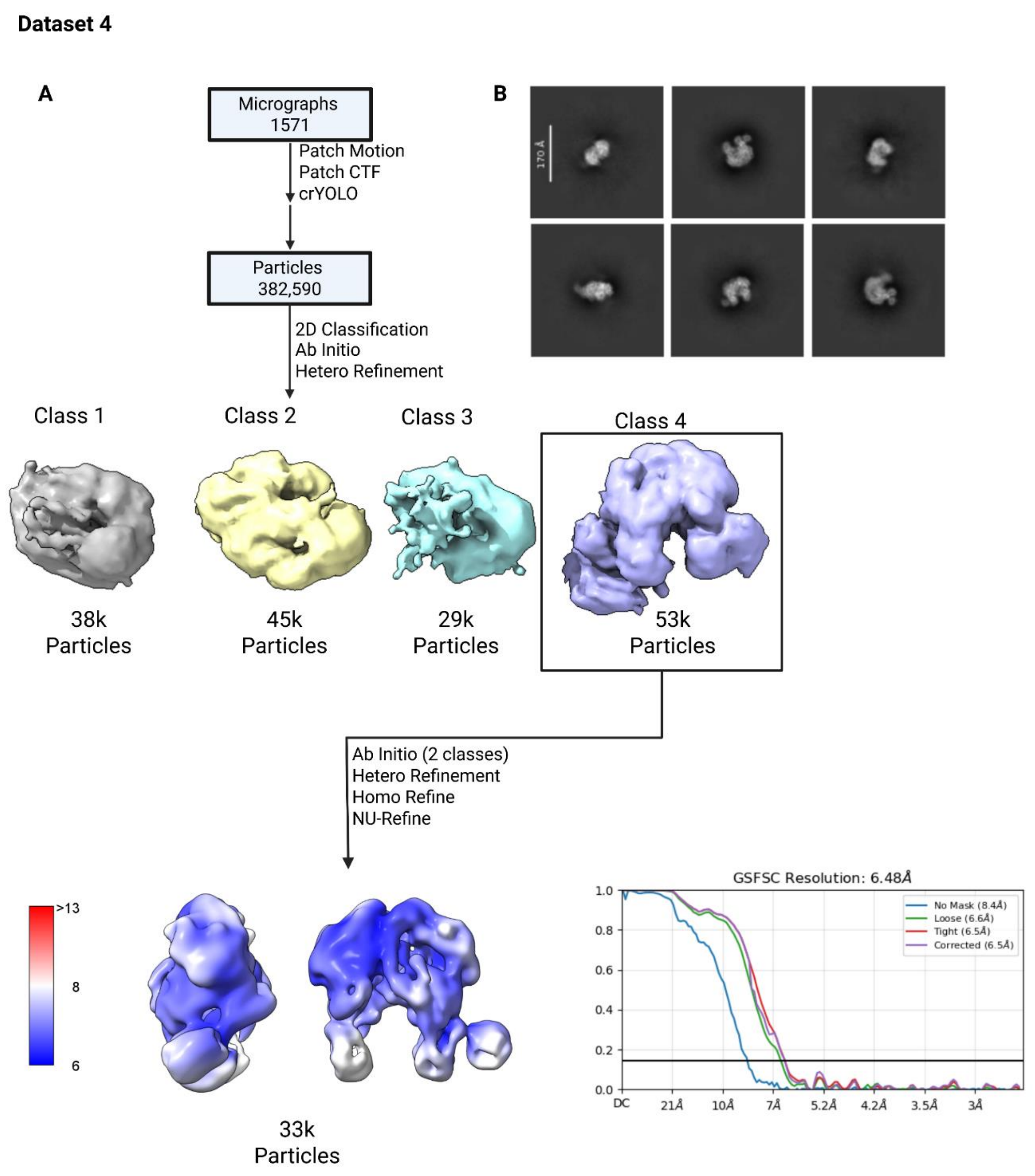


**Figure S6.** Dataset 4 processing scheme and validation. (A) CryoEM processing workflow for dataset 4, syNOS with NADPH/$Ca^{2+}$ including GSFSC curves. (B) Representative 2D classes for NOS-$TO_T$.

**Dataset 5**

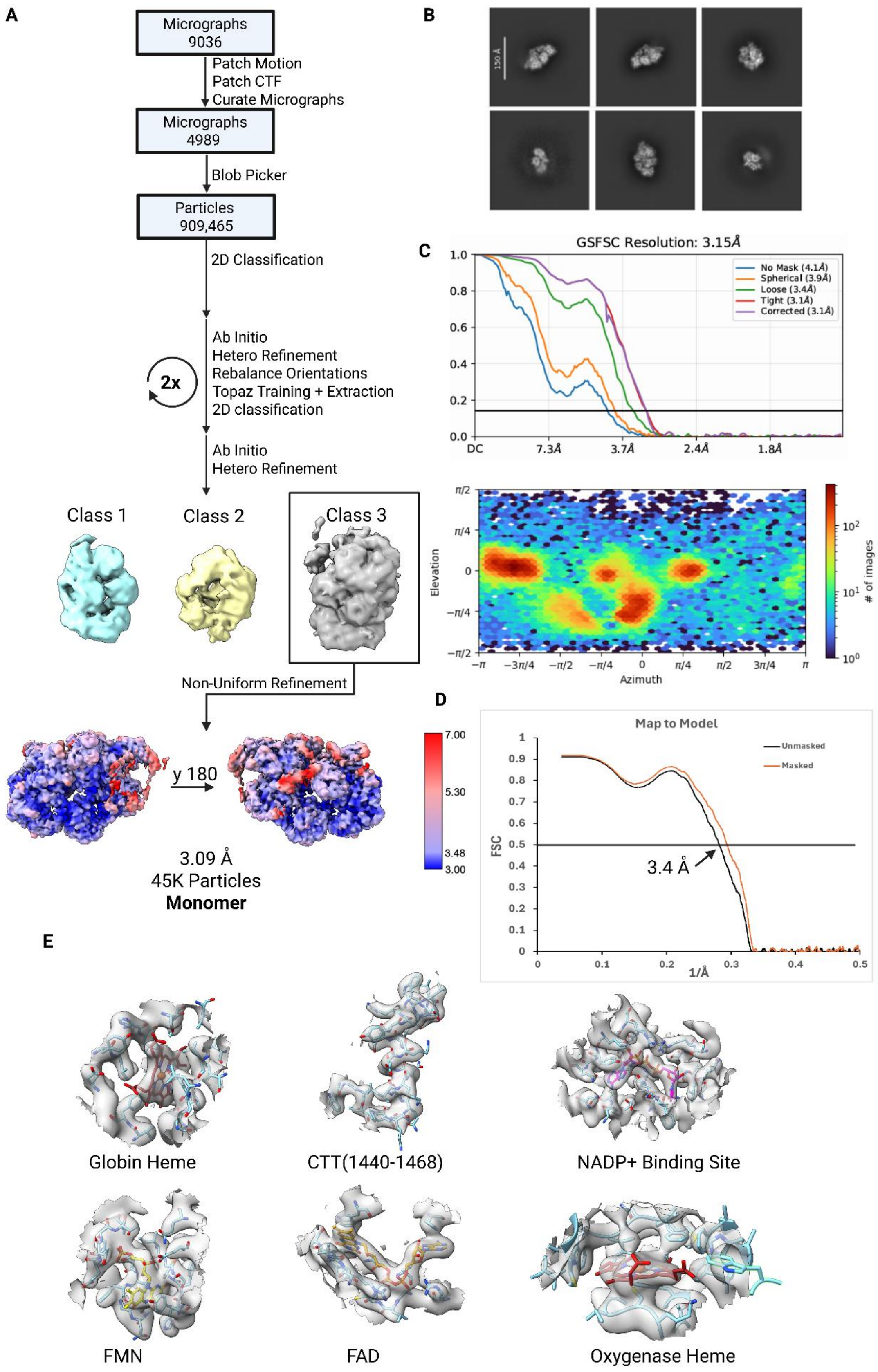

A
Micrographs 9036
Patch Motion
Patch CTF
Curate Micrographs
Micrographs 4989
Blob Picker
Particles 909,465
2D Classification
Ab Initio
Hetero Refinement
Rebalance Orientations
Topaz Training + Extraction
2D classification
2x
Ab Initio
Hetero Refinement
Class 1
Class 2
Class 3
Non-Uniform Refinement
y 180
3.09 Å
45K Particles
Monomer
7.00
5.30
3.48
3.00
B
150 Å
C
GSFSC Resolution: 3.15Å
No Mask (4.1Å)
Spherical (3.9Å)
Loose (3.4Å)
Tight (3.1Å)
Corrected (3.1Å)
DC
7.3Å
3.7Å
2.4Å
1.8Å
Elevation
Azimuth
### of images
D
Map to Model
Unmasked
Masked
FSC
1/Å
3.4 Å
E
Globin Heme
CTT(1440-1468)
NADP+ Binding Site
FMN
FAD
Oxygenase Heme

**Figure S7**. Dataset 5 processing scheme and validation. (A) CryoEM processing workflow for dataset 5: syNOS with L-arginine, NADPH/$Ca^{2+}$/NADPH (B) Representative 2D classes for NOS-Mon. (C) GSFSC curves, orientation validation and (D) Map to Model FSC. (E) Density fit of model in map.

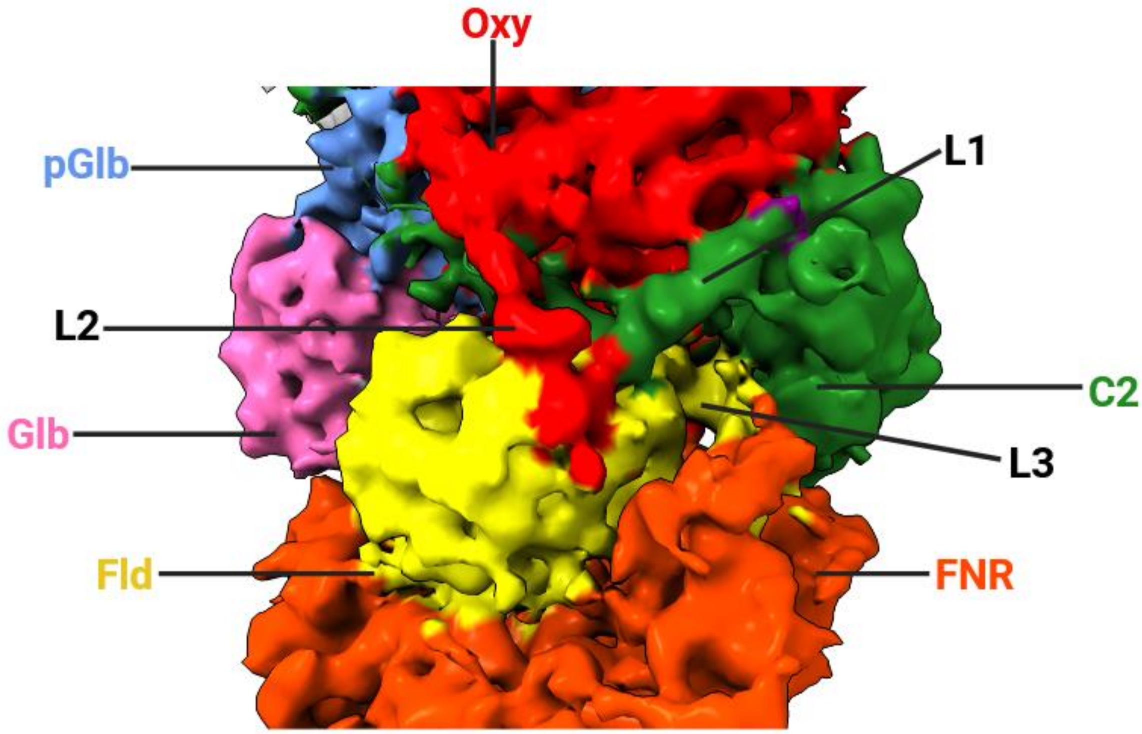


**Figure S8.** SyNOS linker intertwining. Intermediate resolution homodimer cryoEM density with all three linkers colored the same as their preceding domains. Note how L1 threads beneath L2 and on top of L3. Movement of the C2 domain (green) will release the Fld module for $NOS_{Oxy}$ reduction.

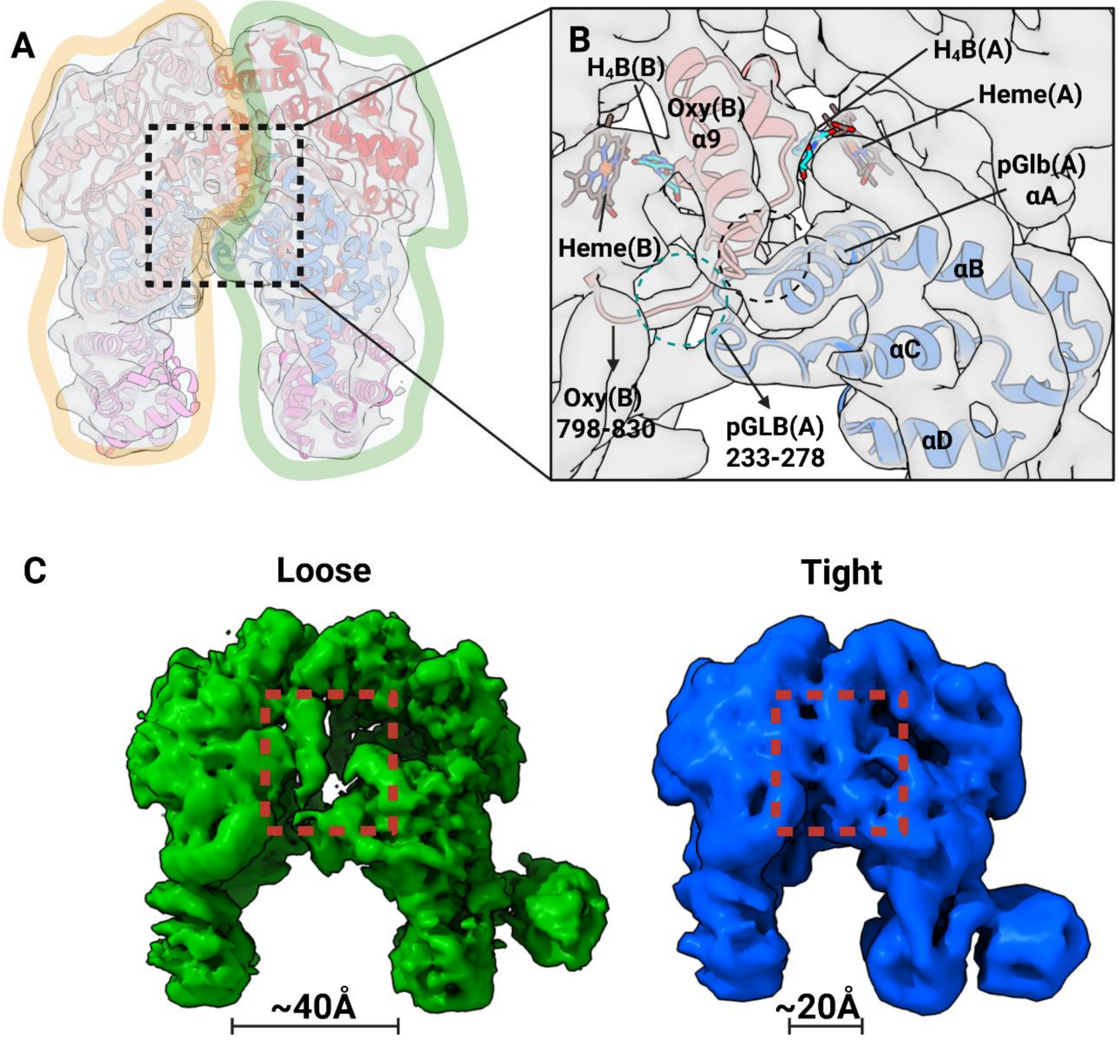


**Figure S9.** SyNOS tight and loose $NOS_{Oxy}$ states. (A) syNOS cryoEM density (dataset 4) with the $NOS_{Oxy}$ dimer interface outlined (orange and green) to show each subunit. (B) Close up of the dimeric interface showing the close association of $NOS_{Oxy}$ α9 to an intersubunit pGlb C/D loop. Although at lower resolution, strong electron density for the core the NOS-$TO_T$ interface includes the helical lariats and $BH_4$. (C) Loose (green) and tight (blue) $NOS_{Oxy}$ dimers differ by a large 20 Å breathing motion normal to the dimer interface that also affects the positioning of the C2 domain (lower right peripheral electron density).

A

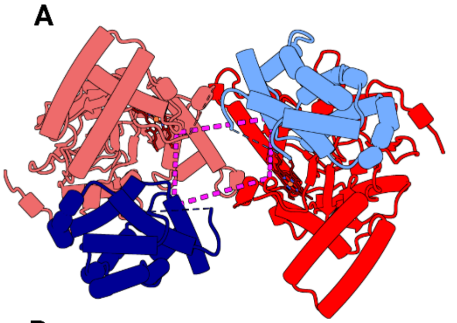

**Inactive Dimer Interface**

| Residue 1 | Residue 2 | Intra Cα- Cα | Inter Cα- Cα | XlinkX Score |
|---|---|---|---|---|
| 191(pGlb) | 538(Oxy) | **17** | **24** | 78 |

B

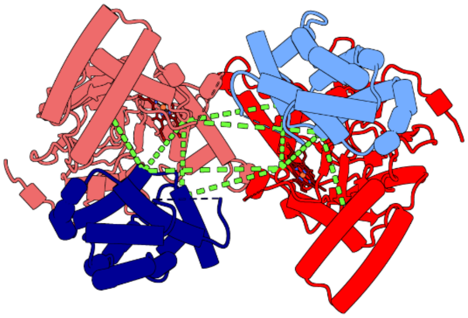

**Turnover Dimer Interface**

| Residue 1 | Residue 2 | Intra Cα- Cα | Inter Cα- Cα | XlinkX Score |
|---|---|---|---|---|
| 196(pGlb) | 538(Oxy) | **16** | **31** | 97 |
| 191(pGlb) | 538(Oxy) | **17** | **24** | 78 |
| 196(pGlb) | 481(Oxy) | **24** | **44** | 48 |

C

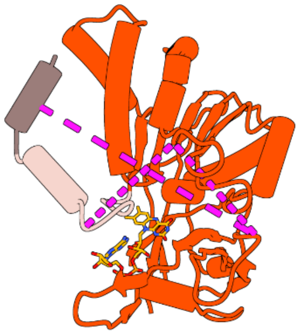

**Inactive CTT**

| Residue 1 | Residue 2 | Intra Cα- Cα | Inter Cα- Cα | XlinkX Score |
|---|---|---|---|---|
| 1366 | 1447(CTT) | **20** | 117 | 118 |
| 1053 | 1463(CTT) | **40** | 121 | 75 |
| 1366 | 1053(CTT) | **21** | 138 | 52 |

**Figure S10.** DSBU crosslinks plotted onto the dimer interface of syNOS Oxy + pGlb and the syNOS FNR + CTT. (A) syNOS inactive dimer interface, subunit 1 Oxy (red) + pGlb (blue) and subunit 2 Oxy (bright red) + pGlb (navy), with magenta crosslinks corresponding to table (left). (B) syNOS inactive dimer interface, protomer 1 Oxy (red) + pGlb (blue) and subunit 2 Oxy (bright red) + pGlb (navy), with green crosslinks corresponding to table (left). (C) syNOS inactive CTT with magenta crosslinks corresponding to table (left). Crosslinks within range based on the structures are underlined and bolded.

A

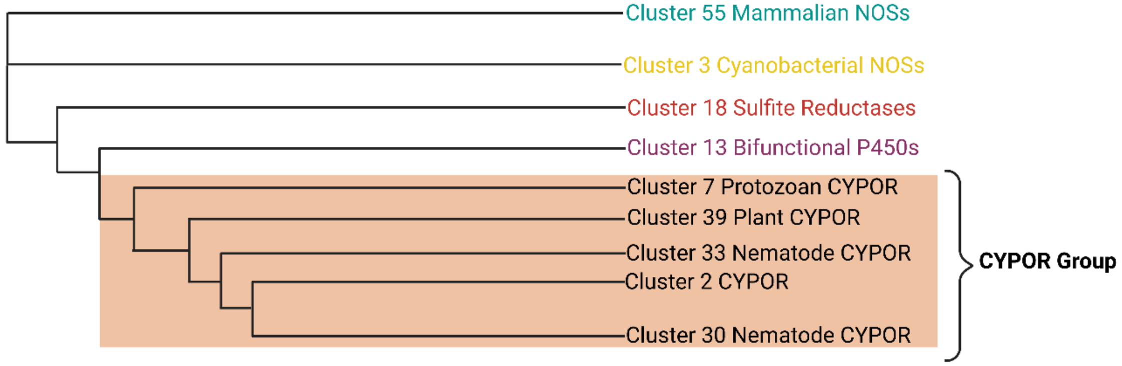


B

| Cluster ID | Average Sequance Length | Standard Deviation of Sequence Length | Sequence Count |
|---|---|---|---|
| 2 | 678 | 63 | 2521 |
| 39 | 702 | 49 | 1588 |
| 55 | 1117 | 103 | 278 |
| 3 | 1479 | 118 | 94 |
| 13 | 976 | 174 | 63 |
| 58 | 700 | 17 | 49 |
| 30 | 1126 | 283 | 47 |
| 7 | 638 | 35 | 45 |
| 18 | 601 | 74 | 44 |
| 33 | 652 | 36 | 44 |

**Figure S11.** Phylogenetic sequence clustering in the context of syNOS$_{Red}$(residues 853-1468). Representative clusters were subjected to PhyML with clusters above 40 sequences represented in the tree. (B) Cluster composition by sequence length and count.

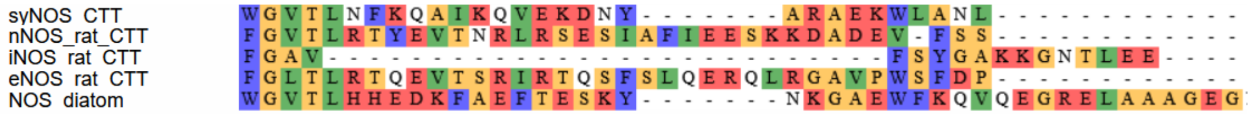


**Figure S12.** C-terminal tail (CTT) sequence alignment of syNOS, mNOSs and diatom NOS.

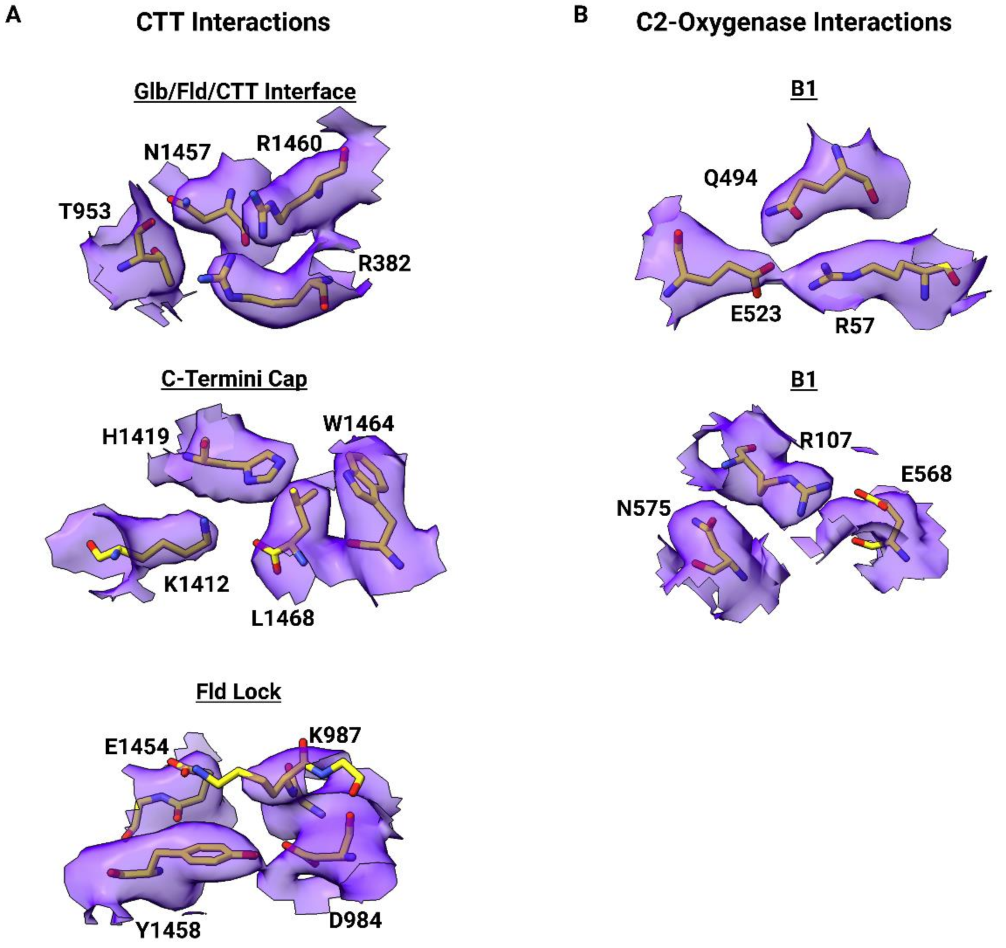


**Figure S13**. Cryo-EM density validation of key interactions. (A) Density fit of syNOS-Asym (9Q15), key interactions shown in Fig 3 C-E (B) Density fit of syNOS-Asym (9Q15), key interactions shown in Fig 4. B1-B2.

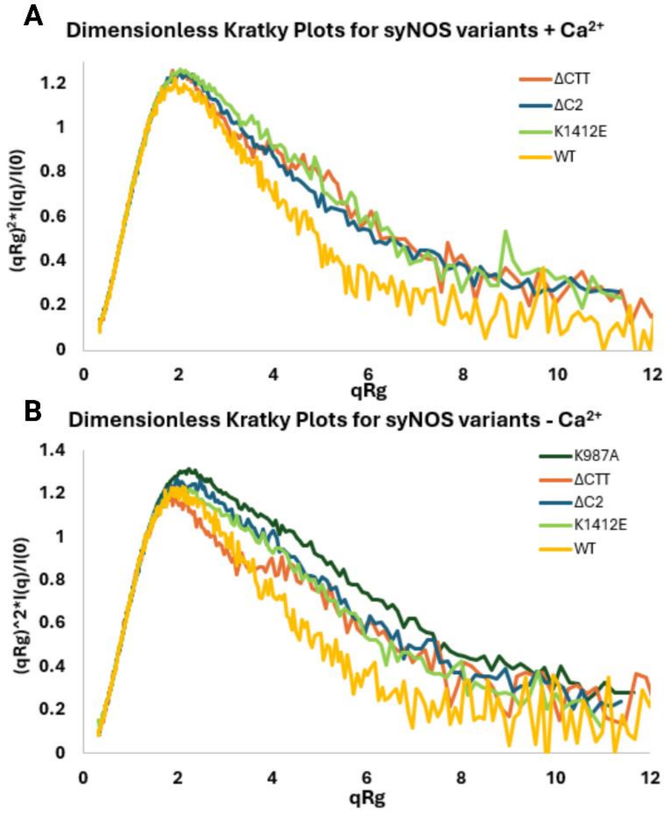


C

| Sample | Rg [Guinier] (Å) | MW(Vc) | MW(Vp) |
|---|---|---|---|
| Predicted WT Dimer | 55.59 | - | - |
| Predicted WT Monomer | 38.81 | - | - |
| WT $Ca^{2+}$ | 52.83 ± 0.44 | 277.8 | 316.8 |
| ΔC2 $Ca^{2+}$ | 56.89 ± 0.29 | 317.9 | 366.9 |
| ΔCTT $Ca^{2+}$ | 62.71 ± 0.37 | 237.9 | 288.6 |
| K1412E $Ca^{2+}$ | 57.16 ± 0.3 | 222 | 284 |
| WT EGTA | 52.83 ± 0.44 | 234.7 | 280.2 |
| ΔC2 EGTA | 57.34 ± 0.3 | 222.7 | 272.2 |
| ΔCTT EGTA | 61.18 ± 0.55 | 313.3 | 360 |
| K1412E EGTA | 55.28 ± 0.24 | 221.4 | 258.2 |
| K987A EGTA | 58.79 ± 0.37 | 213.6 | 266.7 |

**Figure S14.** SEC-SAXS of syNOS and variants. (A) Dimensionless Kratky plots, binned by 4 qRg values, for samples in the presence and absence of $Ca^{2+}$. (B) Table of predicted and experientially characterized radius of gyrations (Rgs) and molecular weights calculated either by volumes of correlation ($V_C$) or Porod volumes ($V_P$). (Samples were measured once)

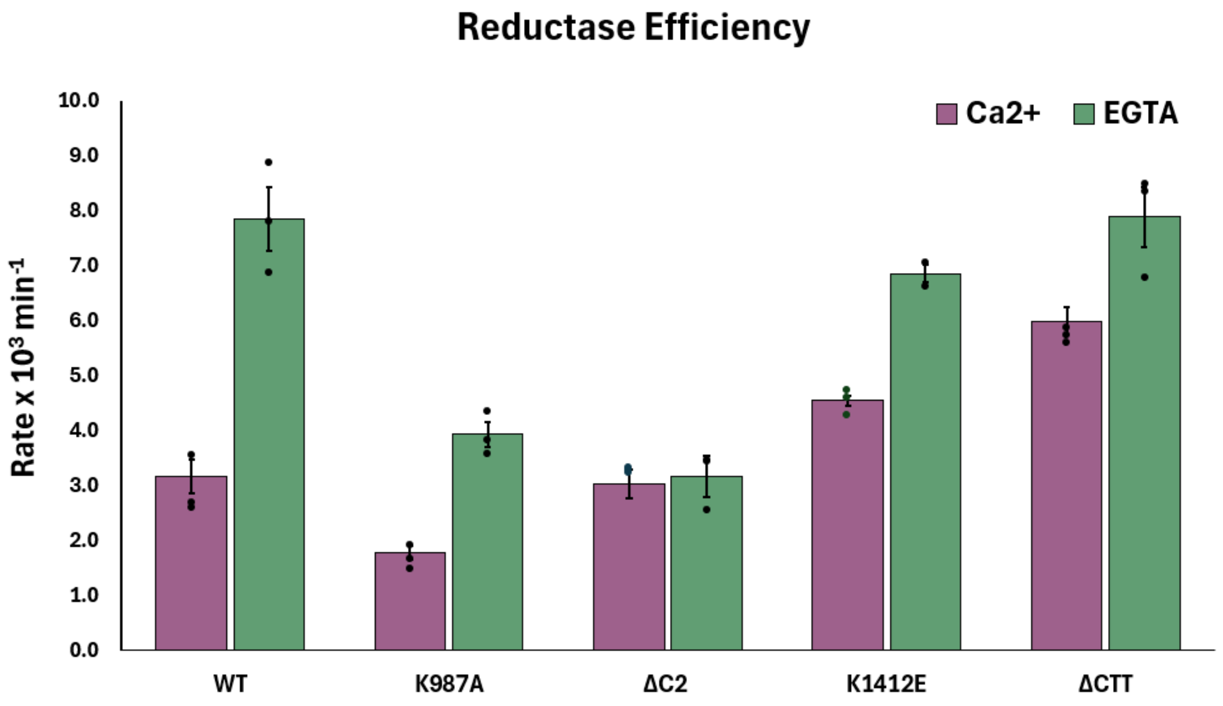


**Figure S15.** Steady state cytochrome c reductase activities of WT syNOS and variants expressed as turnover numbers (Cc reduced $(NOS\ subunit)^{-1}\ min^{-1}$) with standard errors shown as error bars (see Methods). Statistical comparisons between groups were performed with a 2-way ANOVA with post hoc analysis using Tukey's multiple comparisons test. Significant differences are indicated as follows: ns (not significant), * ($p \leq 0.05$), ** ($p \leq 0.01$), *** ($p \leq 0.001$), **** ($p \leq 0.0001$)..Ca:WT vs. Ca:K987A (ns); Ca:WT vs. Ca:ΔC2 (ns); Ca:WT vs. Ca:K1412E (ns); Ca:WT vs. Ca:ΔCTT (ns); Ca:WT vs. EGTA:WT (****) ; Ca:WT vs. EGTA:K987A (ns); Ca:WT vs. EGTA:ΔC2 (ns); Ca:WT vs. EGTA:K1412E (****); Ca:WT vs. EGTA:ΔCTT (****) ;Ca:K987A vs. Ca:ΔC2 (ns); Ca:K987A vs. Ca:K1412E (***); Ca:K987A vs. Ca:ΔCTT (****); Ca:K987A vs. EGTA:WT (**); Ca:K987A vs. EGTA:K987A (**); Ca:K987A vs. EGTA:ΔC2 (ns); Ca:K987A vs. EGTA:K1412E (****); Ca:K987A vs. EGTA:ΔCTT (****); Ca:ΔC2 vs. Ca:K1412E (ns); Ca:ΔC2 vs. Ca:ΔCTT (**); Ca:ΔC2 vs. EGTA:WT (****); Ca:ΔC2 vs. EGTA:K987A (ns); Ca:ΔC2 vs. EGTA:ΔC2 (ns); Ca:ΔC2 vs. EGTA:K1412E (****); Ca:ΔC2 vs. EGTA:ΔCTT (****);Ca:K1412E vs. Ca:ΔCTT (ns); Ca:K1412E vs. EGTA:WT (****); Ca:K1412E vs. EGTA:K987A (ns); Ca:K1412E vs. EGTA:ΔC2 (ns); Ca:K1412E vs. EGTA:K1412E (**); Ca:K1412E vs. EGTA:ΔCTT (****); Ca:ΔCTT vs. EGTA:WT (**); Ca:ΔCTT vs. EGTA:K987A (*); Ca:ΔCTT vs. EGTA:ΔC2 (***); Ca:ΔCTT vs. EGTA:K1412E (ns); Ca:ΔCTT vs.

EGTA:ΔCTT (**); EGTA:WT vs. EGTA:K987A (****); EGTA:WT vs. EGTA:ΔC2 (****); EGTA:WT vs. EGTA:K1412E (ns); EGTA:WT vs. EGTA:ΔCTT (ns); EGTA:K987A vs. EGTA:ΔC2 (ns); EGTA:K987A vs. EGTA:K1412E (***); EGTA:K987A vs. EGTA:ΔCTT (***); EGTA:ΔC2 vs. EGTA:K1412E (****); EGTA:ΔC2 vs. EGTA:ΔCTT (****); EGTA:K1412E vs. EGTA:ΔCTT (ns). (n=3, except for K1412E +EGTA; n = 2).

**A**

| Reductase | $\lambda_{obs}$ | $A_1$ (%) | $k_{obs1}$ ($s^{-1}$) | $A_2$ (%) | $k_{obs2}$ ($s^{-1}$) | $A_3$ (%) | $K_{obs3}$ ($s^{-1}$) | |
|---|---|---|---|---|---|---|---|---|
| $syNOS_{Red}$(-$Ca^{2+}$) | 450 | 89 | 178 ( ± 0.78) | 11 | 4.12 ( ± 0.03) | - | - | This study |
| $syNOS_{FL}$(-$Ca^{2+}$) | 450 | 63 | 2.38 ( ± 0.03) | 37 | 0.004 ( ± 0.001) | - | - | This study |
| $nNOS_{Red}$(-CAM) | 458 | 28 | 57 ( ± 1) | 52 | 5.2 ( ± 0.1) | 20 | 570 ( ± 41) | Craig et al.[17] |
| $nNOS_{Red}$(+CAM) | 458 | 44 | 608 ( ± 66) | 56 | 86 ( ± 2) | - | - | Craig et al.[17] |
| CYPOR | 450 | 90 | 42 ( ± 4) | 10 | 3.8 ( ± 0.5) | - | - | Hamdane et al.[68] |

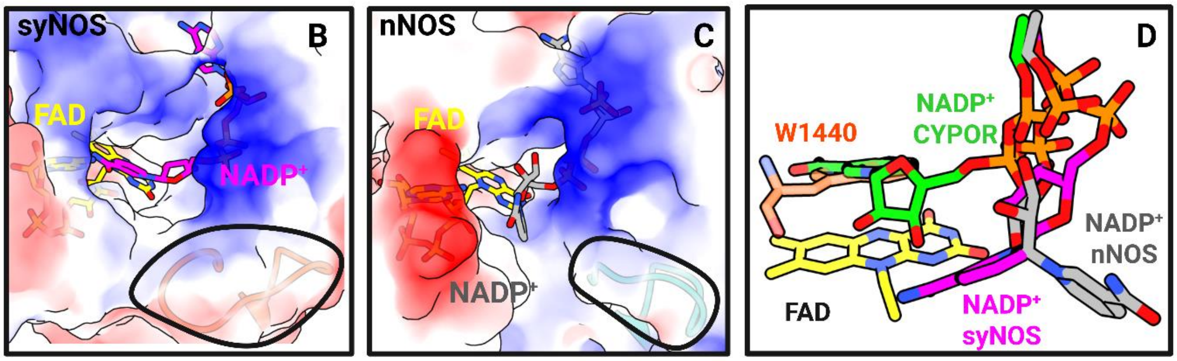


**Figure S16.** Flavin reduction by NADPH. (A) Table of flavin reduction rate constants for syNOS, nNOS and CYPOR(n = 4). (B) Surface representation and model of $NADP^+$ (magenta) bound in the pocket of syNOS-Mon with loop 1050-1062 (orange-red and circled in black) in comparison to (C) nNOS bound to $NADP^+$ (grey) with corresponding loop 1002-1012 (aqua) circled Iin black. (D) Overlay of $NADP^+$ bound conformations of syNOS, nNOS and CYPOR W677X (1-676) (PDB: 1JA0) (green) over a consensus position of FNR-bound FAD (yellow). syNOS CTT residue 1440 blocks access of the nicotinamide ring to the isoalloxazine ring.

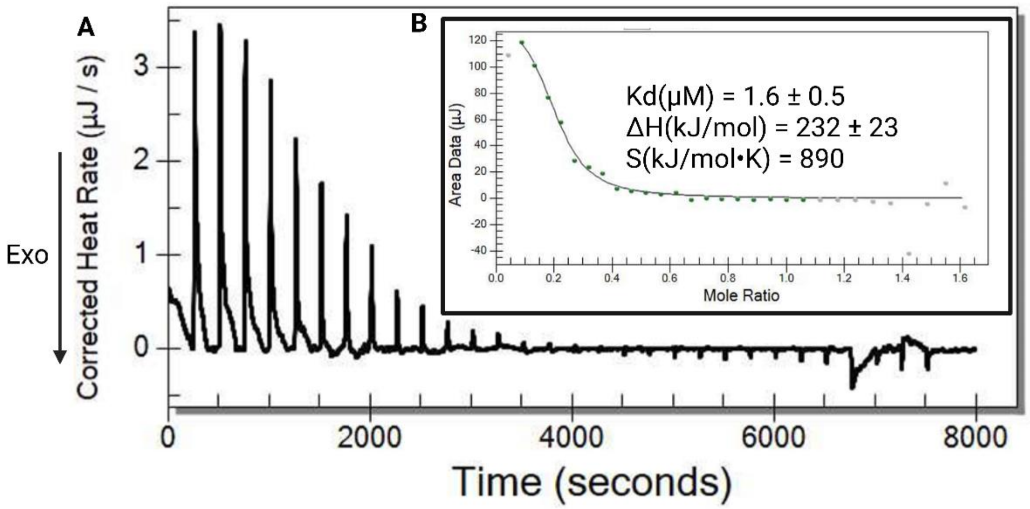


**Figure S17.** Isothermal calorimetric titration of $Ca^{2+}$ with syNOS$_{C2}$ (residues 1-138). (A) Endothermic, baseline corrected isotherm of $Ca^{2+}$ titrated into syNOS$_{C2}$. (B) Binding model fit with the accompanying thermodynamic parameters. (Samples were measured once)

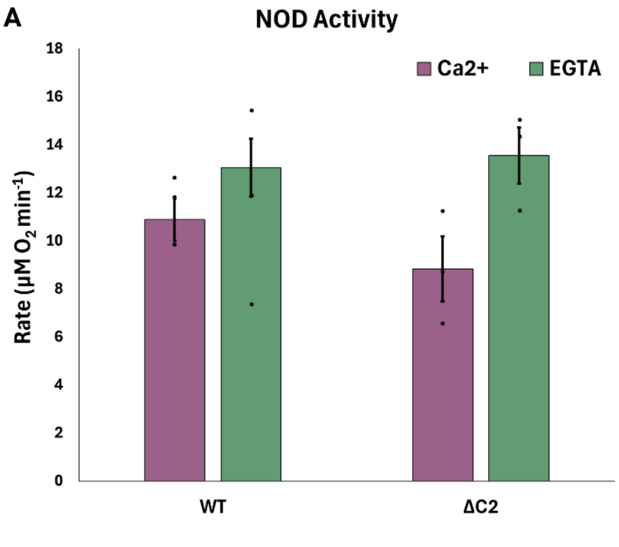


B

| Variant | Condition | Rate constant ($s^{-1}$) |
|---|---|---|
| WT | Ca | 0.33 ( ± 0.02 ) |
| ΔC2 | | 0.54 ( ± 0.05 ) |
| WT | EGTA | 0.22 ( ± 0.02 ) |
| ΔC2 | | 0.35 ( ± 0.03 ) |
| Glb +FNR | | 0.25 ( ± 0.03 ) |

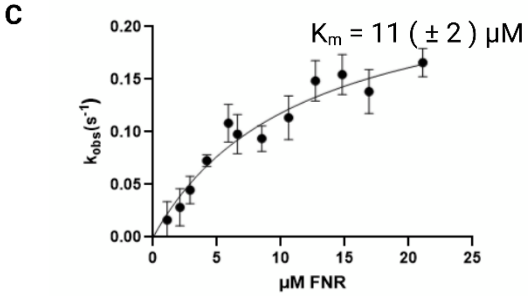


**Figure S18.** SyNOS globin-based NOD activity and heme reduction rates. (A) NOD activity was measured by O2 consumption in the presence of the NO-donor NOC-7 and NADPH for WT and syNOS$_{\Delta C2}$ (n=3, except for syNOS$_{\Delta C2}$ + $Ca^{2+}$ ; n = 2). (B) Heme reduction rate constants of syNOS$_{Glb}$ in the presence or absence of $Ca^{2+}$ for WT or syNOS$_{\Delta C2}$ and the truncated syNOS$_{Glb}$/syNOS$_{FNR}$ couple in the absence of $Ca^{2+}$, with standard error represented. Statistical comparisons between groups were performed with a 2-way ANOVA with post hoc analysis using Tukey's multiple comparisons test, all comparisons were non-significant. (C) Michelis-Menten plot of truncated syNOS$_{Glb}$ reduced by varied concentrations of truncated syNOS$_{FNR}$ until saturation, with standard deviation represented. (Each data point reflects at least n = 3.)

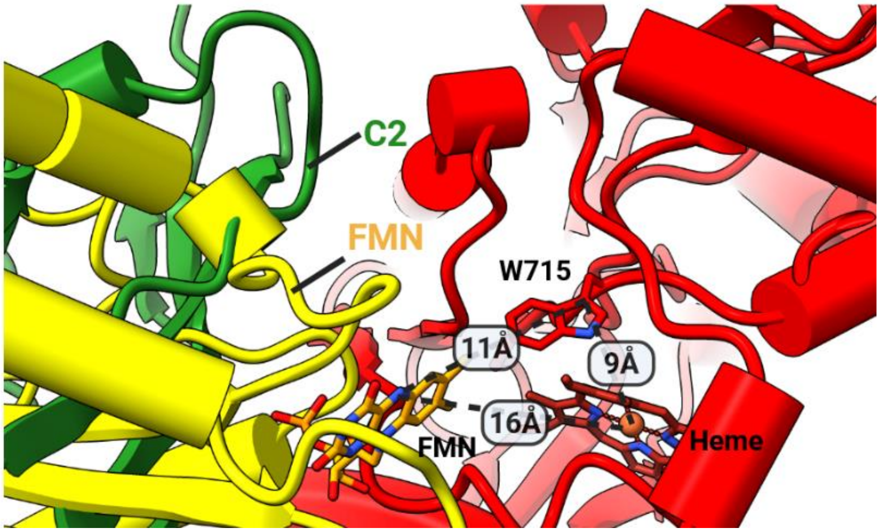


**Figure S19.** Output state predicted by AlphaFold3 with FMN-to-Trp715-to-Heme distances shown. Overlay of the C2 in the locked state (green) with the output Fld conformation (yellow) illustrates that the Fld is sterically blocked by the C2 domain in the NOS-Asym state. The AF3 prediction was made with a minimal unit of two truncated syNOS chains: 1) residues 473-863 (Oxy) and 2) residues 473-1003 (Oxy + Fld). Including additional domains did not predict interaction between syNOS$_{Fld}$ and syNOS$_{Oxy}$.
